# Eukaryotic 45S ribosomal RNA spacers encode an ancient layer of antiviral response

**DOI:** 10.64898/2026.11.17.767335

**Authors:** Katarzyna Michalak, Ariana Faraji, Lin Kang, Sabeen Wazir, Anne Remorca, Erin Vasquez, Trevor Morris, Francheska Brzazgon, Kayla Griffith, Hannah Khairandish, Rose Pwint, Stephen DiGiuseppe, Pawel Michalak

## Abstract

Eukaryotic 45S ribosomal RNA genes produce internal transcribed spacers (ITSs) that are excised and discarded during ribosome biogenesis, with no established function beyond rRNA processing. Here, we uncover an unexpected antiviral role for these seemingly expendable sequences. Comparative genomic analyses reveal extensive, evolutionarily widespread sequence similarities between eukaryotic ITSs and DNA viruses, particularly between vertebrate ITSs and herpesviruses. Across herpesvirus infections, ITS transcripts undergo striking, virus-specific remodeling, largely uncoupled from mature ribosomal RNA production. During herpes simplex virus 1 (HSV-1) infection, hierarchical Bayesian comparison of competing dynamical models favors active host ITS deployment over passive viral remodeling by odds of 174,000:1.

Experimental depletion of ITS2 consistently increases infectious HSV-1 production by approximately 70%, whereas supplementation with synthetic ITS2 RNA generally suppresses viral replication. These findings reveal a previously unrecognized antiviral activity within the 45S rRNA system, challenging the view of ITSs as disposable byproducts of ribosome biogenesis.

## Introduction

The effects of viral infections on host cell functions are multifaceted. Intriguingly, amid the diverse strategies employed by different viral families and their consequences for the host, a shared characteristic emerges: their interaction with the nucleolus, a well-established non-membrane nuclear compartment. Eukaryotic nucleolus encapsulating rRNA genes forms a highly autonomous and dynamic subnuclear structure with fundamental roles in ribosome subunit biogenesis, which is central to all cell metabolism and growth. Nucleolar autonomy can be likened to a quasi-symbiotic mini-life, possessing its own miniature genome, unique origin of replication and replication fork block, a cluster of highly specialized proteins, including the Pol I transcription machinery specific to 45S rRNA, as well as spectacular epigenetic memory (Michalak et al., 2015; Michalak, 2014). rRNA genes are clustered within ancient parts of the genome formed at the time of the Last Universal Common Ancestor (LUCA) (Fox, 2010).

Viruses are obligate intracellular parasites that use host cell machinery for their replication. Typically, DNA viruses replicate within the nucleus, whereas RNA viruses replicate in the cytoplasm, but some viruses with minus RNA strand genomes also replicate their genomic RNA in the nucleus and closely interact with nucleoli (Iarovaia et al., 2021). Viruses interface with nucleoli to hijack host-cell functions, modulate pre-rRNA gene transcription and maturation, and recruit nucleolar proteins to facilitate virus replication (Hiscox, 2007; Jaafar and Kieft, 2019).

The architecture and proteome of nucleoli undergo a dramatic reorganization in response to viral infection, in part under virus control, but in part with the host still in charge (Li, 2019; Mazumder et al., 2014). Alongside the myriad viral components shuttling to and from nucleoli, many endogenous nucleolar proteins undergo relocation from the nucleolus, while other non-nucleolar cellular proteins are trafficked into the nucleolus to perform diverse functions (Salvetti and Greco, 2014). For example, during infection with the herpes simplex virus (HSV), an alpha-herpesvirus from the *Herpesviridae* family, consisting of DNA viruses with relatively large genomes, three major nucleolar proteins, nucleolin, nucleophosmin, and fibrillarin, along with RPA 194, translocate to the Viral Replication Compartments (VRCs). Within the VRCs, they co-localize with the viral DNA-binding protein ICP8, contributing to the replication, transcription, and assembly of viral particles (Iarovaia et al., 2021). During infection with human cytomegalovirus (HCMV), a betaherpesvirus, interaction between nucleolin and the viral DNA polymerase component UL44 is essential for efficient DNA replication and the expression of viral proteins (Strang, 2015).

Due to the hijacking of the nucleoli by viruses, transcription of ribosomal genes is typically enhanced, leading to an increase in the number of ribosomes (Hiscox, 2007; Iarovaia et al., 2021). However, in some cases, the transcription of ribosomal genes is unaffected or even suppressed upon viral infection. The biological significance of the latter phenomenon, which is observed in infections with viruses like poliovirus, rhinovirus, influenza H3N2, Hendra, and Nipah (Iarovaia et al., 2021), is challenging to interpret, unless one presumes that the cell retains control and alterations of rRNA expression may serve the host’s interests. The dramatic redundancy of eukaryotic rRNA genes, with typically hundreds of rRNA cistrons per genome, could be a hallmark of ongoing genomic conflict between ribosomal DNA and viruses. There is little evidence that rRNA gene redundancy is essential for normal cellular metabolism, as a significant proportion of rRNA genes remains in a heterochromatinized state and is never transcribed (Grummt, 2007; Santoro, 2005). A mutant chicken genotype possessing approximately 160 out of the typical complement of 290 45S rRNA genes (accounting for 56%) still sustains normal development (Delany et al., 1995), while yeast species naturally exhibit a four-fold variation in rRNA gene copy number (Maleszka and Clark-Walker, 1993). Sustained protein synthesis has been observed in germline stem cell differentiation in *Drosophila* despite reduced nucleolar volume and decreased rDNA transcription (Neumuller et al., 2008; Sanchez et al., 2016).

The 45S rRNA genes are among the largest repetitive elements in eukaryotic genomes, consisting of highly redundant tandem clusters of rRNA genes. Within each repeat unit, or rRNA cistron, putatively nonfunctional internal transcribed spacers 1 (ITS1) and 2 (ITS2) are imbedded between genes coding for 18S, 5.8S and 28S rRNA (**Figure 1**) (Caudy and Pikaard, 2002). ITS1 and ITS2 sequences are transcribed into a 45S precursor molecule but later this precursor is cleaved at specific sites and the spacers are removed. While the rRNA genes are highly conserved across the tree of life, ITS1 and ITS2 sequences are strikingly divergent between species, to the extent that they provide useful DNA “barcoding” markers for the identification and taxonomy of eukaryotes, including closely related sibling species (Chase and Fay, 2009). In addition to sequence divergence, ITS primary sequences also vary greatly in their length (Hlinka et al., 2002). This pattern of abrupt divergence implies a powerful molecular drive generating genetic novelties, in addition to the accumulation of mutations via natural selection and genetic drift (Dover, 1982). In stark contrast to its sequence, the predicted secondary structure of ITS2 tends to be conserved across a variety of eukaryotic taxa, typically featuring four to five standard helix loops (Coleman, 2013; Hlinka et al., 2002; Schultz et al., 2005). The striking divergence of ITS sequences among species, despite their relative homogeneity within species, is characteristic of concerted evolution, whereby unequal crossing-over, gene conversion, and other mechanisms of molecular drive can rapidly spread sequence variants through multicopy gene families (Dover, 1982; Eickbush and Eickbush, 2007; Gerbi, 1986). Such processes provide a powerful mechanism for generating and propagating evolutionary novelty, but they do not necessarily explain the selective forces shaping that dramatic variation. One particularly compelling candidate is the persistent evolutionary conflict between hosts and their genetic parasites. More than half a century ago, Van Valen’s Red Queen hypothesis proposed that organisms must continually evolve merely to maintain their relative fitness in an ever-changing biotic environment (Van Valen, 1973). Host–parasite coevolution subsequently emerged as a powerful theoretical framework for understanding the maintenance of genetic diversity and the evolution of biological defenses (Hamilton et al., 1990). Viruses, with their extraordinary diversity, rapid evolution, and intimate dependence on host cellular machinery, represent particularly potent agents of such antagonistic coevolution (Daugherty and Malik, 2012; Koonin and Wolf, 2015).

**Figure 1.**
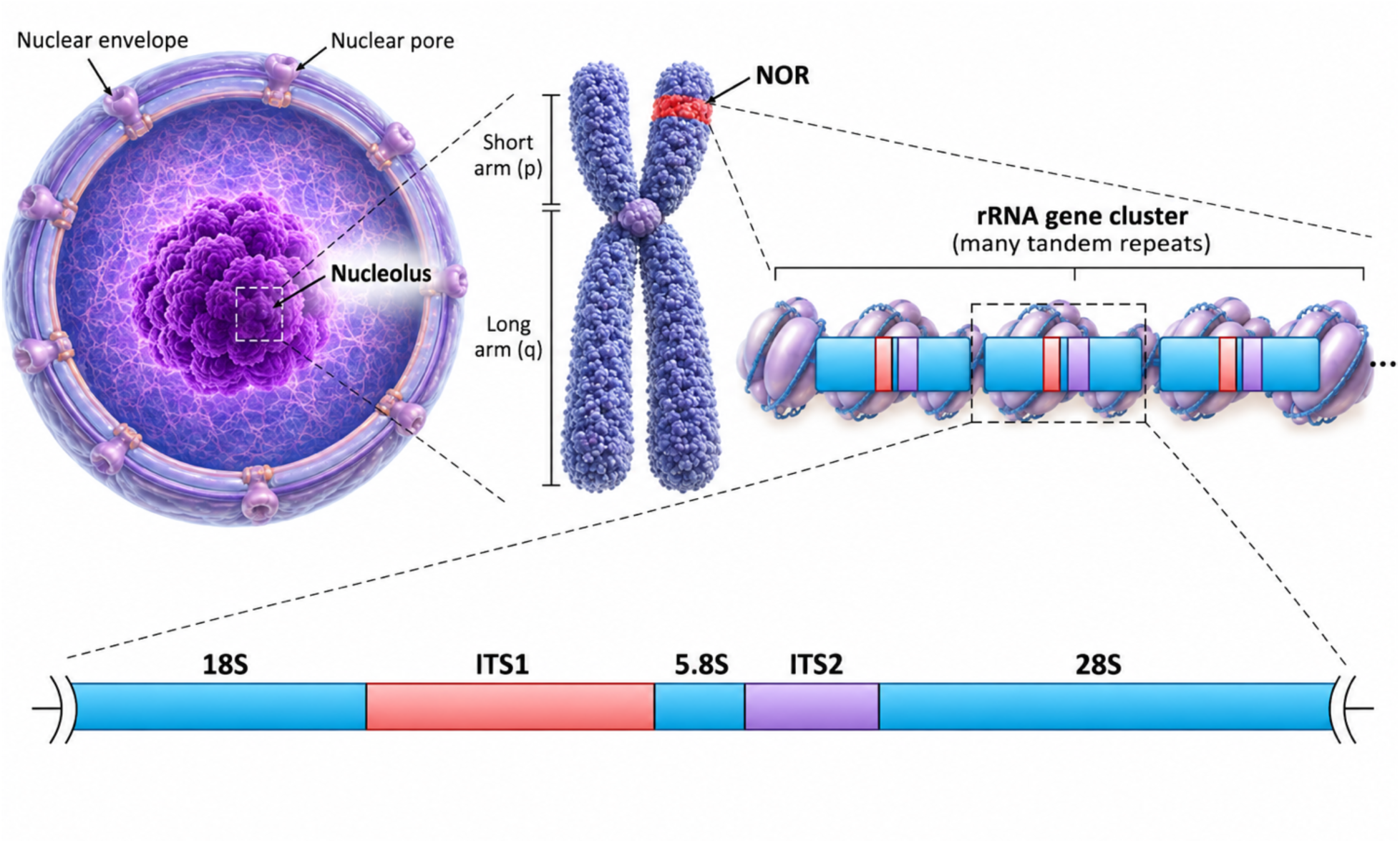
A pictorial representation of human Nucleolus Organizer Region (NOR) and rRNA gene cluster with a close-up of Internal Transcribed Spacer 1 (ITS1) and 2 (ITS2).

The evolutionary impact of viruses is not merely theoretical. Comparative analyses have revealed pervasive signatures of recurrent positive selection in antiviral genes, including the primate restriction factor *APOBEC3G*, whose adaptive evolution predates the emergence of modern lentiviruses (Sawyer et al., 2004). More broadly, viruses have been estimated to account for nearly 30% of adaptive amino acid substitutions in the portion of the human proteome conserved across mammals (Enard et al., 2016). These findings establish viral antagonism as a major evolutionary force extending well beyond genes traditionally associated with immunity. Could similar evolutionary pressures have shaped the rapidly evolving, noncoding sequences embedded within the otherwise highly conserved ribosomal RNA system? The combination of extensive sequence turnover, concerted evolution, and the universal dependence of viruses on host ribosomes makes eukaryotic ITS regions intriguing candidates for investigating this possibility. In principle, molecular drive could facilitate the spread of advantageous sequence variants across rDNA arrays, while recurrent viral selection could influence which variants persist. Whether such an evolutionary interaction exists, however, has remained unexplored.

Here, we investigate whether the eukaryotic 45S rRNA system contains an overlooked component of antiviral defense. We begin by examining sequence relationships between ITS regions and DNA viruses across diverse eukaryotic lineages, drawing on the high-quality, chromosome-scale vertebrate genome assemblies generated by the Vertebrate Genomes Project (VGP) (Formenti et al., 2026a; Kang et al., 2026). We ask whether similarities between ITS sequences and viral genomes exhibit phylogenetic and genomic patterns extending beyond compositional resemblance. We then examine how ITS-derived RNAs respond to herpesvirus infection across multiple viral subfamilies, viral species and strains, and human and animal cell systems. By integrating transcriptomic datasets spanning different infection stages, host backgrounds, and cellular compartments, we seek to distinguish general features of the ITS response from those associated with particular host–virus interactions. HSV-1 provides a particularly informative experimental system because extensive datasets permit the separation of nascent transcription, steady-state RNA accumulation, subcellular redistribution, and the effects of defined viral perturbations.

A central challenge is distinguishing two fundamentally different biological explanations for infection-associated ITS dynamics. Under a passive-remodeling hypothesis, changes in ITS abundance arise from viral interference with rRNA transcription, processing, or RNA stability. Under an active-deployment hypothesis, ITS-derived RNA participates in a host response that influences viral replication. These explanations are not necessarily mutually exclusive, and both may reproduce superficially similar RNA trajectories. Bayesian model comparison is particularly well suited to this problem because it evaluates competing biological mechanisms rather than merely testing whether individual parameters differ from zero or whether a model can reproduce the observed data (Gelman et al., 2013a; Gelman et al., 2013b; Kass and Raftery, 1995). By integrating likelihoods across the full parameter space, Bayesian marginal likelihoods account for parameter uncertainty and naturally penalize unnecessary complexity, allowing simpler and more elaborate mechanistic explanations to compete on a common evidential scale (MacKay, 2003). This is especially important when alternative biological processes can generate similar temporal patterns and when the available datasets differ in sampling density, experimental design, and measurement uncertainty (Schoniger et al., 2014; Toni et al., 2009). We therefore develop a quantitative framework that explicitly compares alternative kinetic models, including models permitting passive remodeling and active deployment to operate simultaneously. We further connect these dynamical analyses to reciprocal experimental manipulation of ITS2, using both RNA depletion and synthetic RNA supplementation to test whether ITS2 influences viral replication independently of the correlations observed during infection.

By combining high-quality comparative genomics, a broad survey of herpesvirus infections across viral lineages and host cell systems, infection-resolved transcriptomics, Bayesian mechanistic modeling, and reciprocal functional perturbations, we ask whether sequences conventionally viewed as transient byproducts of ribosome biogenesis have acquired a second biological role. More broadly, this approach tests the possibility that the eukaryotic machinery responsible for producing ribosomes also contributes to restricting the viruses that depend on them.

## Results

### 1. Ribosomal RNA spacers and DNA virus genomes exhibit sequence convergence

We notice that ITS1 and ITS2 of eukaryotic ribosomal DNA contain segments with substantial nucleotide similarity to DNA virus genomes, despite the fact that ITS sequences are GC-rich, evolve rapidly, and contain abundant low-complexity motifs (**Figure 2**). For example, a 78 bp-long fragment of fairyfly’s (*Gonatocerus morgani*) ITS2 bears a resemblance (e-value=6×10^−13^) to a moth’s *Amsacta moorei* entomopoxvirus **(Figure 2A)**. Both ITS1 and ITS2 of the fruitfly *Drosophila melanogaster* contain 25-75 bp-long sequences highly homologous to Ascoviruses (e-value = 2 × 10^−7^), large double-stranded DNA insect viruses that invade and destroy their host’s cell nuclei **(Figure 2A)**. As much as 11.5% of the chicken’s ITS1 sequence has a 73% or higher identity with a gamma-herpesvirus (retroperitoneal fibromatosis-associated herpesvirus, e-value= 2×10^−10;^ **Figure 2A**), whereas mammalian ITS regions are enriched with alpha-herpesvirus-like motifs, consistent with the congruence between mammalian and alpha-herpesviral phylogenies and cospeciation (McGeoch et al., 2000). We therefore systematically examined whether the distribution of these similarities was consistent with sequence composition alone or showed additional biological structure, particularly enrichment among specific host–virus taxonomic associations. We drew on high-quality genomic resources, including Vertebrate Genomes Project-derived ITS sequences where available (Formenti et al., 2026b; Kang et al., 2026), and examined the sequence context of individual matches. This was particularly important for ribosomal DNA, whose tandemly repeated organization complicates assembly and representation in reference genomes. High-quality assemblies improve confidence in the recovered spacer sequence.

**Figure 2.**
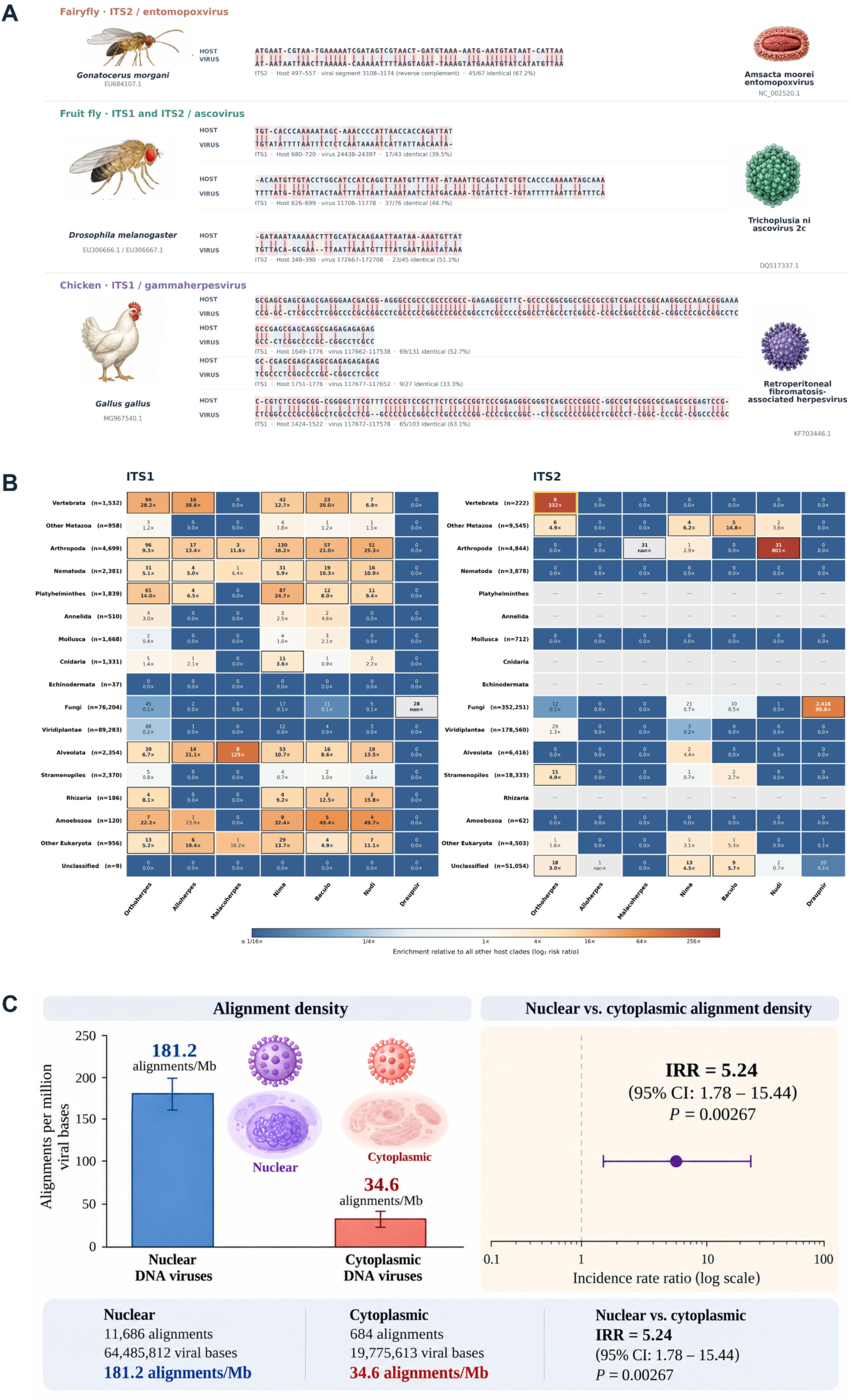
Viral sequence homology of vertebrate ITS1 and ITS2, phylogenetic distribution, and compartmental separation. (A) Representative sequence alignments between eukaryotic ribosomal internal transcribed spacers (ITS1 and ITS2) and DNA viral genomes. Three examples illustrate nucleotide sequence similarity between host ribosomal DNA spacers and phylogenetically diverse DNA viruses. **(Top)** Alignment between ITS2 of the fairyfly *Gonatocerus morgani* (Hymenoptera) and *Amsacta moorei* entomopoxvirus (Poxviridae). **(Middle)** Alignment involving ITS1/ITS2 sequences of *Drosophila melanogaster* (Diptera) and *Trichoplusia ni* ascovirus 2c (Ascoviridae). **(Bottom)** Alignment between ITS1 of the chicken *Gallus gallus* (Aves) and retroperitoneal fibromatosis-associated herpesvirus (Herpesviridae). The alignments display the corresponding host and viral nucleotide sequences, with matching positions indicated by vertical bars and gaps representing insertions or deletions introduced to optimize sequence correspondence. These examples demonstrate that detectable ITS–viral sequence similarities occur across evolutionarily distant eukaryotic hosts and distinct DNA virus families. Although such alignments identify regions of local sequence similarity, they do not independently establish shared evolutionary ancestry, horizontal sequence transfer, or functional interactions between host ITS sequences and viral genomes. **(B) Host-clade-specific enrichment of ITS1 and ITS2 sequence similarities across DNA virus families.** Heatmaps summarize the distribution and relative enrichment of ITS1 (left) and ITS2 (right) sequence similarities to seven DNA virus families across 17 eukaryotic host clades. Each cell represents a host-clade–virus-family combination, with the upper number indicating the number of distinct ITS queries exhibiting sequence similarity to that viral family and the lower number indicating the corresponding enrichment (risk ratio, ×) relative to all other host clades. Enrichment was calculated at the level of distinct ITS queries rather than individual alignment hits (HSPs), avoiding inflation from multiple alignments involving the same query. Cell colors represent log₂-transformed risk ratios, ranging from depletion (blue) through no enrichment (near-white) to increasing enrichment (orange–red). The number of ITS queries assigned to each host clade is indicated in parentheses (*n*) alongside the clade name. Statistical significance was evaluated using Fisher’s exact tests with Benjamini–Hochberg correction for multiple comparisons. Cells meeting a false discovery rate (FDR) threshold of 0.05 are outlined and displayed in bold type. Gray cells marked with a dash indicate combinations lacking an entry in the source dataset and should not be interpreted as zero similarity or absence of enrichment. The ITS2–Vertebrata–Orthoherpesviridae combination is additionally highlighted with a yellow border to emphasize its relevance to the proposed relationship between ITS2 and herpesvirus infection. Together, the two heatmaps reveal the taxonomic heterogeneity of ITS–viral sequence similarities and distinguish broadly distributed matches from host-clade-specific enrichments. **(C) Comparison of the density of ITS1/ITS2 sequence alignments across DNA virus genomes classified according to their nuclear or cytoplasmic replication compartments. (Left)** Alignment density, expressed as the number of high-scoring segment pairs (HSPs) per million viral nucleotides (Mb), was substantially higher in nuclear DNA viruses (181.2 alignments/Mb) than in cytoplasmic DNA viruses (34.6 alignments/Mb). The analysis included 11,686 alignments across 64,485,812 nucleotides of nuclear viral genomes and 684 alignments across 19,775,613 nucleotides of cytoplasmic viral genomes. **(Right)** The difference was evaluated using Poisson regression with viral sequence length as an exposure offset and heteroskedasticity-consistent (HC3) robust standard errors. Nuclear DNA viruses exhibited a 5.24-fold higher alignment incidence rate than cytoplasmic DNA viruses (incidence rate ratio [IRR] = 5.24; 95% confidence interval [CI], 1.78–15.44; *P* = 0.00267). The purple point represents the estimated IRR, and the horizontal line indicates its 95% CI on a logarithmic scale. The vertical dashed line marks the null expectation (IRR = 1). The bottom panel summarizes the alignment counts, total viral sequence lengths, alignment densities, and regression statistics. Together, these results demonstrate a pronounced enrichment of ITS–viral sequence similarity among nuclear DNA viruses, consistent with a potential relationship between viral nuclear localization and the occurrence of ITS-homologous sequences.

Following moderate filtering of mononucleotide- and dinucleotide-repeat-dominated alignments, 336,232 ITS–DNA virus alignments were retained. Their distribution differed markedly among host clades and viral families, revealing several prominent taxonomic associations that broadly reflected the known host ranges and ecological relationships of these viruses (**Figure 2B**; **SI, Table S1**). The strongest associations involved fungal ITS2 and *Draupnirviridae*, arthropod ITSs and *Nudiviridae* or *Baculoviridae*, and vertebrate ITSs and *Orthoherpesviridae*. These associations were highly enriched, reaching 90.8-fold for fungal ITS2–*Draupnirviridae* and 800.6-fold for arthropod ITS2–*Nudiviridae* (**SI, Table S2**). Vertebrate ITS1 queries were enriched 28.2-fold for orthoherpesvirus matches relative to queries from other host clades (one-sided Fisher *P* = 5.29 × 10⁻⁹⁶); for ITS2, the corresponding enrichment was 331.8-fold (*P* = 3.18 × 10⁻²⁰; **SI, Table S3**). These analyses counted distinct ITS queries carrying at least one retained alignment, rather than treating overlapping alignments as independent observations. Although such enrichments do not identify their evolutionary origin, the concentration of similarities within particular host–virus combinations warranted closer investigation. Thus, despite the potential for spurious similarities arising from the unusual sequence composition of ITS regions, the retained alignments exhibited a striking degree of biological and ecological coherence.

Parallel comparisons involving RNA viruses did not reveal a comparable pattern of statistically significant sequence alignments surviving the same filtering criteria applied to DNA viral genomes. Unlike most RNA viruses, which replicate primarily in the cytoplasm (Hiscox, 2007), most DNA viruses replicate in the nucleus, where their genomes may come into close proximity with nucleolar rDNA, creating opportunities for direct molecular interactions and potentially even recombination between viral and host sequences. We thus asked and examined whether the distribution of ITS similarities differed according to DNA viral replication compartment. Among viral genomes with an assigned replication compartment, DNA nuclear-replicating viruses showed higher ITS alignment density than DNA cytoplasmic-replicating viruses (**Figure 2C**). In a genome-level Poisson model incorporating viral genome length as an offset, nuclear replication was associated with a 5.24-fold increase in retained alignment density (HC3-robust 95% CI, 1.78–15.44; *P* = 0.00267;). The association remained after accounting for genome length but does not distinguish shared sequence constraints from historical sequence exchange or other mechanisms.

#### Vertebrate ITS orthoherpesvirus convergence

*Orthoherpesviridae* are nuclear-replicating double-stranded DNA viruses with long evolutionary associations with vertebrates (Alba et al., 2001; Guellil et al., 2022; McGeoch et al., 2000; Whitby et al., 2003). Herpesvirus phylogenies closely parallel vertebrate and, in particular, primate phylogeny (McGeoch et al., 2000), although host switching and lineage diversification have also contributed importantly to their histories (Brito et al., 2021). Given the longstanding coevolutionary relationship between vertebrates and Orthoherpesviruses, we focused subsequent analyses on vertebrate ITS–orthoherpesvirus similarities. This background provides a relevant context for evaluating whether the ITS–viral similarities reflect shared sequence architecture, evolutionary history, or both. To examine these alternatives, we analyzed selected high-scoring loci in greater detail, comparing their positions and repeat composition, the distribution of homologous segments among related hosts, and their behavior under dinucleotide-preserving sequence randomization. The association was recovered for both ITS1 and ITS2 and was especially pronounced for ITS2. Four vertebrate examples illustrate the range of patterns recovered.

#### Rabbitfish (Siganus): an exceptional ITS1 candidate

A particularly pronounced ITS1–orthoherpesvirus sequence similarity was identified in the rabbitfish *Siganus spinus* (**Figure 3**). A discrete region of its ITS1 exhibited significant local alignments with multiple orthoherpesvirus genomes, prompting a comparative analysis of ITS1 sequences from 20 *Siganus* species against 20 selected viral genomic loci. The *S. spinus* sequence consistently produced higher alignment scores than those of its congeners. For example, against a region of the human alphaherpesvirus 2 genome (NC_001798; positions 4620–4870), the *S. spinus* ITS1 yielded an alignment score of 141, whereas the highest score among the remaining 19 species was 88 (**Figure 3A; SI, Table S4**). This marked interspecific difference indicated that the similarity was not simply a conserved feature of rabbitfish ITS1 sequences. To investigate the origin of this correspondence, we compared the candidate ITS1 region with homologous sequences from the remaining 19 *Siganus* species (**Figure 3B**). We also assessed whether the observed similarities could arise from local nucleotide composition by comparing the original ITS1 sequence with dinucleotide-preserving shuffled controls (**Figure 3C**). For two representative viral loci, the observed alignment scores exceeded those obtained from all 100 shuffled sequences (empirical *P* = 0.0099), indicating that dinucleotide composition alone did not fully account for these similarities. Despite this statistical evidence, the comparative sequence patterns did not provide convincing support for horizontal transfer from an orthoherpesvirus. The correspondence was localized and involved recurrent short sequence motifs rather than an extended, uniquely identifiable viral-derived segment. The positional distribution of alignment gaps across congeners revealed substantial heterogeneity within the candidate region, with numerous positions absent from most other *Siganus* species **(Figure 3D).** Importantly, the available alignments did not resolve a discrete insertion with identifiable host–viral junctions, nor did they establish a continuous viral-derived tract whose boundaries and sequence divergence could be reconstructed across congeners. Furthermore, similarity to multiple orthoherpesvirus loci did not identify a specific donor lineage or establish a phylogenetically coherent transfer history. The pronounced divergence between *S. spinus* and its congeners therefore cannot, by itself, distinguish a historical insertion from lineage-specific evolution of the ITS1 sequence, including repeat turnover and the accumulation of short motifs that independently resemble viral sequences. Nevertheless, these observations identify an exceptional, composition-resistant ITS1–viral sequence correspondence, while leaving its evolutionary origin unresolved.

**Figure 3.**
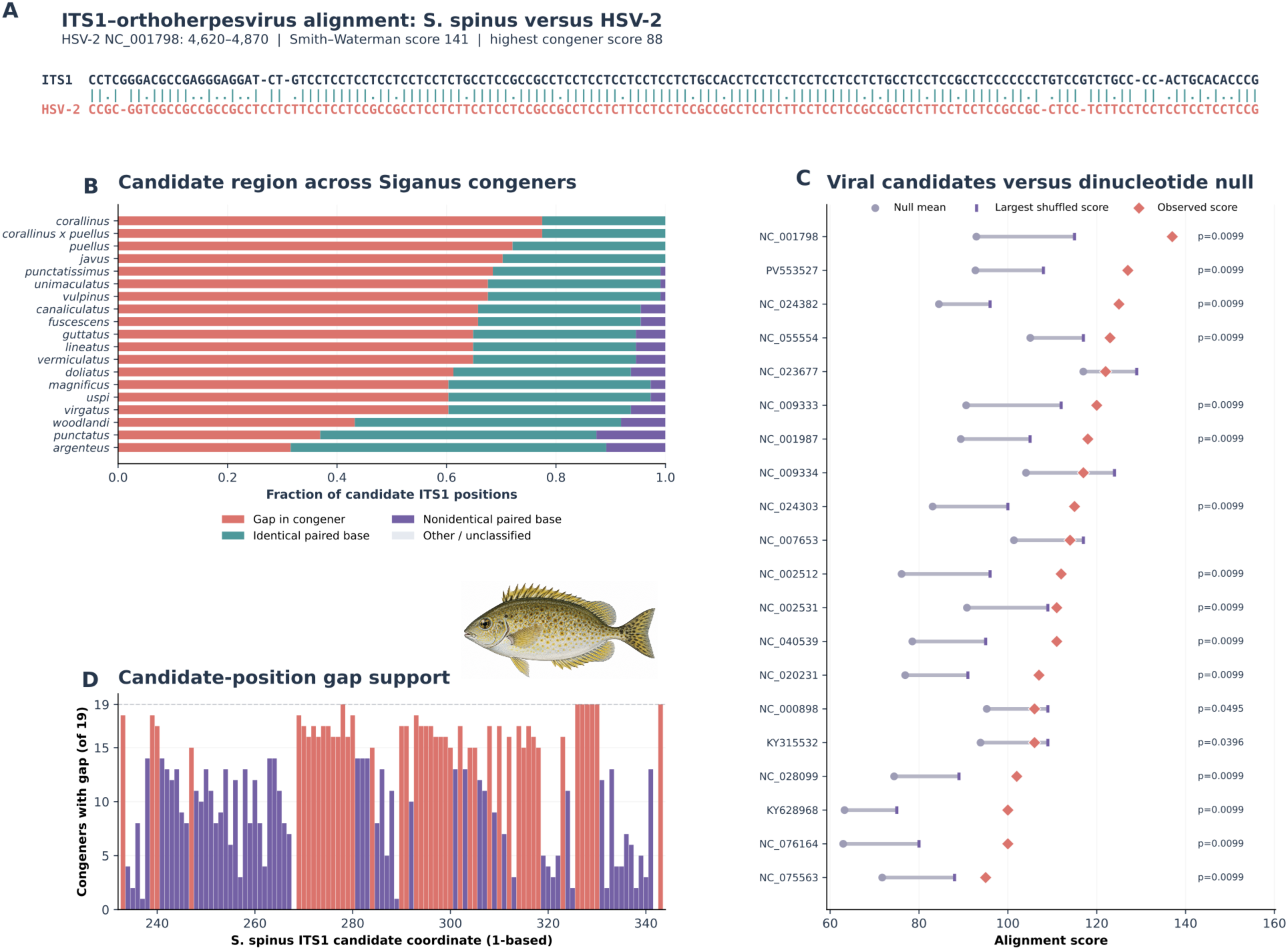
ITS1 similarity to viral genomes in *Siganus spinus* and sequence divergence among *Siganus* congeners. (A) Representative local nucleotide alignment between *S. spinus* ITS1 and human alphaherpesvirus 2 (HSV-2; NC_001798, genomic interval 4620–4870). The alignment was reconstructed using Smith–Waterman local alignment with match, mismatch, and gap scores of +2, −2, and −3, respectively. Vertical bars indicate identical nucleotides; periods indicate mismatches, and spaces indicate gaps. The observed alignment score was 141, compared with a maximum score of 88 among the remaining 19 *Siganus* species. (B) Comparative alignment of the *S. spinus* ITS1 candidate region against 19 congeners. Horizontal stacked bars indicate the fractions of candidate positions corresponding to gaps (coral), identical aligned nucleotides (teal), nonidentical aligned nucleotides (purple), or other/unclassified alignment states (light gray). **(C) Comparison of observed alignment scores for 20 viral candidates** (coral diamonds) with the mean (gray circles) and maximum (purple markers) scores obtained from 100 dinucleotide-preserving shuffled controls per candidate. Horizontal lines connect the null mean and maximum, and empirical *P* values are displayed for candidates meeting the nominal *P* ≤ 0.05 threshold. **(D) Position-specific distribution of gaps across the 111-nt candidate region.** Bars indicate the number of congeners exhibiting a gap at each position relative to *S. spinus*. Coral identifies positions with gaps in at least 15 congeners; purple indicates lower frequencies. The dashed line marks the maximum possible count of 19 congeners.

#### Anolis: repeated correspondence within extreme GC-rich ITS1

We next examined ITS1–orthoherpesvirus similarities in *Anolis*, a diverse lizard genus that provided an opportunity to assess the distribution of viral-like sequences among related species (**SI, Table S5**).

Multiple*Anolis* ITS1 sequences contained localized regions with significant similarities to orthoherpesvirus genomes (**Figure 4A-B**). In *A. garmani*, a 96-nt interval (positions 226–321) generated 14 retained alignments involving eight viral accessions, whereas a 119-nt interval in *A. cristatellus* produced 18 alignments involving seven viral accessions. Both regions were exceptionally GC-rich (87.5% and 93.3%, respectively) and contained abundant repetitive sequence, with approximately 65–67% of their nucleotides associated with recurrent 8-mer motifs (). Thus, the apparent viral similarities were concentrated within ITS1 regions sharing distinctive compositional and repetitive features. Comparative examination of these intervals suggested that their resemblance to viral sequences was largely associated with the underlying architecture of the host spacers rather than with clearly delineated viral-derived insertions. In particular, the concentration of alignments within short, highly GC-rich, repeat-enriched regions complicates their interpretation as evidence of horizontal transfer. Under a recombination/introgression scenario, one would expect additional support from extended donor-specific sequence correspondence, identifiable insertion boundaries, or a phylogenetically interpretable distribution of the putative insertion among related species. The observed similarities instead involved multiple viral accessions and localized host intervals dominated by recurrent sequence motifs, without establishing a unique viral donor or a discrete transferred segment. These features are compatible with independent accumulation or turnover of GC-rich motifs within rapidly evolving ITS1 sequences, potentially generating convergent similarities to viral genomes. Although ancient sequence exchange cannot be excluded, the *Anolis* comparisons illustrate how shared spacer architecture can produce taxonomically recurrent ITS–viral similarities without requiring repeated horizontal transfer events.

**Figure 4.**
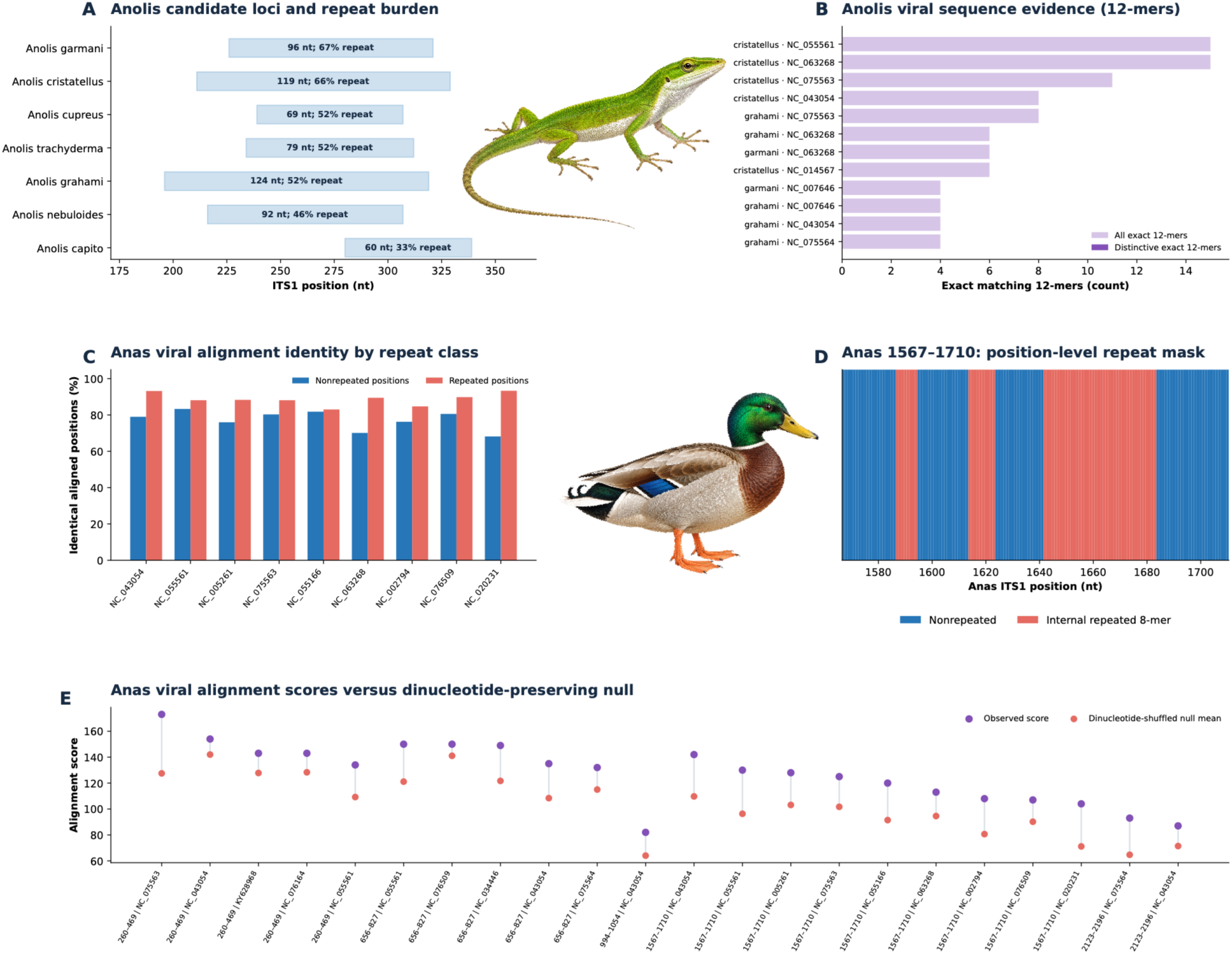
Comparative analysis of ITS1 regions exhibiting sequence similarity to orthoherpesvirus genomes, including representatives of the Alphaherpesvirinae and Betaherpesvirinae, examining the contribution of internally repeated sequences and the persistence of sequence similarity in nonrepetitive regions. (A) Positions and repeat composition of seven orthoherpesvirus-matching ITS1 regions from seven *Anolis* species. Horizontal rectangles indicate the positions and lengths of the candidate regions within the respective ITS1 sequences. Annotations specify region length (nt) and the proportion of nucleotides associated with internally repeated 8-mers. This panel characterizes the organization and repeat content of the candidate regions rather than the extent of their similarity to orthoherpesvirus genomes. (B) Sequence similarity between *Anolis* ITS1 regions and orthoherpesvirus genomes, quantified by the number of exact matching 12-mers. Light purple bars represent all exact 12-mer matches, whereas dark purple bars indicate distinctive matches, providing a more conservative measure of sequence correspondence after accounting for broadly shared motifs. Viral reference sequences are identified by GenBank accession numbers. (C) Percentage of identical aligned positions in repeated and nonrepeated portions of the candidate *Anas* ITS1 region (positions 1567–1710) across comparisons with orthoherpesvirus genomes. Blue bars indicate sequence identity at nonrepeated positions, whereas coral bars indicate identity at positions associated with internally repeated 8-mers. Percentages were calculated separately for each category using the corresponding number of aligned positions as the denominator. (D) Distribution of internally repeated and nonrepeated nucleotides across positions 1567–1710 of *Anas* ITS1. Coral indicates positions belonging to internally repeated 8-mers, whereas blue indicates nonrepeated positions. The position-specific repeat classification provides the basis for the comparisons of sequence identity shown in (C). (E) Observed alignment scores between five candidate *Anas* ITS1 regions and orthoherpesvirus sequences compared with dinucleotide-preserving shuffled controls. Purple points represent observed alignment scores, and coral points indicate the mean scores obtained from 100 dinucleotide-preserving shuffled sequences. Connecting lines illustrate the difference between observed and null mean scores for each comparison. Labels identify ITS1 coordinates and the corresponding viral reference sequences.

#### Anas: Recurrent sequence motifs contribute to ITS1–orthoherpesvirus similarities

We next examined the distribution and sequence composition of ITS1–orthoherpesvirus similarities in *Anas* ducks (**SI, Table S6**). Unlike the relatively localized candidates identified in*Siganus* and *Anolis*, the analyzed *Anas* ITS1 sequence contained multiple regions with significant viral alignments, including intervals at positions 260–469, 656–827, 994–1054, and 1567–1710, collectively spanning approximately one-quarter of the ITS1 sequence. This dispersed distribution prompted us to investigate whether the different regions represented independent instances of viral sequence similarity or reflected recurrent motifs within the host spacer. The interval at positions 1567–1710 was particularly informative because it aligned to several distinct orthoherpesvirus genomic regions. We therefore examined the contribution of repetitive sequences to these alignments by comparing sequence identity at repeat-associated and nonrepeated positions (**Figure 4C**). Against viral accession NC_043054, 64 of 81 aligned nonrepeated positions (79.0%) were identical, compared with 55 of 59 repeat-associated positions (93.2%). Thus, although repetitive motifs contributed disproportionately to sequence identity, substantial similarity also extended beyond the repeat-associated positions. Mapping the distribution of internally repeated 8-mers across the 1567–1710 interval further revealed the positional organization of repetitive and nonrepetitive sequences underlying these comparisons (**Figure 4D**). Comparisons among five candidate ITS1 intervals further identified shared short sequence motifs, indicating that the apparent correspondence to multiple viral regions was at least partly associated with recurrent sequence features within the host spacer (). To determine whether these similarities exceeded expectations based on nucleotide composition, we compared the observed alignments with those obtained from dinucleotide-preserving randomized sequences (**Figure 4E**). The results varied among ITS1 intervals and viral targets: some observed scores exceeded the corresponding shuffled distributions, whereas others fell within the range expected under the compositional null model. These differences indicate that the viral similarities cannot be attributed uniformly either to nucleotide composition or to a distinct evolutionary origin. In particular, the recurrence of related motifs across spatially separated ITS1 intervals, together with their correspondence to multiple viral regions, does not support a straightforward interpretation involving discrete integrations from identifiable viral donors.

Conversely, the persistence of substantial nonrepeat sequence identity and the elevated scores of selected alignments suggest that simple dinucleotide composition does not fully explain every correspondence. The *Anas* results therefore highlight the contribution of higher-order motif organization to ITS1–orthoherpesvirus sequence similarity while leaving open the possibility that additional evolutionary events contribute to particular regions.

#### Primates: Extensive ITS similarities converge on shared orthoherpesvirus genomic regions

A particularly intriguing pattern emerged from comparisons involving primate ITS1 sequences (**Figure 5**; **SI, Table S7**). Representatives of seven genera – *Homo*, *Pongo*, *Macaca*, *Pan*, *Gorilla*, *Callithrix*, and *Nomascus* – exhibited local sequence similarities to a common orthoherpesvirus genomic region spanning approximately 136.6–137.4 kb (**Figure 5A; SI, Table S7**). Unlike the relatively short, repeat-rich candidate intervals identified in several other vertebrates, these alignments frequently encompassed substantial portions of the host ITS1 sequences. In six of the seven genera, the corresponding ITS1 intervals extended across 50.0–61.7% of the full spacer, with alignment spans ranging from 666 to 852 nt. The most extensive intervals were observed in *Nomascus* (852 of 1,381 nt; 61.7%), *Gorilla* (797 of 1,305 nt; 61.1%), and *Homo* (826 of 1,398 nt; 59.1%). Although the aligned sequences contained mismatches and gaps, they nevertheless exhibited appreciable nucleotide identity: 69.9% across 765 aligned pairs in *Nomascus*, 66.4% across 767 pairs in *Gorilla*, and 69.3% across 755 pairs in *Pongo* (**Figure 5B**). *Pan* showed a more restricted correspondence, involving 218 nt, or 16.1% of its ITS1 sequence. Thus, the shared viral-coordinate association was not confined to isolated short motifs but extended over substantial, partially corresponding regions of primate ITS1 sequences. Closer examination revealed a more strongly conserved local component within this broader correspondence. Near viral coordinate 137 kb, a *Pongo* ITS1 segment exhibited 107 identical nucleotides across 125 aligned positions (85.6% identity), with comparable local similarities in *Gorilla* (103/122; 84.4%) and *Nomascus* (93/112; 83.0%). To assess whether these alignments could be explained by nucleotide composition, we compared the observed scores against distributions generated from 100 dinucleotide-preserving randomized sequences. For the 137-kb region, the observed alignment scores exceeded all randomized controls in each of the three genera (empirical *P* = 0.0099). The corresponding scores were 133, 132, and 118 for *Pongo*, *Gorilla*, and *Nomascus*, respectively, compared with null-distribution means of 97.9, 102.6, and 91.6. In contrast, alignments involving a separate viral region near 104 kb were not distinguishable from their respective randomized distributions (*P* = 0.891, 0.871, and 0.634; **Figure 5C-D**). These contrasting outcomes demonstrate that the correspondence cannot be explained uniformly by dinucleotide composition and that its statistical distinctiveness depends strongly on the particular viral region examined. The convergence of ITS1 sequences from phylogenetically separated primate lineages onto a narrowly defined viral genomic neighborhood is notable, particularly given the extensive proportions of the host spacers involved.

**Figure 5.**
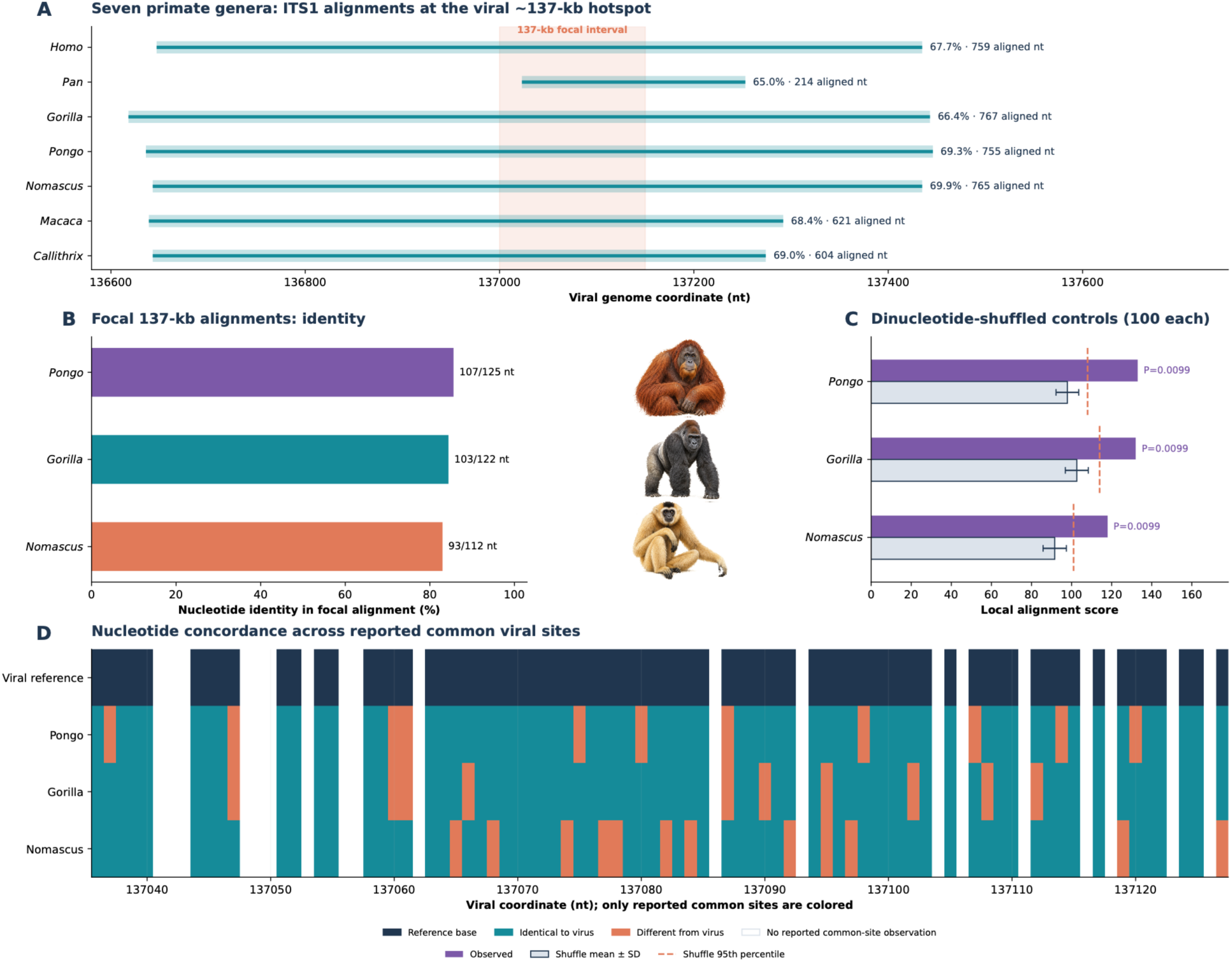
Conserved ITS1 sequence similarity to a viral genomic hotspot near 137 kb across seven primate genera. Comparative analysis of primate ITS1 sequences and their nucleotide similarity to a localized region of an orthoherpesvirus genome, integrating cross-taxon alignment conservation, sequence identity, and dinucleotide-preserving shuffled controls. **(A) Distribution of ITS1 alignments along the viral genome near position 137 kb across seven primate genera (*Homo, Pan, Gorilla, Pongo, Nomascus, Macaca,* and *Callithrix*).** Horizontal segments indicate the viral coordinates spanned by each alignment, with annotations reporting nucleotide identity and the number of aligned nucleotide pairs. The shaded interval highlights the focal region near 137 kb, revealing the spatial clustering of ITS1–viral sequence similarities across evolutionarily divergent primate lineages. **(B) Nucleotide identity of focal ITS1–viral alignments in *Pongo, Gorilla,* and *Nomascus*.** Bars indicate the percentage of identical nucleotides within each alignment, with annotations reporting the numbers of identical and aligned bases. **(C) Comparison of observed local alignment scores with expectations under a dinucleotide-preserving shuffled-sequence null model for the same three genera.** Purple bars represent observed scores, gray bars indicate mean scores from 100 shuffled controls (±SD), and dashed coral lines mark the corresponding 95th percentiles of the null distributions. Empirical *P* values were calculated from the shuffled controls (*P* = 0.0099 for each genus), indicating that the observed sequence similarities exceed those obtained under the tested compositional null model. **(D) Nucleotide-level concordance between the viral reference sequence and ITS1 sequences from *Pongo, Gorilla,* and *Nomascus* at reported common viral positions.** Dark blue denotes viral reference nucleotides, teal indicates identical nucleotides in the corresponding primate sequence, and coral indicates mismatches; white positions lack reported common-site observations. The comparison illustrates the distribution of conserved and divergent nucleotides within the shared hotspot. Illustrations depict representatives of *Pongo, Gorilla,* and *Nomascus*, aligned with their corresponding results in panels B and C. Together, these analyses identify a localized region of ITS1–viral sequence similarity shared across multiple primate lineages, with focal matches in three genera exceeding expectations based on dinucleotide composition alone.

The association between primate ribosomal spacers and orthoherpesvirus genomes was not restricted to ITS1. Indeed, the broader taxonomic analysis revealed a particularly pronounced enrichment of ITS2–*Orthoherpesviridae* similarities among vertebrates (331.8-fold relative to other host clades), with human ITS2 sequences contributing prominently to the retained alignments. Approximately 26% of human and 18% of chimpanzee ITS2 regions exhibit homology with alpha-herpesvirus genomes, and these specific motifs significantly contribute to the sequence divergence observed between human and chimpanzee ITS2s: while human and chimpanzee ITS2 sequences are 88% identical (Needleman-Wunsch score 654), 63% (n=90) of their divergent single nucleotide polymorphisms (SNPs) map to the herpesvirus homology areas (Fisher’s exact test *P* = 1×10^−4^). Examination of individual human ITS2 sequences identified several discrete regions exhibiting similarity to different orthoherpesvirus genomic loci. For example, a 173-nt alignment between human ITS2 and human alphaherpesvirus 2 (NC_001798) exhibited 74.0% nucleotide identity and encompassed approximately 14.4% of the 1,155-nt host spacer. Additional alignments involved a 167-nt region with 74.3% identity to NC_007653 and a shorter, 72-nt region with 83.3% identity to NC_043054. Closely corresponding alignments were also recovered from an independently cataloged human ITS2 sequence, indicating that the pattern was reproducible across database representations of the human spacer. The ITS2 similarities differed in their organization from the extensive ITS1 alignments observed across several primate genera. Whereas the latter frequently encompassed more than half of the host spacer and converged on a common viral genomic neighborhood, the human ITS2 correspondences were concentrated within shorter intervals distributed among several viral loci. The extensive sequence correspondence across phylogenetically divergent primate genera, particularly its convergence on the same viral genomic region, raises intriguing questions about the evolutionary origin of these similarities. Although the broader alignments exhibit moderate nucleotide identity, they contain shorter segments of substantially higher conservation, some of which exceed expectations based on dinucleotide composition alone. The recurrence of these features across multiple primate lineages suggests an evolutionarily persistent component of ITS sequence architecture, potentially reflecting shared ancestry, sequence constraints, or historical interactions between host and viral genomes. Importantly, even if these similarities initially arose through neutral sequence evolution or coincidental convergence, their emergence could have created opportunities for molecular interactions between host and viral nucleic acids, potentially initiating new selective pressures, genetic conflicts, and coevolutionary dynamics. Thus, the extensive ITS similarities concentrated within a narrowly defined orthoherpesvirus genomic region may be functionally consequential regardless of their exact mechanism of origin.

#### RNA secondary structure of vertebrate ITS regions exhibiting orthoherpesvirus sequence similarity

We next investigated the predicted RNA secondary structures of ITS1 and ITS2 regions from diverse vertebrate hosts that exhibited nucleotide sequence similarity to orthoherpesvirus genomes (**SI, Table S8**). The host sequences represented mammals, birds, reptiles, amphibians, and fishes, including primates (*Homo, Gorilla,* and *Callithrix*), birds (*Anas* and *Gallus*), reptiles (*Anolis*), amphibians (*Xenopus*), and numerous teleost fishes (*Siganus, Anchoa, Acanthurus,* and others). The corresponding viral sequences were genomic segments from multiple orthoherpesviruses, including the reference genomes NC_063268, NC_075564, and NC_043054. We examined three distinct structural contexts: the behavior of virus-matching host ITS regions within full-length ITS RNA molecules, the intrinsic folding properties of those regions when isolated from their native spacers, and the predicted RNA folding properties of the matching orthoherpesvirus genomic segments. We first evaluated the extent of base pairing within predicted full-length host ITS secondary structures. The virus-matching regions exhibited modestly but consistently elevated fractions of paired nucleotides relative to compositionally matched control regions from the same host spacers. The mean increase was 4.32 percentage points for ITS1 (58 intervals) and 2.21 percentage points for ITS2 (29 intervals). Positive differences occurred in 41 of 58 ITS1 intervals (70.7%) and 22 of 29 ITS2 intervals (75.9%), significantly exceeding the expectation of equal frequencies of positive and negative differences (two-sided exact sign tests, *P* = 0.00223 and *P* = 0.00813, respectively). Thus, vertebrate ITS regions exhibiting orthoherpesvirus sequence similarity tended to participate more extensively in predicted RNA secondary structure than comparable regions elsewhere within the same host ITS molecules. We next examined whether this enrichment reflected an intrinsic propensity of the host ITS regions to fold independently or depended on their surrounding RNA context. When extracted from their native spacers and folded as isolated fragments, the virus-matching host ITS regions exhibited less favorable predicted folding than GC-matched within-spacer controls (median minimum-free-energy [MFE] Z-scores of +0.940 for ITS1, *n* = 67, and +1.524 for ITS2, *n* = 33). These differences diminished when 50-nt flanking sequences were included. Together, these results suggest that the increased base pairing observed within full-length vertebrate ITS molecules depends substantially on interactions with surrounding spacer sequences rather than unusually stable autonomous folding of the matching fragments.

We then analyzed the corresponding orthoherpesvirus genomic segments identified through their sequence similarity to these vertebrate ITS regions. For each viral segment, RNA secondary structure was predicted computationally from the viral DNA sequence and compared with compositionally matched control sequences from the same viral genome. Unlike the host ITS regions examined within full-length RNA molecules, the viral segments generally exhibited less favorable folding and reduced predicted base pairing relative to their matched viral controls.

Among 112 viral segments analyzed independently, the median MFE Z-score was +3.065, with 92 regions (82.1%) exhibiting positive Z-scores. Their median paired-nucleotide fraction was 32.7 percentage points lower than that of matched viral controls. Extending the viral segments by 50 nt on either side reduced these differences: the median MFE Z-score decreased to +1.325 across 127 regions, while the median deficit in predicted base pairing narrowed to 9.9 percentage points.

These analyses reveal a context-dependent difference in the predicted structural properties of vertebrate ITS regions and their matching orthoherpesvirus genomic segments. Within full-length host ITS RNA molecules, regions exhibiting orthoherpesvirus sequence similarity tend to participate more extensively in predicted secondary structure than matched host controls.

However, when isolated from their surrounding ITS sequences, these same regions exhibit less favorable folding. The corresponding viral genomic segments likewise generally exhibit reduced predicted folding stability and base pairing relative to controls from their respective viral genomes. Thus, sequence similarity between vertebrate ITS regions and orthoherpesvirus genomes does not necessarily imply equivalent RNA structural properties. Instead, the results emphasize the importance of the surrounding sequence environment in determining the structural behavior of matching nucleotide sequences.

### 2. 45S rRNA internal transcribed spacers undergo dramatic, sequence-dependent remodeling in response to *Orthoherpesviridae* infections

The presence of virus-like sequences in eukaryotic ITS regions raises an intriguing question about their potential role and significance in the context of immunity to viral infections. First, we asked whether experimentally controlled herpesvirus infections change the abundance and intramolecular stoichiometry of ITS1 and ITS2 relative to the mature-rRNA portions of the same 45S precursor, and whether any such remodeling is spatially structured or preferentially associated with host–viral sequence resemblance. To this end, we quantified infection-associated changes in ITS1 and ITS2 across multiple independent mouse and human RNAseq datasets, with a variety of host cell types and viral strains from all three major *Orthoherpesviridae* subfamilies, gamma-, beta-, and alphaherpesviruses. Infection-associated changes in ITS1 and ITS2 were quantified both as absolute fold changes and relative to the remainder of the 45S precursor, with the differential spacer statistic D measuring the infection-induced change in the ITS2/ITS1 ratio. Robustness was assessed using independent alignment-depth, unique-fragment, and exact canonical 31-mer measurements, while replicate uncertainty was propagated using Bayesian posterior inference with 95% credible intervals. Positional and viral-homology associations were evaluated against spatially preserving circular-shift and block-bootstrap nulls, with dinucleotide-preserving shuffled viral genomes providing an additional test of viral-sequence specificity.

#### Gammaherpesvirus infection

The *Gammaherpesvirinae* is a large subfamily of the family *Herpesviridae*. Gammaherpesviruses usually cause limited disease upon primary infection, but establish lifelong infections, associated with numerous malignancies, including cancer. They have been observed to establish a variety of gene expression programs during latent and lytic infection. This capacity enables the virus to control host-cell proliferation, prevent programmed cell death, elude immune cell detection, and ultimately adapt to a wide range of environmental and developmental changes in the host cell. We reanalyzed a 4-thiouridine (4sU) nascent-RNA dataset generated in a mouse embryonic fibroblast cell line (NIH 3T3) infected for 24 h with wild-type murine gammaherpesvirus 68 (wild-type MHV68 or MHV68.ΔHS), alongside mock-infected cells (Abernathy et al., 2015) (**SI, Table S9**). MHV68 infects and replicates efficiently in several laboratory cell lines, serving as an attractive model to study the lytic replication of gammaherpesviruses (Rajcani and Kudelova, 2005), while 4sU-labeled RNA can be used to track genome-wide nascent RNA kinetics (Barrass et al., 2015; Rabani et al., 2011). The ΔHS virus carries the R443I substitution in ORF37/muSOX that selectively impairs the mRNA-degradation/host-shutoff function while preserving the essential viral nuclease framework (Abernathy et al., 2015).

The dominant MHV68 result was not a subtle local fluctuation but a broad reorganization of the rDNA transcriptional landscape. At 24 h after wild-type infection, newly synthesized ITS1 and ITS2 signals declined to 0.182× and 0.179× mock levels, respectively (−2.46 and −2.48 log2). However, this depletion did not reflect a general reduction in RNA signals across the 45S transcription unit. In the same libraries, 18S and 28S signals increased to 3.65× and 3.20× mock levels, respectively, while total 45S-derived sequencing depth rose to 3.22× (**Figure 6A; SI, Table S10**). Consequently, ITS1 and ITS2 were each depleted by more than four log2 units relative to the total 45S signal (ΔITS1 = −4.146; ΔITS2 = −4.168). Importantly, the reduction in spacer-derived RNA was not confined to particular ITS regions. Every local spacer window exhibited depletion: all 10 ITS1 windows showed negative changes, with a median of −2.47 log2, as did all 11 ITS2 windows, with a median of −2.42 log2 (**Figure 6B**). The near-zero difference between the relative depletion of the two spacers (D = −0.022) is biologically informative: wild-type MHV68 did not selectively deplete one spacer but produced a coordinated reduction in both internal spacers against a background of preserved or increased mature-rRNA-region signals.

**Figure 6.**
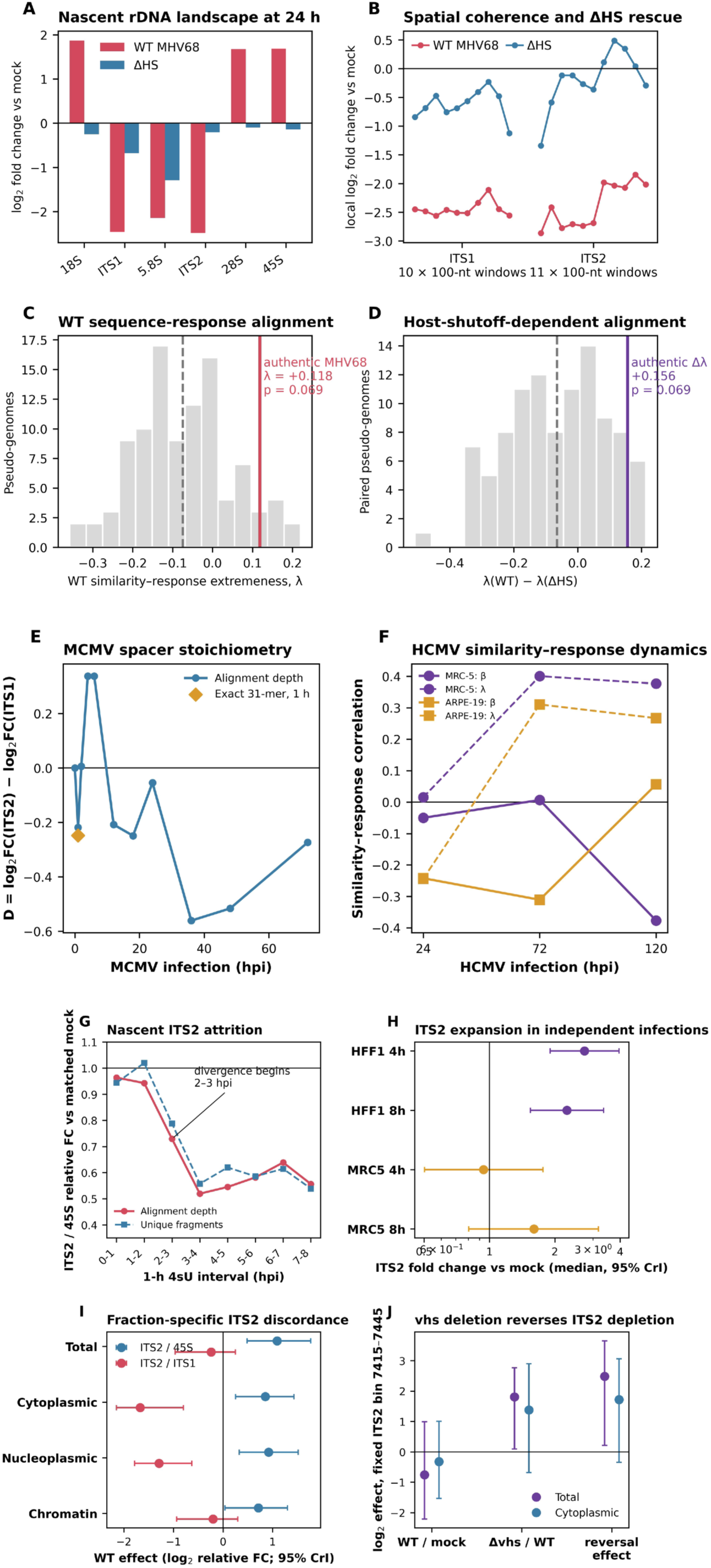
Dynamic remodeling of ITS stoichiometry and sequence-associated responses during *Herpesviridae* infection. (A) Regional changes across the 45S rDNA transcription unit and host-shutoff-dependent remodeling of nascent rDNA spacer signals during MHV68 infection. NIH 3T3 cells were infected with wild-type (WT) MHV68 or the host-shutoff-defective MHV68.ΔHS mutant, and newly synthesized RNA was analyzed 24 h post-infection from 4-thiouridine (4sU)-labeled RNA-seq data. Bars show log2 fold change relative to mock infection for 18S, ITS1, 5.8S, ITS2, 28S, and total 45S-derived signal. WT infection produced strong depletion of both internal transcribed spacers despite increased signal from mature-rRNA regions and total 45S, whereas the ΔHS mutation substantially rescued the spacer phenotype. **(B) Spatial distribution of the response across consecutive 100-nt windows spanning ITS1 and ITS2 in WT and ΔHS infection.** WT MHV68 depleted all 10 ITS1 and all 11 ITS2 windows, demonstrating spatially coherent spacer loss. In ΔHS infection, ITS1 depletion was markedly attenuated and ITS2 responses became heterogeneous and approached baseline, revealing local rescue when host shutoff was impaired. **(C) Composition-controlled test of MHV68–ITS sequence-response alignment during WT infection.** The gray distribution shows the combined-spacer similarity-response extremeness statistic (λ) obtained with 100 dinucleotide-preserving pseudo-MHV68 genomes; the dashed line denotes the pseudo-genome mean and the red line the value for the authentic MHV68 genome (λ = +0.118). The authentic genome lay near the upper tail of the composition-matched null distribution (empirical upper-tail *P* = 0.0693). **(D) MHV68 host-shutoff dependence of sequence-response alignment.** The gray distribution shows the paired pseudo-genome null distribution of Δλ = λ(WT) − λ(ΔHS), with each pseudo-genome evaluated in both infection conditions; the dashed line denotes the null mean and the purple line the observed authentic-genome contrast (Δλ = +0.156; empirical upper-tail *p* = 0.0693). Positive Δλ indicates stronger alignment between viral-sequence similarity and ITS response during WT infection than during ΔHS infection. Together, the data identify a broad, spatially coherent depletion of nascent ITS-containing RNA that depends strongly on efficient MHV68 host shutoff, accompanied by a more modest composition-controlled association between viral sequence similarity and the magnitude of the local ITS response. **(E) Dynamic remodeling of ITS stoichiometry and sequence-associated responses during MCMV infection of NIH-3T3 fibroblasts.** The trajectory shows D=\og_2_FC_ITS2_-log_2_FC_ITS1_ estimated from alignment depth, with negative values indicating preferential reduction of ITS2 relative to ITS1. An independent region-exclusive exact-31-mer estimate at 1 hpi (diamond) reproduced the early negative shift (D = −0.248 versus −0.219 by alignment). The alignment-based trajectory was non-monotonic but became consistently negative late in infection, with D = −0.561, −0.516, and −0.273 at 36, 48, and 72 hpi, respectively (late mean D = −0.450). **(F) Time-dependent relationship between HCMV–ITS sequence similarity and the host ITS response in MRC-5 fibroblasts and ARPE-19 epithelial cells.** Curves show the directional similarity-response correlation (β) and direction-independent response-extremeness correlation (λ) for ITS2 at 24, 72, and 120 hpi. In MRC-5 cells, ITS2 similarity-response coupling strengthened during infection, with λ = 0.401 at 72 hpi and, by 120 hpi, greater HCMV similarity associated simultaneously with stronger depletion (β = −0.377) and greater response magnitude (λ = 0.377). The relationship was weaker and more variable in ARPE-19 cells, although ITS2 response magnitude was positively associated with HCMV similarity at 120 hpi (λ = 0.267), illustrating the temporal and cell-context dependence of viral sequence-associated ITS remodeling. **(G) Temporal behavior of nascent ITS2 relative to the 45S rRNA precursor during the first 8 h of HSV-1 infection, quantified from 1-h 4-thiouridine (4sU) labeling intervals.** Curves show ITS2/45S fold change relative to matched mock samples estimated independently from alignment depth and region-exclusive unique fragments. ITS2 began to diverge from total 45S by 2–3 hpi and remained preferentially depleted thereafter; across 2–8 hpi, all 12 replicate-by-interval contrasts were negative, with a combined relative fold change of 0.6125. **(H) Independent evidence for infection-associated ITS2 expansion in HSV-1 infection RNA-seq datasets.** Points show Bayesian posterior median ITS2 fold changes relative to mock infection, with 95% credible intervals (CrI), in HFF1 and MRC-5 fibroblasts at 4 and 8 hpi. HFF1 cells showed an early and sustained increase in ITS2 (approximately 2.7-fold at 4 h and 2.3-fold at 8 h), whereas MRC-5 cells showed a delayed response, remaining near baseline at 4 h and increasing to approximately 1.5-fold at 8 h. **(I) Subcellular compartment-specific discordance of ITS2 during wild-type HSV-1 infection.** Bayesian posterior estimates (median and 95% CrI) compare ITS2 with total 45S-derived RNA and with ITS1 in total, cytoplasmic, nucleoplasmic, and chromatin-associated RNA. ITS2 was enriched relative to total 45S across fractions but depleted relative to ITS1 most strongly in the cytoplasmic and nucleoplasmic compartments. The joint posterior probabilities of simultaneous ITS2/45S enrichment and ITS2/ITS1 depletion were 0.9938 in cytoplasmic RNA and 0.9951 in nucleoplasmic RNA, supporting selective redistribution or differential persistence of ITS2-containing species rather than uniform modulation of the 45S precursor. (J) Dependence of a focal ITS2 depletion signal on the HSV-1 virion host shutoff factor *vhs*. Bayesian posterior estimates (median and 95% CrI) are shown for the fixed ITS2 interval 7415–7445 in total and cytoplasmic RNA: the WT/mock effect, the Δvhs/WT effect, and their reversal contrast. In total RNA, WT infection depleted the interval by −0.829 log2 whereas deletion of vhs increased it by +1.864 log2 relative to WT, yielding a +2.48-log2 reversal (95% CrI, +0.21 to +3.65; *P*[reversal > 0] = 0.982). The corresponding cytoplasmic reversal was +1.72 log2 (95% CrI, −0.35 to +3.06; *P*[reversal > 0] = 0.959). Together, these datasets reveal that HSV-1-driven ITS2 remodeling is dynamic, cell-context dependent, compartmentalized, and at least partly dependent on the viral host-shutoff machinery.

We then conducted a multiscale analysis of viral DNA–mouse ITS sequence similarity to ask whether ITS regions resembling the MHV68 genome showed systematically stronger infection-associated responses (Δᵢ). For WT infection, the combined-spacer directional correlation was β=−0.143: positions with greater MHV-68 similarity tended, modestly, to be more depleted. The direction-independent extremeness correlation was λ=+0.118, indicating that greater similarity also tended to coincide with a larger absolute response. Against 100 exact dinucleotide-preserving pseudo-MHV68 genomes, the real MHV68 sequence produced WT λ=+0.118 compared with a null mean of approximately −0.075. The authentic genome ranked about seventh highest among 101 real-plus-null genomes, giving an empirical upper-tail p=0.0693 (**Figure 6C; SI, Table S11**). The directional WT statistic was similarly unusual: β=−0.143 versus a pseudo-genome mean of approximately +0.072, again with empirical *P*=0.0693 in the depletion-oriented tail. This does not meet the conventional 0.05 threshold, but it is biologically coherent: the real infecting genome generates a local similarity landscape more aligned with WT responsiveness than roughly 94% of composition-equivalent viral sequences. The pattern was largely ITS1-centered: WT ITS1 λ=+0.224 had composition-null *P*=0.0792, while its directional β=−0.260 had *P*=0.0594. WT ITS2 showed no sequence-specific relationship (composition-null p≈0.465 for both corresponding endpoints). The same pseudo-virus panel provided a useful paired mechanistic contrast. The observed combined-spacer difference Δλ=λWT−λΔHS was +0.156 and lay near the upper edge of the paired pseudo-genome distribution (*P*=0.0693) (**Figure 6D; SI, Table S12**). For ITS1, the contrast was larger (Δλ=+0.364) and again yielded *P*=0.0693; thus, the modest sequence-response alignment is strongest in the biological condition that also produces the dramatic spacer phenotype and is weakened when viral host shutoff is disabled.

#### Betaherpesvirus infections

Betaherpesviruses typically have highly restricted host ranges, mostly to mammals. Cytomegaloviruses (CMVs) are among the most genetically tractable betaherpesviruses, with murine and human CMVs (MCMV and HCMV, respectively) in particular providing a powerful system for combining precise viral genetics with experimental infection. To test whether the abundance and intramolecular stoichiometry of ITS1 and ITS2 are consistent with the pattern observed in MHV68, we reanalyzed a dense MCMV time course dataset (GSE212286) (**SI, Table S13**), comprising duplicate mock libraries and duplicate MCMV-infected NIH-3T3 fibroblast libraries collected from 1 to 72 h post infection (hpi).

Nascent RNA was labelled with 400µM 4sU one hour prior to RNA harvesting. 4sU labelled RNA was subject to alkylation to mediate U to C conversions to track newly synthesized RNAs. The dataset also contains one 24-h MCMV+Phosphonoacetic acid (PAA) library and one 4-h MCMV+Cycloheximide (CYC/CHX) library. PAA inhibits herpesviral DNA polymerase/DNA replication and thereby prevents the normal DNA-replication-dependent late (γ) viral gene-expression program, while allowing immediate-early and early programs, whereas the CYC/CHX assay allows selective expression of viral immediate early (α) gene (Ostermann et al., 2018). This is the classic way of distinguishing early from true late herpesvirus expression.

We quantified mouse 18S, ITS1, 5.8S, ITS2, 28S and the complete transcribed 45S interval using two deliberately different sequence measurements: multi-mapping-aware Bowtie2 depth against the established mouse rDNA reference BK000964.3 (Grozdanov et al., 2003) and exact region-exclusive canonical 31-mers. The alignment-depth analysis revealed a time-dependent shift in the relative representation of ITS1 and ITS2 (**SI, Table S14**). At 1 hpi, ITS2 was reduced more strongly than ITS1, yielding D=log2FCITS2 – log2FCITS1=-0.219. Both biological replicates supported this direction, although the small replicate number precluded significance in an exact 2-versus-2 test (*P*=0.333). Notably, the independent exact-31-mer analysis reproduced the relative shift at 1 hpi (**SI, Table S15**) (D=-0.248), despite differing in absolute abundance estimates: ITS1 increased 21.4% above mock whereas ITS2 remained near baseline. Thus, both estimators independently identified an early downward shift of ITS2 relative to ITS1. The alignment trajectory was not monotonic, with positive D at 4–6 hpi followed by renewed negative values at 12–18 hpi. The strongest and most consistent alignment-based shift occurred late in infection: D=-0.561, -0.516, and -0.273 at 36, 48, and 72 hpi, respectively (mean D=-0.450) (**Figure 6E**). This late effect was concordant across biological replicates and remained strongly negative after exclusion of any individual late time point (leave-one-time-point-out mean D=-0.395 to -0.539). Statistical resolution was nevertheless limited by only three replicated late time points (exact cluster-level directional sign-flip P=0.125), so the late trajectory is supported primarily by its effect size and consistency rather than conventional significance. The homology screen produced two raw short HSPs (**SI, Table S16**), one in each spacer. ITS1 contained a 17-nt sequence (**SI, Table S17**) (CCGCGGCGGCGGCTGCT) with 100% identity to MCMV KY348373.1 at viral coordinates 130230-130214 (E = 0.012; bit score 34.2).

We then reanalyzed 24 biological HCMV libraries (**SI, Tables S18–S25**): human MRC-5 and ARPE-19 cells, each with mock and TB40/E-infected samples at 24, 72, and 120 hpi, and two biological replicates per condition (Oberstein and Shenk, 2017). Similar to MHV68 and MCMV, the central HCMV result shows a dynamic remodeling of pre-rRNA composition dependent on timing and cell type. At 24 h, HCMV is maintained in MRC-5 cells and increased in ARPE-19 cells, while ITS1 is increased in both backgrounds. At 72–120 h, absolute ITS2 generally declines, but mature-rRNA regions often decline much more strongly, producing pronounced relative preservation of spacer-containing RNA. At the same time, local ITS2 neighborhoodscan rise several-fold even when whole-ITS2 abundance falls. Analysis of human ITS- HCMV homology coupled with expression quantification revealed a strikingly dynamic relationship between viral sequence similarity and the host ITS response. In MRC-5 fibroblasts, HCMV-like ITS1 positions were preferentially induced early in infection (24 h; β = 0.368, spatial *P* = 0.0059), an effect that was directionally reproduced in every infected-versus-mock replicate comparison. By 72 h, the relationship shifted from response direction toward response magnitude: viral similarity predicted the absolute strength of the ITS response across the combined spacers (λ = 0.211, *P* = 0.030), with a particularly strong association in ITS2 (λ = 0.401, *P* = 0.037). Most remarkably, by 120 h ITS2 had undergone broad depletion (mean Δ = −1.17 log2), yet the sequence relationship did not disappear – it reversed direction. Positions with greater HCMV similarity were preferentially depleted (β = −0.377, *P* = 0.0051) while simultaneously showing greater response magnitude (λ = 0.377, *P* = 0.0051). Thus, viral sequence similarity predicted where the strongest ITS response occurred even as the biological direction of that response changed over the course of infection. The effect was weaker and less reproducible in ARPE-19 epithelial cells, although ITS2 response magnitude was again associated with HCMV similarity at 120 h (λ = 0.267, *P* = 0.0468), indicating substantial cell-state dependence (**Figure 6F**). Together, gamma- and betaherpesvirus observations suggest a broader model of dynamic ITS deployment during viral infection, in which ITS sequences emerge not as passive by-products of rRNA processing, but a potentially responsive, host-mobilized RNA repertoire whose abundance, sequence composition, and cellular distribution can be remodeled during infection. Within this scenario, local viral–host sequence similarity may help identify ITS regions that are particularly response-prone, whereas infection stage, viral biology, and cellular context determine which ITS sequences are deployed, retained, redistributed, or depleted.

#### Alphaherpesvirus infections

We next sought a system in which the proposed deployment of ITS sequences during viral infection could be examined at greater experimental depth.

Alphaherpesviruses, and HSV-1 in particular, provided a powerful setting because independent transcriptomic datasets span multiple cell types and stages of infection, nascent and steady-state RNA, subcellular RNA fractions, and genetically defined viral perturbations (Rutkowski et al., 2015; Shi et al., 2018). This experimental breadth allowed us to ask whether ITS deployment is reproducible across infections, how it develops over time, which ITS regions are preferentially mobilized or depleted, how these responses relate to local viral–ITS sequence similarity, and whether they depend on specific viral functions. We therefore used HSV-1 as the principal system for deeper computational and experimental interrogation of the ITS deployment model within a rigorous Bayesian framework. The analysis included five independent alphaherpesvirus RNA-seq datasets comprising 76 libraries and spanning early through late infections, primary HFF, HFF-1, MRC-5, and KMB17 fibroblasts infected with strain 17, along with sampling from hourly 4sU-labeled nascent-RNA intervals across the first 8 h of infection through a 48-h endpoint. A complementary fractionation experiment resolved total, cytoplasmic, nucleoplasmic, and chromatin-associated RNA at 8 h and included WT and Δvhs HSV-1, allowing the contribution of the viral host-shutoff nuclease to be examined directly (Friedel et al., 2021; Hennig et al., 2018). Additionally, we infected primary NHDFs with HSV-1 KOS strain, generated four infected and four mock-infected libraries without rRNA depletion for small RNA sequencing and fragmentome analysis.

The 4sU experiment provides the cleanest temporal evidence because it measures newly synthesized RNA without rRNA depletion (Rutkowski et al., 2015). During 0–2 hpi, total 45S and ITS2 were close to matched mock. A sharp divergence began at 2–3 hpi. Total 45S remained ∼1.02–1.05× mock through 5–6 hpi, whereas ITS2 fell to 0.74× at 2–3 h, 0.54× at 3–4 h, 0.52× at 4–5 h, and 0.61× at 5–6 h. At 3–4 hpi, newly synthesized ITS2 RNA was therefore present at only about half the abundance expected from the contemporaneous 45S pool even though total 45S itself was essentially unchanged. Unique-fragment analysis gave a geometric ITS2/45S relative FC of 0.788 at 2–3 h, 0.557 at 3–4 h, and 0.539–0.620 through 7–8 h. Across 2–8 hpi, all 12 replicate×interval contrasts were negative; combined relative FC=0.6125 (Δlog2=−0.707), ∼39% relative attrition (**Figure 6G; SI, Table S27**). Replicate 1 alone gave 0.564 and replicate 2 0.666. Raw-FASTQ 31-mers independently reproduced the spacer-wide loss: at 7–8 hpi ITS2 FC=0.439, and 99.8% and 100% of positional ITS2 keys decreased in the two trajectories.

Because these values are expressed relative to matched mock samples, constitutive excision and degradation of ITS2 during normal pre-rRNA processing cannot by itself explain the emerging divergence. Instead, the selective fall in ITS2 beginning at 2–3 hpi indicates that HSV-1 alters the production, processing, stability, or subsequent fate of ITS2-containing RNA relative to uninfected cells. The most economical interpretation is high early precursor supply followed by accelerated spacer-specific processing, deployment, or turnover; broad late 45S deterioration is a second phase. Two additional datasets with KMB17 (Shi et al., 2018) and HFF-1/MRC-5 (GSE124118) independently show that HSV-1 can also disproportionally expand ITS2-containing RNA pools. KMB17 ITS2 increased 1.589× by depth and 1.532× by fragment counting; Bayesian median 1.644× (95% CrI 0.828–3.251; P[FC>1]=0.943), all 9 Leave-One-Out (LOO) contrasts >1, and all 12 ITS2 windows increased. HFF-1 in GSE124118 increased ∼2.72× at 4 h and ∼2.26× at 8 h by depth, with Bayesian medians 2.74× and 2.28×. MRC-5 was delayed – near baseline at 4 h but ∼1.54× by 8 h (**Figure 6H**).

The analysis coupled with subcellular fractionation resolves ITS accumulation and attrition into different physical pools. At 8 hpi, alignment-depth ITS2/45S increased in every WT/mock fraction-replicate comparison: geometric relative FCs were 2.664 in Total, 1.740 in Cytoplasmic, 1.963 in Nucleoplasmic, and 1.378 in Chromatin RNA **(SI, Table S28)**. Yet direct raw-FASTQ ITS2/ITS1 decreased in seven of eight comparisons and was most depleted in Cytoplasmic (GM 0.291; ∼71% relative loss) and Nucleoplasmic RNA (0.390; ∼61% loss), while Chromatin was comparatively preserved (0.918). A hierarchical Bayesian analysis treating biological replicates as the inferential units strongly supported this compartment-specific discordance. Posterior probabilities of ITS2/ITS1 depletion were 0.9995 in Cytoplasmic and 0.9988 in Nucleoplasmic RNA, while the corresponding probabilities of ITS2/45S enrichment were 0.9943 and 0.9962.

Consequently, the joint posterior probability that HSV-1 infection simultaneously enriches ITS2 relative to 45S while depleting it relative to ITS1 was 0.9938 in Cytoplasmic and 0.9951 in Nucleoplasmic RNA (**Figure 6I**). ITS2/ITS1 depletion was also stronger in these fractions than in Chromatin with posterior probabilities of 0.9853 and 0.9746, respectively. These results provide quantitative evidence that ITS2 can be enriched relative to the broad 45S precursor while being selectively depleted relative to another spacer, consistent with compartment-specific redistribution and/or processing rather than a scalar change in rDNA output.

Under an active ITS-deployment scenario, removing a viral factor that promotes ITS consumption, destabilization, or trafficking should reverse WT depletion into accumulation. We therefore asked whether deletion of the HSV-1 virion host shutoff factor (*vhs*) reversed any of the ITS2 changes observed during WT infection. A prominent example occurred within ITS2 positions corresponding to 45S coordinates 7266–7504. In Total RNA, the 7407–7437 interval **(SI, Table S29)** was depleted during WT infection relative to mock (mean log₂FC = −0.692), whereas the overlapping 7416–7446 interval accumulated strongly in Δvhs relative to WT (+1.870). A second strongly WT-depleted interval, 7529–7559 (−0.997), likewise overlapped a Δvhs/WT accumulation signal at 7531–7561. Across the broader 7266–7504 region, all 36 replicate-consistent Total WT/mock windows were depleted (median log₂FC = −0.723), whereas all 269 replicate-consistent Total Δvhs/WT windows increased (median = +1.588). Cytoplasmic RNA showed similarly pervasive accumulation following vhs deletion, with all 240 replicate-consistent windows increasing relative to WT (median = +1.458). Because these sliding 31-nt windows overlap and are therefore not independent observations, we tested the intervention effect using a fixed, non-overlapping 31-nt bin at 7415–7445, with biological replicates as the inferential units. The same sign reversal was reproduced in both compartments: Total RNA changed from −0.829 during WT infection to +1.864 in Δvhs relative to WT, whereas Cytoplasmic RNA changed from −0.334 to +1.586. Bayesian analysis directly testing this WT-to-Δvhs reversal estimated an intervention effect of +2.48 log₂ units in Total RNA (95% CrI, +0.21 to +3.65; P[reversal > 0] = 0.982) and +1.72 log₂ units in Cytoplasmic RNA (95% CrI, −0.35 to +3.06; P[reversal > 0] = 0.959) (**Figure 6J**). The direction was retained under substantially broader priors (P = 0.959 and 0.920, respectively). Thus, deletion of *vhs* reproducibly converted depletion of this ITS2 interval during WT infection toward accumulation, providing direct intervention-based support for a dynamic relationship between *vhs* activity and the abundance or processing of specific ITS2-derived species.

#### Viral-ITS sequence similarity identifies response-prone ITS regions

Similar to MHV68 and CMVs, HSV-1 did not remodel the ribosomal spacers uniformly. For example, the small-RNA experiment with primary NHDFs infected with HSV-1 KOS or mock treated independently recovered the same central principle: viral similarity predicted where the fragmentome responded more strongly than it predicted a uniform direction of response. Across ITS1+ITS2, KOS similarity was associated with response magnitude **(SI, Table S30)** (λ=0.13884, spatial P=0.00310). To ask whether the affected positions carried a sequence relationship to the virus, we compared the infection response at every ITS nucleotide with a sequence-only HSV-1 similarity landscape across all available RNAseq datasets. Rather than defining homology by a single word length, we scanned ITS1 and ITS2 against the complete strain-17 genome at six scales (k=11, 15, 21, 31, 51, and 75 nt), projected the best-match score for each overlapping k-mer back onto the nucleotides it covered, standardized each scale, and averaged the six landscapes. The resulting composite score, σᵢ, therefore measures how consistently each host ITS position resembles HSV-1 across short to intermediate sequence scales:

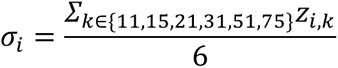

We then compared this fixed sequence landscape with the independently measured positional infection response, Δᵢ. Two correlations distinguish effects that would otherwise be conflated: β tests whether HSV-like sequence predicts the direction of the response **(SI, Table S31)**, whereas λ tests whether it predicts the magnitude of the response regardless of sign:

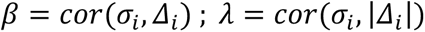

Significance was evaluated by 10,000 independent circular shifts of the intact similarity landscape within ITS1 and ITS2, preserving each spacer’s sequence structure and autocorrelation while breaking its true alignment with the RNA response. Thus, the test asks a direct spatial question: does HSV-like host sequence fall on the positions that actually respond during infection more often than expected from other registrations of the same ITS landscape? The earliest strong signal appeared before broad ITS2 loss was established. At 1–2 hpi in newly synthesized 4sU RNA, HSV similarity predicted response magnitude across ITS1+ITS2 (λ=0.3003, spatial P=0.0084), with a clear ITS2 component (λ=0.2829, P=0.0149) (**Figure 7A; SI, Table S32**). At this point the informative feature was not whether a position increased or decreased, but whether it responded at all. Only subsequently did the bulk phenotype emerge: beginning at 2–3 hpi, newly synthesized ITS2 progressively separated from the contemporaneous 45S pool, reaching roughly half of its mock-normalized expectation by 3–5 hpi. The sequence signal therefore preceded the broad spacer-wide attrition.

**Figure 7.**
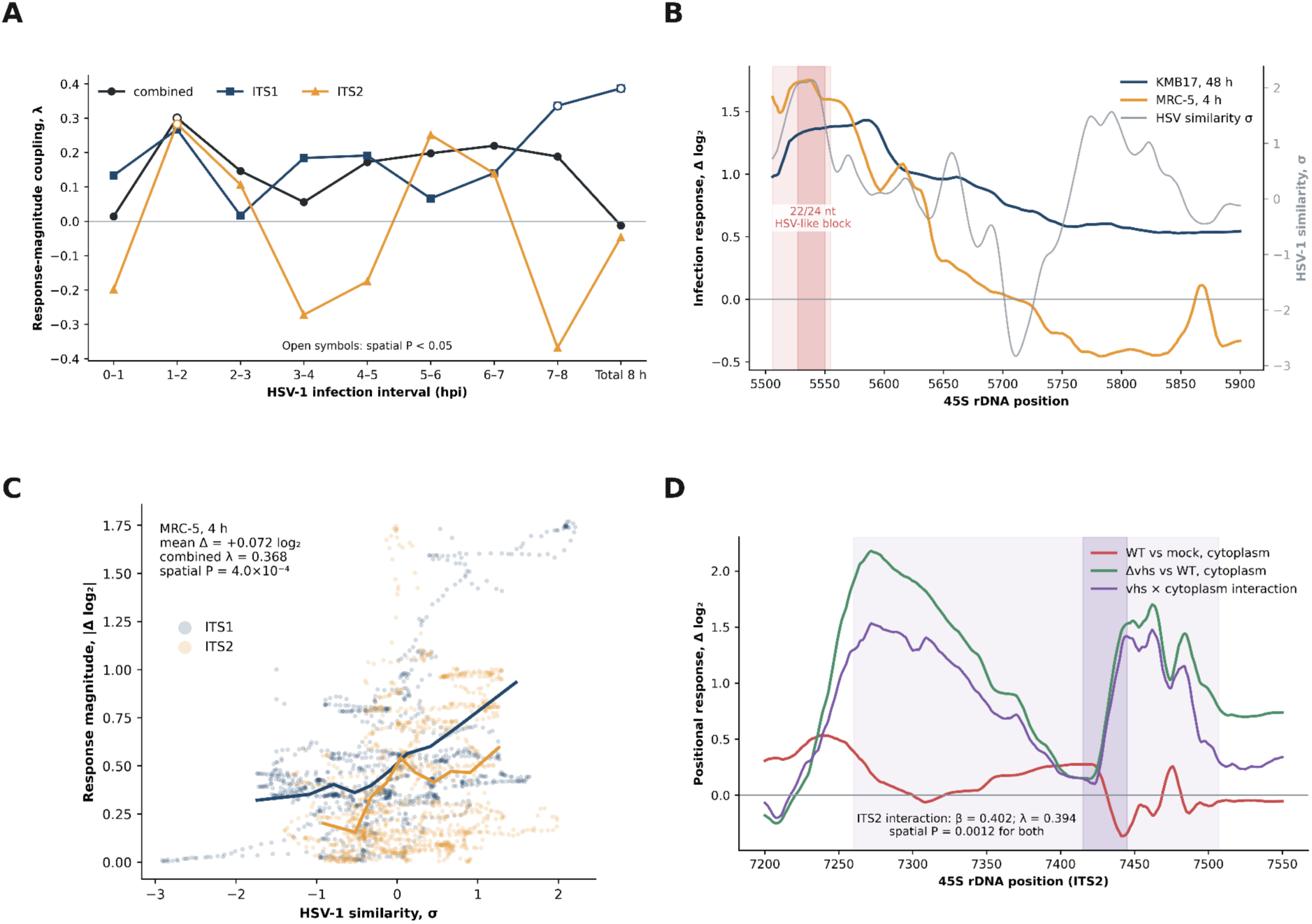
Sequence-dependent remodeling of ITS RNA during HSV-1 infection. (A) Temporal emergence of sequence-dependent ITS responses during HSV-1 infection. Spatial coupling between HSV-1 sequence similarity and the magnitude of host ribosomal internal transcribed spacer (ITS) RNA responses across successive intervals of infection. Curves show the response-magnitude correlation (\lambda) for ITS1, ITS2, and the combined ITS1 + ITS2 regions. Viral sequence similarity was quantified using a multiscale local homology score (\sigma), integrating matches of 11, 15, 21, 31, 51, and 75 nucleotides between HSV-1 and human ITS sequences. At each time point, \lambda was calculated as the Pearson correlation between \sigma and the absolute log₂ change in ITS RNA abundance relative to mock-infected controls. Open circles indicate significant spatial associations (*P*<0.05), assessed using 10,000 within-ITS circular permutations that preserve the spatial autocorrelation structure of the sequence-similarity and RNA-response profiles. Positive \lambda values indicate that ITS positions with greater viral sequence similarity exhibit larger transcriptional responses, irrespective of whether RNA abundance increases or decreases. The early emergence of positive spatial coupling, preceding more extensive ITS remodeling at later infection stages, is consistent with viral sequence similarity identifying regions particularly susceptible to HSV-1-associated RNA perturbation. **(B) A recurrent 5′ ITS1 response domain coincides with a region of high HSV-1 sequence similarity.** Spatial profiles of infection-associated ITS1 RNA responses in two independent HSV-1 infection datasets: KMB17 cells at 48 hours post-infection (blue; GSE103763) and MRC-5 fibroblasts at 4 hours post-infection (orange; GSE124118). RNA responses are expressed as per-nucleotide log₂ fold changes relative to mock-infected controls (left axis). The gray curve represents the multiscale HSV-1 sequence-similarity score (σ; right axis), calculated from local sequence matches of 11, 15, 21, 31, 51, and 75 nucleotides. The lightly shaded region (positions 5506–5555 of the human 45S rDNA reference) identifies a recurrent 5′ ITS1 response domain, while the darker shading highlights an embedded 24-nucleotide segment (positions 5527–5550) exhibiting 22/24 nucleotide identity to HSV-1 strain 17. Curves were smoothed using a centered 9-nucleotide moving average for visualization. The convergence of infection-associated RNA responses and viral sequence similarity within this region, despite differences in cell type and infection duration, identifies the 5′ ITS1 domain as a candidate hotspot of HSV-1-associated RNA remodeling. **(C) HSV-1 sequence similarity predicts the magnitude of ITS RNA remodeling despite minimal net changes in abundance.** Relationship between local HSV-1 sequence similarity (σ) and the absolute magnitude of infection-associated RNA responses (|Δ log₂|) across ITS1 (blue) and ITS2 (orange) in MRC-5 fibroblasts at 4 hours post-infection (GSE124118). Each point represents an individual nucleotide position within the ITS regions, with RNA responses expressed as log₂ fold changes relative to mock-infected controls. Solid curves show mean absolute responses within deciles of the sequence-similarity distribution for each ITS region. Despite a small overall mean change in ITS abundance (+0.072 log₂), response magnitude was strongly associated with viral sequence similarity (combined ITS1 + ITS2, λ = 0.368; spatial *P* = 4.0 × 10⁻⁴). The correlation coefficient λ was calculated as the Pearson correlation between local sequence similarity and absolute RNA response magnitude. Statistical significance was assessed using 10,000 within-ITS circular permutations, preserving the spatial structure of the underlying profiles. These results reveal substantial, sequence-associated positional remodeling of ITS RNA that is largely obscured by conventional analyses of mean transcript abundance. (D) *vhs*-dependent remodeling of a recurrent ITS2 response domain during HSV-1 infection. Positional profiles of ITS2 RNA responses in human fibroblasts infected with wild-type (WT) HSV-1 or a mutant lacking the viral host-shutoff nuclease (*Δvhs*), using subcellular fractionation RNA-seq datasets (GSE100576 and GSE129582). The red curve shows the cytoplasmic response to WT infection relative to mock-infected controls, whereas the green curve represents the difference between *Δvhs* and WT infection in the cytoplasmic fraction. The purple curve shows the *vhs* × cytoplasm interaction, calculated as the difference between the *Δvhs*–WT contrasts in cytoplasmic and chromatin-associated RNA. Responses are expressed as per-nucleotide log₂ abundance differences and smoothed using a centered 9-nucleotide moving average for visualization. Light purple shading identifies a recurrent ITS2 response domain (45S rDNA positions 7260–7507), with darker shading highlighting the 7415–7445 interval examined in the intervention analysis. Within ITS2, the compartment-dependent *vhs* effect was associated with local HSV-1 sequence similarity, measured using a multiscale homology score (σ) integrating sequence matches of 11, 15, 21, 31, 51, and 75 nucleotides. Both the directional association (β = 0.402) and response-magnitude association (λ = 0.394) were significant (spatial *P* = 0.0012 for each), as determined by 10,000 within-ITS circular permutations preserving the spatial autocorrelation of the sequence-similarity landscape. The profiles illustrate how loss of *vhs* alters the abundance and subcellular distribution of RNA originating from a recurrent ITS2 response domain.

The sign of the relationship then changed as infection progressed, while its spatial organization persisted. At 5–6 hpi, higher HSV similarity predicted a more positive ITS2 response (β=0.4411, P=0.0329), and the relationship remained positive at 6–7 hpi (β=0.4034, P=0.0163). By 7–8 hpi the dominant signal had shifted to ITS1 and reversed direction (β=−0.3356, P=0.0240), while HSV similarity still predicted response magnitude (λ=0.3358, P=0.0248). Total RNA at 8 h reproduced the same ITS1 pattern (β=−0.3866, P=0.0415; λ=0.3866, P=0.0442). Thus, viral similarity predicted where the response was concentrated more consistently than it predicted whether RNA at those positions would accumulate or disappear. Independent HSV-1 infections reproduced this architecture. At 48 hpi in KMB17 fibroblasts, ITS1 showed both strong directional and magnitude coupling (β=0.3856, P=0.0021; λ=0.3808, P=0.0106). The strongest localization was the 5′ ITS1 interval at rDNA positions 5506–5555, which contains a 24-nt segment with 22/24 identities to strain 17 and recurred as a major response region in an independent fibroblast experiment (**Figure 7B; SI, Table S33**). Yet the signal was not simply the consequence of this short match: removing the 24-nt block left positive β and λ, and even excluding a ±200-nt neighborhood retained a positive residual relationship. The viral-like segment therefore sits within a broader response-prone ITS1 domain.

The MRC-5 4-h infection provided an especially revealing test because bulk ITS abundance barely changed. Across ITS1+ITS2 the mean positional response was only +0.072 log2, yet HSV similarity strongly predicted where remodeling occurred (λ=0.3679, P=0.00040), with signals in both ITS1 (λ=0.5149, P=0.0081) and ITS2 (λ=0.2841, P=0.0097) (**Figure 7C**). A second recurrent domain in ITS2, spanning approximately rDNA positions 7260–7507, became mechanistically informative in the WT/Δ*vhs* fractionation experiment. Deletion of the HSV-1 host-shutoff nuclease changed the cytoplasmic sequence-response relationship relative to chromatin: the combined-spacer *vhs*×cytoplasm interaction was strongly associated with HSV similarity (β=0.3992, P=0.0018), and ITS2 showed both directional and magnitude effects (β=0.4019 and λ=0.3938; P=0.0012 for each) (**Figure 7D**). The analogous nucleoplasmic interaction was not significant. Within the recurrent ITS2 domain, WT infection produced broad depletion whereas Δ*vhs* shifted the same region toward accumulation. At the fixed 7415–7445 interval, the WT-to-Δ*vhs* intervention effect was +2.48 log2 units in total RNA (95% CrI +0.21 to +3.65; P[reversal>0]=0.982) and +1.72 log2 units in cytoplasmic RNA (95% CrI −0.35 to +3.06; P[reversal>0]=0.959). Thus, a viral RNA-destabilizing function acts directly on the fate of one of the sequence-defined ITS response domains.

Remarkably, these results reveal a reproducible spatial architecture beneath the changing bulk ITS phenotype. HSV-like sequence identifies ITS regions with an increased propensity to respond, but it does not prescribe a single direction of response. The same sequence-defined regions can be retained, enriched, or depleted depending on infection stage, cellular context, and *vhs* activity. This pattern is difficult to reduce to uniform changes in 45S transcription or ordinary spacer turnover alone: the response is regional, appears before broad ITS2 attrition, recurs across independent infections, and is altered by a defined viral RNA-processing function. These observations raised a more fundamental question. Does the changing ITS pool simply record virus-driven remodeling of pre-rRNA processing and stability, or does infection move ITS RNA through a distinct host response state in which loss from the measurable precursor pool can accompany biological deployment? To distinguish these alternatives, we next modeled the HSV-1 time course as competing dynamical hypotheses ranging from passive ribosome biogenesis to virus-driven remodeling and infection-responsive host ITS deployment.

### 3. Dynamic modeling identifies host ITS deployment as the parsimonious explanation for HSV-1-induced ribosomal spacer remodeling

The central modeling problem here is not whether HSV-1 perturbs ribosome biology; that is biologically expected. The more discriminating question is whether the unusual behavior of ITS1 and ITS2 can be explained as a passive consequence of precursor-rRNA remodeling, or whether the data require a distinct host ITS state whose production, loss, trafficking, or deployment is regulated during infection. This distinction matters because abundance alone is not equivalent to biological activity. A pool can decrease precisely because material is being transferred out of it. Accordingly, a fall in measurable ITS2 is compatible with accelerated degradation, but it is also compatible with accelerated flux from a measurable precursor/intermediate pool into a short-lived deployed state.

The HSV-1 4sU dataset provided uniquely useful information because it reports newly synthesized RNA over successive 1-h intervals rather than only steady-state abundance. Across eight intervals from 0-1 to 7-8 hpi, total nascent 45S output remained near baseline through the period in which ITS2 fell sharply. At 3-4 hpi, for example, total 45S was approximately 1.03-fold relative to baseline whereas ITS2 was approximately 0.54-fold. The relative ITS2/45S signal remained approximately 0.52-0.64 through much of 3-8 hpi. By 7-8 hpi, ITS2 was approximately 0.46-fold whereas total 45S remained approximately 0.82-fold. ITS1 also declined, but weaklier, producing progressive ITS2/ITS1 decoupling. These observations motivated a model in which the measurable ITS pool is governed by both influx from 45S processing and outflux into degradation or a functionally distinct state (**Figure 8A**).

**Figure 8.**
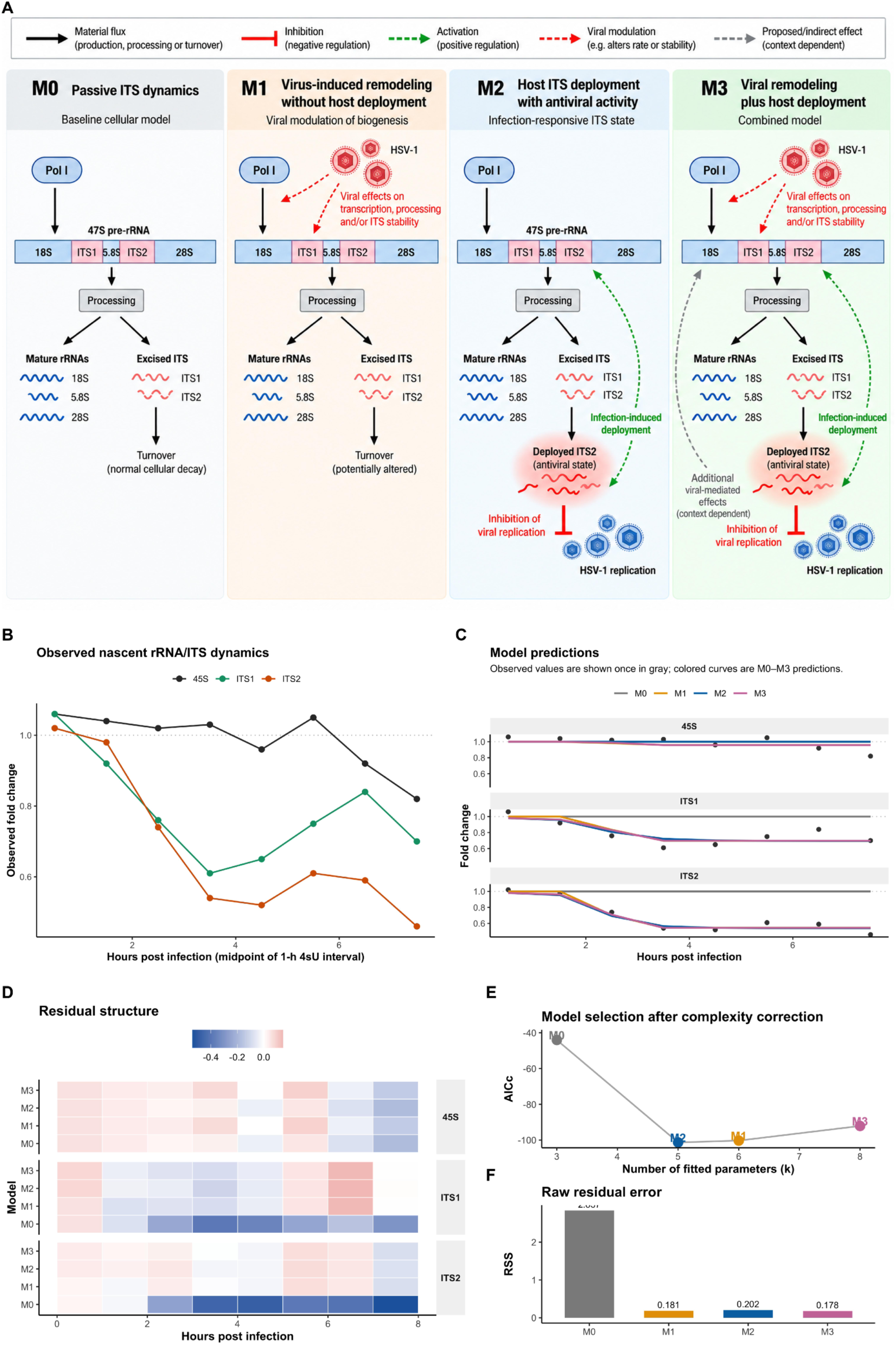
Temporal dynamics of rRNA and ITS transcripts during HSV-1 infection and comparison of mechanistic models. (A) Canonical dynamical hypotheses for HSV-induced ITS remodeling. Four alternative biological models describe increasingly complex explanations for ITS behavior during HSV-1 infection. M0, passive ITS dynamics, represents the baseline null model: RNA polymerase I produces the 47S precursor, mature 18S, 5.8S, and 28S rRNAs are generated by processing, and excised ITS1 and ITS2 undergo ordinary cellular turnover. No virus-dependent regulation or antiviral ITS state is required. M1, virus-induced remodeling without host deployment, allows HSV-1 to alter rRNA transcription, precursor processing, and/or ITS stability, but treats the resulting ITS changes as consequences of infection rather than an antiviral host response. M2, host ITS deployment with antiviral activity, introduces an infection-responsive transition of excised ITS RNA, represented here by ITS2, into a distinct deployed state capable of inhibiting viral replication. Thus, ITS abundance need not increase for ITS antiviral activity to increase. M3, viral remodeling plus host deployment, combines the mechanisms represented by M1 and M2: HSV can remodel rRNA/ITS metabolism while the host simultaneously deploys ITS-derived RNA into an antiviral state. Black arrows indicate material production, processing, or turnover; red inhibitory symbols indicate negative regulation of viral replication; green dashed arrows denote activation of ITS deployment; and red dashed arrows denote virus-dependent modulation of cellular RNA metabolism. M0–M3 are alternative biological hypotheses rather than sequential analysis stages. Importantly, M3 is biologically the most inclusive, and arguably the most intuitively plausible, hypothesis, but its additional mechanisms are justified statistically only if they explain information not already captured by the simpler models. **(B) Nascent rRNA and ITS dynamics during HSV infection and comparison of alternative dynamical models.** Observed nascent 45S, ITS1, and ITS2 fold changes across the HSV-1 infection time course. Whereas nascent 45S production remains comparatively preserved during much of infection, ITS1 and particularly ITS2 progressively decline, indicating that ITS abundance does not simply track precursor-rRNA production. **(C) Predictions from the four canonical dynamical models are compared against the same observed trajectories** (gray points). M0 represents passive rRNA biogenesis and ITS turnover; M1 allows HSV-dependent remodeling of rRNA/ITS metabolism without an antiviral ITS state; M2 introduces infection-responsive deployment of ITS RNA into an antiviral state; and M3 combines viral remodeling with host ITS deployment. The models differ in mechanism rather than in the observations to which they are fitted. **(D) Residuals (observed minus predicted values) reveal the structure of model misfit across RNA species and infection time.** The large, systematic deviations under M0 show that passive production and turnover alone cannot reproduce the observed RNA dynamics, whereas infection-responsive models substantially reduce these discrepancies. **(E) Model selection by the small-sample corrected Akaike information criterion (AICc)** accounts for both goodness of fit and the number of fitted parameters. M2 provides the lowest AICc despite having fewer parameters than M3, indicating that the additional mechanisms in M3 do not provide sufficient improvement in fit to compensate for their added complexity. **(F) Raw residual sum of squares (RSS)** is shown separately to distinguish descriptive fit from complexity-corrected model selection. M3 achieves the smallest RSS, as expected for the most flexible model, but improves only modestly over M2; by contrast, the large reduction from M0 to the infection-responsive models demonstrates that an infection-dependent process is required to explain the RNA trajectories. Overall, M2 stands out as the most parsimonious explanation of the RNA dynamics while showing explicitly that its selection does not result from having the smallest raw residual error.

Consequently, four biological hypotheses were encoded as nested dynamical models. We defined four biological models, M0-M3 (Figure **8A****; SI, Table S34**). These names refer only to biological hypotheses; subsequent stages such as identifiability analysis, Bayesian robustness, fractionation analysis, or functional integration are analysis stages rather than additional biological models.

#### M0, passive ribosome-biogenesis null

M0 assumes that ITS behavior is an ordinary consequence of 45S transcription, processing, and turnover. There is no host ITS antiviral state and no explicit virus-directed remodeling term. In its simplest form,

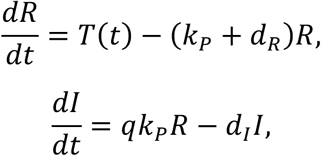

where *R* denotes the precursor-rRNA pool, *I* the measurable ITS precursor/intermediate pool, *T*(*t*) precursor transcription, *k*, processing from precursor into ITS-containing intermediates, *q* the effective yield into the measured ITS pool, and *d*_-_ and *d*_._ loss terms. Virus abundance *V* can change with time, but it does not alter ITS kinetics in M0.

#### M1, viral remodeling without host ITS function

M1 gives HSV substantial freedom to alter ribosomal transcription, processing, or ITS stability, but it still denies an antiviral ITS state. A convenient representation is

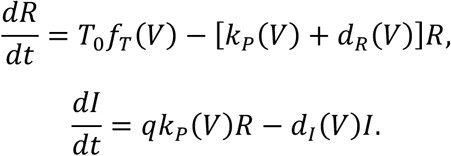

Virus dependence was represented by smooth bounded response functions, for example

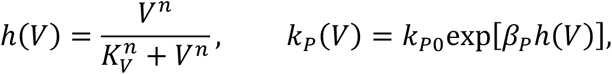

with analogous terms available for transcription or decay. Thus, M1 is a deliberately generous nonfunctional alternative: it can reproduce ITS loss by allowing HSV to remodel the machinery that creates or destroys ITS-containing RNA.

#### M2, host ITS deployment

M2 introduces a distinct deployed state *D*. The measurable ITS pool can therefore decrease because ITS material leaves that pool, while the deployed state can increase or turn over rapidly:

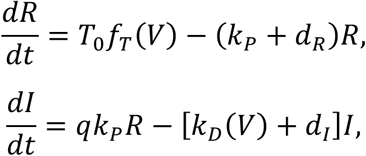

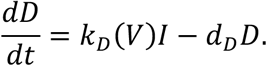

The infection-responsive deployment rate was represented as

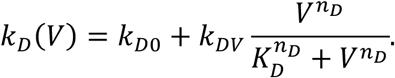

For functional experiments, the deployed state was allowed to inhibit viral expansion:

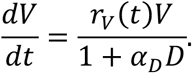

The key quantity is therefore not only standing ITS abundance but deployment flux,

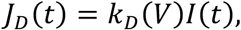

and its cumulative activity,

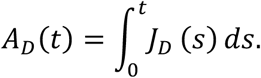

The intuition is analogous to a reservoir with an inlet and an outlet. A declining water level does not by itself show that the inlet has shut down; if the inlet remains active while the level falls, the outlet must have increased. The 4sU experiment provides information about the inlet. Maintained nascent 45S production together with declining ITS2 therefore constrains ITS2 outflux to exceed its replenishment during the critical phase of infection.

#### M3, hybrid viral remodeling plus host deployment

M3 combines the mechanisms of M1 and M2:

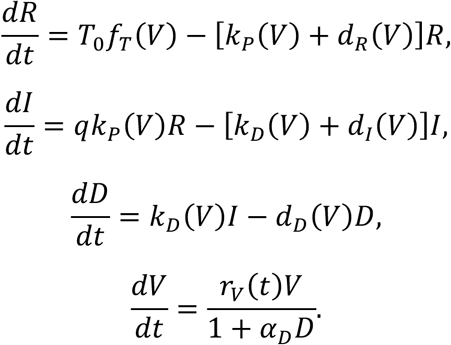

Biologically, M3 may initially appear the most realistic model because HSV unquestionably remodels host RNA metabolism. Statistically, however, the relevant question is not whether viral remodeling occurs somewhere in the infected cell. It is whether explicit remodeling parameters are needed to explain the distinctive ITS observations after host deployment has already been included. M3 was therefore retained as a strong, deliberately permissive competitor rather than excluded as an obvious biological truth.

### Simulated RNA trajectories reveal an intrinsic ambiguity between remodeling and deployment

The first formal comparison used 24 log-scale observations representing total 45S, ITS1, and ITS2 behavior across the HSV-1 temporal series (**Figure 8B**). Viral-feedback parameters were fixed at zero at this stage because RNA trajectories alone cannot identify an ITS-to-virus causal effect; those parameters were reserved for the later perturbation analysis. Baseline kinetic constants were constrained in the normalized ODE formulation so that model comparison focused on infection-induced changes rather than unconstrained rescaling. All three infection-responsive models dramatically improved fit over M0, reproducing the principal temporal features of the observed RNA trajectories (**Figure 8C; SI, Table S35**). Examination of the residual structure further illustrated the improved fits of M1, M2, and M3 relative to M0 (**Figure 8D**). M0 produced RSS = 2.837 and AICc = −44.05. M1 reduced RSS to 0.1815 and M2 to 0.2022, whereas M3 achieved the smallest RSS, 0.1779, as expected for the most flexible model. However, accounting for model complexity shifted the preference to M2 (**Figure 8E**). M2 used five fitted parameters and achieved AICc = −101.30, the best of the four models. M1 used six parameters and had AICc = −100.29 (ΔAICc = 1.01), whereas M3 required eight parameters and fell to AICc = −92.11 (ΔAICc = 9.19). Akaike weights were 0.619 for M2, 0.374 for M1, 0.0063 for M3, and effectively zero for M0. The contrast between raw residual error and complexity-adjusted model selection is particularly informative (**Figure 8F**). Although M3 achieved the smallest RSS, its additional mechanisms reduced residual error by only 0.0036 relative to M1 and 0.0243 relative to M2. These modest improvements did not compensate for the additional fitted parameters. Thus, once an infection-responsive ITS deployment route is included, explicit viral remodeling of several additional kinetic terms contributes little explanatory information.

This does not imply that HSV-1 fails to remodel ribosome biology. Rather, the characteristic ITS1/ITS2 trajectories can be explained without invoking the additional remodeling mechanisms included in M3. The distinction is central to interpreting the biological differences among the models.

### Hessian geometry reveals why the apparent flexibility of M3 is not equivalent to information

M3 is the most biologically elaborate model: it proposes that HSV actively remodels rRNA/ITS metabolism while the host simultaneously deploys ITS-derived RNA into an antiviral state.

Adding both processes gives M3 more parameters and therefore more ways to reproduce the observed RNA trajectories. However, a more elaborate model is informative only if the data can determine how those additional processes behave. We therefore asked a simple question: if the parameters are moved away from their best-fitting values, how quickly does the model cease to fit the data? If even large changes in a parameter, or in a combination of parameters, leave the predicted RNA trajectories almost unchanged, the experiment cannot determine that parameter reliably. We thus examined local curvature of the objective function around each optimum using the Hessian matrix. The Hessian is the matrix of second derivatives of the objective with respect to parameters,

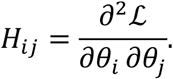

A large positive Hessian eigenvalue corresponds to a direction in parameter space in which the fit worsens rapidly when parameters are moved away from the optimum; that combination of parameters is well constrained. An eigenvalue near zero corresponds to a nearly flat direction: many parameter combinations produce almost indistinguishable trajectories. A negative eigenvalue at a numerical optimum can indicate a saddle-like or poorly resolved local geometry, numerical approximation error, or a boundary/near-boundary problem rather than a clean, isolated minimum. M1 and M3 showed small negative minimum Hessian eigenvalues (approximately -3.05 x 10^-4 in the normalized ODE fit), and M3 had an extreme Hessian condition measure, consistent with substantial parameter sloppiness. M2 also contained a flat direction in the RNA-only stage, which is unsurprising because the causal ITS-to-virus feedback coefficient cannot be learned from RNA measurements alone. The important contrast is that M3 adds several poorly constrained directions without materially improving the trajectory. Thus, its apparent mechanistic richness is partly unidentified flexibility (**Figure 9A)**.

**Figure 9.**
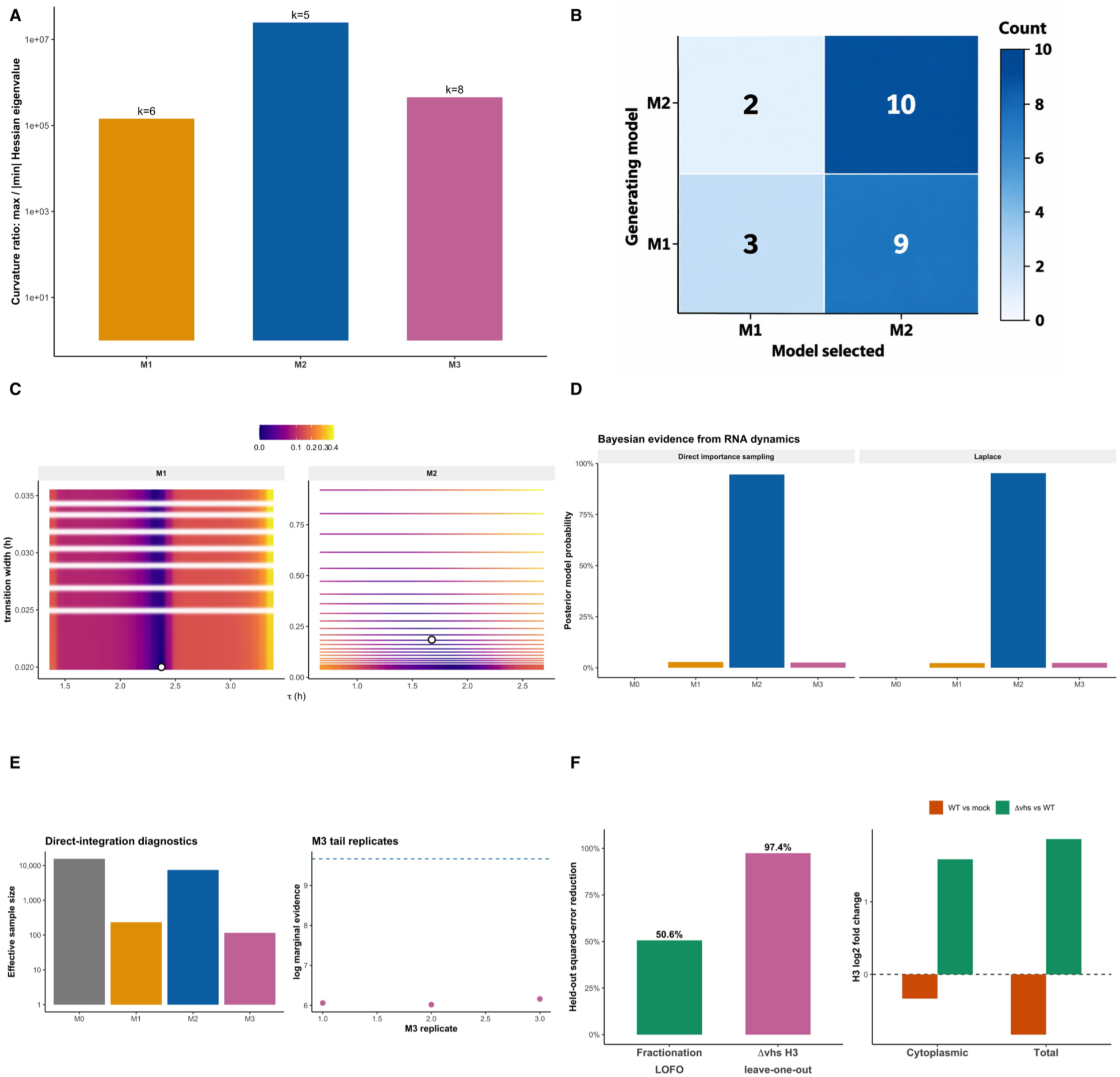
**Dynamical model discrimination and independent tests of ITS regulatory structure during HSV infection**. **(A) Local parameter geometry of the RNAseq-only dynamical models.** Local curvature of the fitted likelihood surface was evaluated for M1–M3 using the ratio of the largest to the absolute smallest Hessian eigenvalue. Larger ratios indicate greater disparity among parameter-space directions: some combinations of parameters are tightly constrained by the RNA trajectories, whereas others can change substantially with comparatively little effect on model fit. M1, M2, and M3 contain 6, 5, and 8 fitted parameters, respectively (values above bars). All three models show appreciable anisotropy, with M2 displaying the largest curvature ratio. Importantly, these ratios are diagnostics of local parameter geometry rather than measures of model fit or biological support; a large ratio indicates a weakly constrained or “sloppy” parameter direction and does not by itself favor or disfavor a model. The presence of weakly constrained directions, together with problematic local curvature for M1 and M3, motivated subsequent Bayesian marginal-evidence calculations and their verification by direct importance sampling rather than reliance on local optimization alone. **(B) Synthetic-data recovery reveals limited discrimination between M1 and M2 from RNAseq dynamics alone.** Model-recovery analysis tested whether the observed RNA time-series design contains sufficient information to distinguish virus-induced RNA remodeling (M1) from host ITS deployment (M2). Twelve synthetic datasets were generated independently under each model using the fitted dynamical framework and realistic observation noise, and each dataset was then fitted and selected by AICc without using knowledge of its generating model. Numbers indicate the number of 12 replicate datasets assigned to each candidate model; darker shading denotes a larger count. Of datasets generated under M1, only 3/12 were recovered as M1, whereas 9/12 were classified as M2. Conversely, M2 was correctly recovered in 10/12 datasets, with 2/12 classified as M1. Thus, the RNA design preferentially selects M2 but does not reliably identify the mechanism that generated the data. The asymmetric recovery demonstrates an intrinsic M1–M2 identifiability limitation and parsimony bias in the RNA-only comparison, providing an important qualification to the RNA-only preference for M2 and motivating the use of independent causal perturbation evidence to distinguish remodeling from antiviral ITS deployment. **(C) Transition geometry of alternative mechanisms underlying the rRNA dynamics.** Diagnostic parameter surfaces show how the fit of M1 and M2 depends on the timing and temporal width of the infection-responsive transition. The horizontal axis gives the transition midpoint, \tau, in hours after infection, whereas the vertical axis gives the transition width, which determines how abruptly or progressively the modeled response develops. Color represents the increase in residual error (ΔRSS) relative to the best-fitting point within each model; darker regions therefore identify combinations of transition timing and width that remain close to the optimum. Open circles mark the minimum-RSS solutions. M1 favors a transition centered at approximately 2.37 h with its width driven to the lower boundary (∼0.02 h), corresponding to an effectively abrupt, step-like change in virus-dependent RNA remodeling. In contrast, M2 favors an earlier transition centered at approximately 1.68 h and a finite width of ∼0.18 h, consistent with a progressively developing ITS-deployment process. The different geometries illustrate that M1 and M2 can reproduce similar RNA abundance trajectories through mechanistically distinct temporal responses. These surfaces are fixed-nuisance diagnostic scans rather than fully reoptimized profile likelihoods and are therefore intended to visualize transition geometry and parameter constraint, not to provide an additional formal model-selection test. **(D) Bayesian model comparison of the rRNA dynamics and numerical verification of evidence.** Bayesian evidence was evaluated for the four biological models using equal prior model probabilities and the primary weakly informative specification for infection-effect parameters (SD = 1.5). Posterior model probabilities obtained by Laplace approximation (right) and by direct importance sampling (left). Both calculations strongly favor M2 under the primary prior specification, while assigning substantially less support to M1 and M3 and essentially none to the passive M0 model. Importantly, the two methods evaluate the same prior-dependent Bayesian evidence; their agreement therefore tests the numerical robustness of the marginal-likelihood calculation rather than independently validating the biological model or the choice of prior. **(E left)** Effective sample sizes (ESS) for the production importance-sampling calculation using 30,000 draws per model. M2 achieved substantially greater ESS than M1 or M3, indicating more efficient exploration of its posterior-supported parameter space, whereas the lower ESS for M3 motivated additional numerical checks. **(E right)** Independent tail-focused importance-sampling replicates for M3. Points show the estimated log marginal evidence from separate M3 calculations; the dashed horizontal line indicates the corresponding M2 estimate (log marginal evidence = 9.67), accounting for the broader displayed y-axis range. Despite variation among M3 replicates resulting from its more difficult parameter geometry, the estimates were tightly clustered around 6 and all remained well separated from M2, indicating that the primary Bayesian preference for M2 was not an artifact of the Laplace approximation or an unstable M3 evidence estimate. (F) Orthogonal analyses of non-scalar ITS behavior and a *vhs*-sensitive regional response. Independent RNA measurements were used to test features of ITS regulation that are not captured by the primary abundance time series. **(Left)** Improvement in held-out prediction when additional structure was introduced into leave-one-out analyses. Accounting for cellular fractionation reduced held-out squared error by 50.6%, indicating that ITS behavior differs substantially between cellular compartments and cannot be adequately represented by a single scalar abundance. A separate leave-one-out analysis of the H3 ITS2 region produced a 97.4% reduction in prediction error when a common Δ*vhs*-associated reversal was included, demonstrating that this regional response is highly reproducible across the available contrasts. **(Right)** Direction of the H3 response in cytoplasmic and total-RNA fractions. H3 abundance decreases during wild-type HSV-1 infection relative to mock (WT vs mock, orange), but this effect reverses in the absence of functional *vhs* (Δ*vhs* vs WT, green) in both fractions. The concordant reversal indicates that the localized H3 response depends strongly on the viral *vhs* context rather than simply tracking global ITS2 abundance. Together, these orthogonal analyses support a view of ITS2 as a spatially and regionally structured RNA system, while not by themselves establishing whether the *vhs*-sensitive behavior reflects direct *vhs* action or an indirect *vhs*-dependent regulatory process.

### M1 remains the viable RNA-only alternative, and synthetic recovery shows why

Although M2 decisively outperformed M0 and strongly outperformed M3 by AICc, M1 remained within approximately one AICc unit. We therefore explicitly tested whether the available eight-interval RNA design can reliably discriminate viral remodeling from host deployment. Synthetic datasets were generated under fitted M1 and M2 models, noise was added at the observed scale, and both candidate models were refitted without revealing the generating model. Recovery was imperfect and asymmetric. Of 12 M1-generated datasets, only three were selected as M1 whereas nine were selected as M2. Of 12 M2-generated datasets, 10 were correctly selected as M2 and two as M1. This experiment is important because it prevents overinterpretation of the frequentist RNA-only result. The design contains enough information to reject the passive M0 model and to disfavor unnecessary M3 complexity, but it does not guarantee mechanistic discrimination between M1 and M2 from trajectory shape alone. In particular, the relative parsimony of M2 can cause it to win some datasets actually generated by M1. Thus, RNA dynamics favored M2 as the minimal explanation for the observed ITS2 response, even though did not fully distinguish host ITS deployment from virus-driven remodeling. This residual M1–M2 ambiguity reflected a fundamental limitation of the RNA measurements: both mechanisms can generate loss of measurable ITS2 through different underlying fluxes. Resolving these alternatives therefore required information beyond the RNA abundance trajectories themselves (**Figure 9B**).

### Remodeling and deployment predict distinct transition dynamics

We next examined the timing and width of the infection-responsive transition. In the fitted M1 solution, the transition was centered at approximately 2.37 hpi and its width collapsed toward the lower numerical boundary, producing an almost step-like change. In M2, the transition was earlier (approximately 1.68 hpi) and had a finite width of approximately 0.185 h, yielding a progressive change rather than a numerical switch. A fixed-nuisance tau-width surface supported this contrast (**Figure 9C**). This analysis was treated as transition-geometry diagnostics rather than a new biological model or a formal profile likelihood, because nuisance parameters were not fully reoptimized at every grid point. Nevertheless, the geometry is biologically suggestive. M1 explains the trajectory most efficiently by invoking an abrupt virus-driven kinetic switch, whereas M2 can explain it through a finite-width rise in deployment. This observation strengthens the mechanistic coherence of M2 but was not used as decisive evidence by itself.

### Bayesian model evidence asks a different question from AICc

AICc compares penalized maximum fit. Bayesian model evidence instead averages the likelihood over the entire parameter space allowed by the prior:

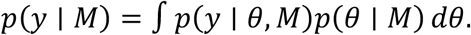

Posterior model probabilities follow from Bayes’ rule,

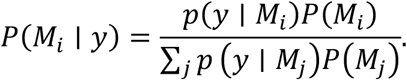

This distinction is especially important for M3. A highly flexible model can contain a parameter combination that fits very well, yet still have low marginal evidence if most of its allowed parameter space fits poorly. Bayesian evidence therefore implements an automatic complexity penalty through integration rather than simply counting parameters. We first approximated the evidence with a Laplace expansion around the posterior mode. If *θ* is the posterior mode and *H* is the Hessian of the negative log posterior at that mode, the approximation is

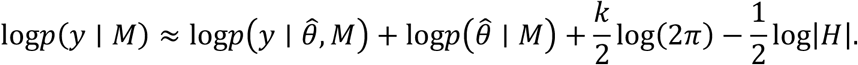

Intuitively, the first term rewards a high posterior peak, whereas the Hessian determinant measures how broad or narrow the supported parameter region is. A model whose good fit exists only in a narrow or awkwardly curved region receives less integrated evidence than a model with a broad region of good explanations. Under the primary proper priors, the Laplace calculation yielded posterior probabilities of 0.9527 for M2, 0.0232 for M1, 0.0241 for M3, and approximately 3.1 x 10^-9 for M0 (). The corresponding Bayes factor was approximately 41:1 for M2 over M1 and approximately 39:1 for M2 over M3. This was a striking strengthening of the M2 result relative to AICc, but we did not accept the Laplace calculation uncritically because M1 and M3 had already shown nonideal curvature in the frequentist fits.

### Direct importance sampling independently corroborates the primary Bayesian result

To determine whether the Laplace result was an artifact of local Gaussian approximation, we performed direct posterior-centered importance sampling. For draws *θ*_6_ from proposal density *q*(*θ*), marginal evidence is estimated from importance weights

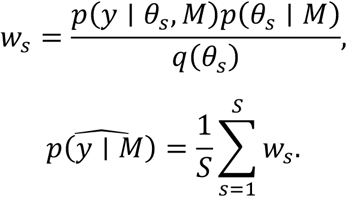

The effective sample size,

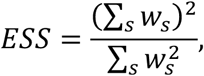

was used to distinguish stable evidence estimates from calculations dominated by a handful of extreme weights. With 30,000 draws per model under the primary effect-prior scale (SD = 1.5), direct importance sampling produced log evidences of -10.282 for M0, 6.158 for M1, 9.668 for M2, and 6.057 for M3. Posterior model probabilities **(SI, Table S36)** were therefore 2.1 x 10^-9^, 0.0283, 0.9461, and 0.0256, respectively. M2 had ESS = 7,458 and a Monte Carlo standard error of only 0.0097 log units. M3 was harder to integrate because of its tail geometry (ESS = 115), so three additional independent 30,000-draw M3 tail replicates were performed. Their log evidences were 6.060, 6.019, and 6.160. All remained more than 3.5 log-evidence units below M2. Thus, the central M2-over-M3 conclusion did not depend on the Laplace approximation (Figure **9D-E**). Prior sensitivity was informative rather than ignored. A tighter effect prior (SD = 1.0) distributed evidence much more evenly among M1-M3, and the M3 importance estimate had very low ESS, preventing a reliable exact ranking. Broader SD = 2.5 priors strongly favored M2. Prior-predictive simulation showed that broader M1/M3 priors also generated increasingly extreme fold changes. We therefore report the prespecified SD = 1.5 analysis as primary and the sensitivity analyses as evidence that the strength of the RNA-only model probability is prior dependent. The robust conclusion becomes narrower: M0 is untenable; M2 is strongly supported under the primary defensible prior; and M3’s extra remodeling structure is not rescued by a more reliable direct integration method.

#### vhs experimental perturbation reveals a virus-dependent, regionally structured ITS2 response

Additional HSV datasets were used as structurally critical tests rather than as extra degrees of freedom in the primary 24-observation fit. Fractionation data showed that ITS behavior could not be represented as a single scalar change shared across compartments. In an RNA-only integration containing temporal, fractionation, and *vhs*-perturbation observations, allowing assay-specific fractionation structure reduced leave-one-fraction-out squared error by approximately 50.6% and RMSE by approximately 29.7%. A common regional response to deletion of the HSV virion host shutoff factor (*vhs*) reduced held-out error by approximately 97.4% for the tested H3 region. H3 showed reproducible WT depletion and strong accumulation/reversal in Delta-*vhs* infection, with replicate reversal magnitudes of approximately 2.69 and 1.92 log2 units. These observations do not independently prove M2, because *vhs* could influence production, degradation, trafficking, or deployment. They do, however, reject a simplistic picture in which all ITS sequence is merely one homogeneous by-product pool. Together with the primary temporal model and causal plaque/titer perturbations, they support a mechanistic picture in which infection reorganizes flux among distinguishable ITS states (**Figure 9F**).

### 4. Causal manipulation of ITS2 converts the deployment signal into a reciprocal HSV-1 phenotype

The dynamical analyses above identify infection-responsive ITS deployment as the most parsimonious explanation of the RNA trajectories, but statistical preference among kinetic models is not itself evidence that ITSs affect viral fitness. We therefore moved from inference to intervention. We prioritized ITS2 for functional perturbation because it was the spacer showing the strongest HSV-responsive departure from the stoichiometry expected for products of a common 45S precursor. ITS2 repeatedly decoupled from both total 45S and ITS1 across time and subcellular compartments, and its sequence-resolved behavior was altered by deletion of the viral *vhs* RNA-processing factor. The experimental intervention used two deliberately opposite perturbations (**Figure 10A**). In a loss-of-function arm, we depleted endogenous ITS2-containing RNA with sequence-specific siRNAs and asked whether removing ITS2 activity released HSV-1 replication. In a reciprocal gain-of-function arm, we supplied synthetic sense-strand single-stranded ITS2 RNA oligonucleotides (ssRNAs) and asked whether increasing the cellular pool of the same ITS2 sequence elements suppressed replication. These manipulations make opposite predictions under an antiviral model: depletion should increase viral output, whereas exogenous enrichment should decrease it (**Figure 10A**).

**Figure 10.**
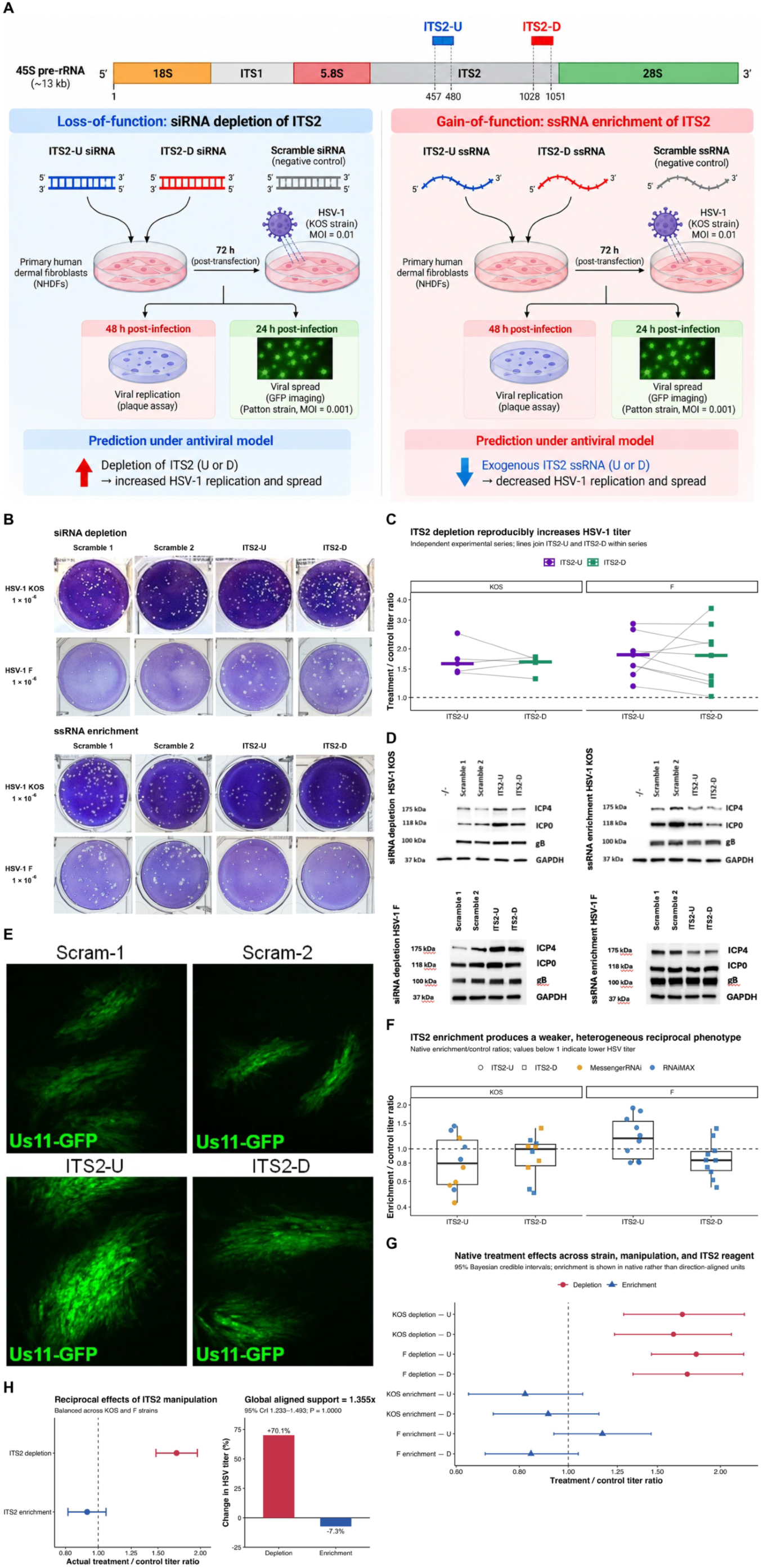
Effects of reciprocal manipulation of ITS2 on HSV-1 replication and spread. (A) Experimental design for reciprocal loss- and gain-of-function perturbations of ITS2 during HSV-1 infection. Schematic of the human 45S pre-rRNA transcription unit showing the positions of two ITS2 target regions: ITS2-U (nucleotides 457–480) and ITS2-D (nucleotides 1028–1051), numbered relative to the 5′ end of ITS2. Primary normal human dermal fibroblasts (NHDFs) were transfected with either sequence-specific duplex siRNAs targeting endogenous ITS2-U or ITS2-D (depletion) or synthetic sense-strand single-stranded RNA oligonucleotides (ssRNAs) corresponding to the same regions (enrichment). Randomized scramble oligonucleotides served as negative controls. At 72 h post-transfection, cells were infected with HSV-1, and infectious viral progeny were quantified by plaque assay. **(B) Representative plaque assays following ITS2 depletion (upper two rows) or enrichment (lower two rows) in NHDFs infected with HSV-1 KOS or F strains.** Columns show the two scrambled controls (Scramble 1 and Scramble 2) and the ITS2-U and ITS2-D treatments. Images illustrate the reciprocal effects of ITS2 manipulation on infectious viral output. Plaques are shown at a viral dilution of 10⁻⁶. **(C) Reproducibility of the ITS2 depletion phenotype across independent experimental series.** Bayesian estimates of treatment-to-control viral titer ratios are shown separately for ITS2-U and ITS2-D in HSV-1 KOS (five independent experiments) and F (nine independent experiments). Both siRNAs increased infectious viral output in every independent series (14/14 for each target). Pooled posterior titer ratios were 1.713 (95% credible interval [CrI], 1.490–2.016) for ITS2-U and 1.667 (95% CrI, 1.384–2.015) for ITS2-D, indicating approximately 67%–71% higher viral titers following depletion of endogenous ITS2-containing RNA. **(D) Western blot analysis of HSV-1 immediate-early proteins ICP0 and ICP4 following ITS2 manipulation.** Depletion of either ITS2-U or ITS2-D increased viral protein accumulation relative to scrambled controls, corroborating the increased infectious viral output observed in plaque assays. GAPDH served as a loading control. **(E) Effects of ITS2 manipulation on HSV-1 cell-to-cell spread.** NHDFs were transfected with ITS2-targeting siRNAs or synthetic ITS2-derived ssRNAs and infected with recombinant HSV-1 Patton expressing US11-GFP. Viral spread was monitored by fluorescence microscopy, and plaque areas were quantified in µm². ITS2 depletion increased plaque size relative to scrambled controls, consistent with enhanced viral spread (Šidák multiple-comparison tests, *P* < 0.05 for significant comparisons). **(F) Heterogeneous effects of exogenous ITS2-derived ssRNA enrichment on HSV-1 replication.** Native enrichment-to-control viral titer ratios were estimated independently for ITS2-U and ITS2-D across KOS and F experimental series. In KOS, posterior ratios were 0.804 (95% CrI, 0.559–1.160) for ITS2-U and 0.889 (95% CrI, 0.653–1.218) for ITS2-D. Enrichment with ITS2-D also tended to suppress F-strain replication (posterior ratio approximately 0.844), whereas ITS2-U enrichment in F produced an opposite-direction response (approximately 1.168). Thus, synthetic ITS2 enrichment generally favored reduced viral replication but exhibited substantially greater heterogeneity than endogenous ITS2 depletion. **(G) Unified comparison of the reciprocal ITS2 perturbations, retaining native treatment-to-control viral titer ratios.** Depletion ratios greater than 1 indicate increased viral replication following removal of endogenous ITS2-containing RNA, whereas enrichment ratios below 1 indicate reduced replication following addition of synthetic ITS2-derived RNA. Posterior estimates and credible intervals summarize effects across HSV-1 strains and ITS2 target regions, illustrating the strong, consistent depletion phenotype alongside the weaker and more heterogeneous enrichment response. **(H)** Integrated biological summary of the reciprocal perturbation experiments. In the unified Bayesian analysis, depletion of endogenous ITS2 increased HSV-1 titer by a factor of 1.701 (95% CrI, 1.478–1.960), corresponding to approximately 70% greater infectious viral output. Conversely, synthetic ITS2 enrichment produced a smaller overall reduction, with a native enrichment/control ratio of 0.927 (equivalent to a direction-aligned factor of 1.079; 95% CrI, 0.948–1.229). The balanced direction-aligned support factor across viral strains, target regions, and manipulation directions was 1.355 (95% CrI, 1.233–1.493), with posterior probability of the hypothesized antiviral direction >0.9999. Together, these experiments support an antiviral role for endogenous ITS2-containing RNA while revealing that supplementation with short synthetic ITS2 fragments does not fully reproduce its endogenous activity.

Within ITS2, we perturbed two spatially separated motifs: ITS2-U located in the upstream half of the spacer (positions 457–480) and ITS2-D mapping near the 3′ end (positions 1028–1051). For depletion, primary NHDFs were transfected with custom duplex siRNAs directed against U or D, with randomized scramble siRNAs as negative controls. For enrichment, parallel NHDF cultures received chemically synthesized sense-strand ssRNA oligonucleotides corresponding to the U or D sequence, again compared with scrambled RNA controls. While dermal fibroblasts were initially considered immune-neutral cells, it is now recognized that they actively contribute to skin defense against pathogens through direct recognition, production of immunological molecules, and immune regulation (Davidson et al., 2021; Wang et al., 2023). Cells were incubated for 72 h after either manipulation before low-MOI HSV-1 infection, viral spread was allowed for 48 h, and infectious progeny were quantified by plaque assay. Thus, U and D were tested twice in opposite experimental directions: sequence-specific removal of endogenous target-containing RNA versus addition of the corresponding RNA sequence. Before infecting the cells, we verified that siRNA treatment produced a local ITS2 perturbation without globally suppressing ribosomal RNA production. RNA sequencing of uninfected NHDFs without rRNA depletion was performed 72 and 120 h after transfection, and reads containing the cognate U or D motif were normalized to RNA derived from the remainder of the same rRNA cistron. At 72 h, the interval used before infection, the cognate siRNAs reduced target-containing ITS2 RNA by approximately 40–83% relative to randomized scramble controls (**Figure 10B**); by 120 h the effect had attenuated to approximately 22–29%. Total rRNA abundance changed by <2% and was not significant (*P*=0.662). Thus, the depletion arm produced a measurable, local loss of ITS2 sequence on the relevant experimental timescale rather than a gross shutdown of ribosome biogenesis. The enrichment arm was conceptually different: it did not attempt to reverse this knockdown quantitatively, but supplied a large exogenous pool of the corresponding synthetic U or D ssRNA to test the reciprocal gain-of-function prediction. Only after the two experimental arms had been analyzed in their native units did we integrate them in a single reciprocal-effect model. To place the opposite predictions on one common scale, enrichment, not the experimental data themselves, was sign-reversed on the log-ratio scale, so that positive Y always denotes movement in the predicted antiviral direction:

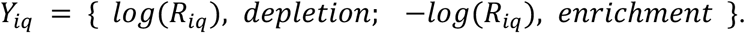

The unified hierarchical model included HSV-1 strain (KOS or F) used, manipulation (depletion or enrichment), oligo (U or D) and their interactions, with experimental block as a random effect and a Student-t residual distribution (ν=4) to limit the leverage of highly variable series.

Conceptually,

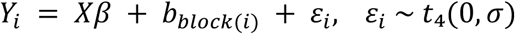

The global estimand was deliberately balanced across strain × manipulation × oligo cells rather than allowing the more numerous observations in any one subgroup to dominate. On this direction-aligned scale, exponentiating the balanced posterior mean gives a support factor A; A>1 indicates that the data, taken together, follow the predicted pattern of higher HSV yield after ITS2 depletion and lower yield after ITS2 enrichment. Native enrichment effects are reported as ordinary enrichment/control ratios, not as direction-aligned factors.

The depletion result was strikingly reproducible (**Figure 10C; SI, Table S37**). In HSV-1 KOS, both ITS2-U and ITS2-D increased viral titer in all five independent experiments. The Bayesian treatment/control ratio **(SI, Table S38)** was 1.630 for ITS2-U (95% credible interval [CrI], 1.274–2.184; P[R>1]=0.9989) and 1.655 for ITS2-D (95% CrI, 1.382–1.883; P[R>1]=0.99993).

The effect was not peculiar to KOS. In nine independent experiments with HSV-1 F, both U and D again increased titer in 9/9 series, yielding posterior ratios of 1.760 (95% CrI, 1.416–2.196; P[R>1]=1.000) and 1.724 (95% CrI, 1.243–2.414; P[R>1]=0.9979), respectively. Across the 14 independent KOS+F experiments, the pooled effects were 1.713 for U (95% CrI, 1.490–2.016) and 1.667 for D (95% CrI, 1.384–2.015), with posterior directional probability 1.000 for both. In biological terms, lowering either of two distant ITS2 motifs increased infectious HSV output by roughly two-thirds to three-quarters, and it did so across two viral genetic backgrounds. The RNAseq time courses indicated that HSV infection changes the flux and fate of ITS2-containing RNA, consistent with the deployment model favored rather than a passive precursor model; the intervention now shows that reducing this ITS2 pool before infection benefits the virus. The direction is therefore the one predicted if the infection-responsive ITS2 state is functionally restrictive to HSV rather than a neutral by-product of altered ribosome biogenesis. The agreement of U and D is especially informative because their sequences, positions and local contexts differ substantially, yet depletion of either produces essentially the same phenotype.

We corroborated our findings by repeating these experiments and harvesting cell lysate to probe for changes in the levels of viral proteins by Western blot analysis. We observed increased levels of both HSV-1 proteins ICPO and ICP4 during infection when ITS2-U and -D RNAs were depleted by siRNAs (**Figure 10D**). Since we postulated that immune function is impaired after knockdown, we also suspected that HSV-1 would spread more efficiently during infection. To test this, we infected NHDFs with HSV-1 encoding recombinant Us11 protein fused to a GFP reporter (HSV-1-Us11-GFP) and monitored changes in plaque sizes by live-cell fluorescence microscopy (**Figure 10E**). We quantified the average area (µm^2^) of plaques after ∼36 hours post infection and observed that the average plaque sizes overall increased in HSV-1 infected cells (Šidák tests *P*<0.05), a pattern consistent with the viral titer results.

The reciprocal gain-of-function arm—exogenous addition of synthetic U or D ssRNA—produced a weaker and more heterogeneous effect on HSV-1 replication than siRNA depletion (**Figure 10F; SI, Table S39**). In KOS, enrichment produced native enrichment/control posterior ratios of 0.804 for ITS2-U (95% CrI, 0.559–1.160; P[titer reduction]=0.904) and 0.889 for ITS2-D (95% CrI, 0.653–1.218; P=0.803), both in the predicted direction. In F-strain experiments, ITS2-D likewise tended to reduce titer (native ratio ≈0.844; P[titer reduction]≈0.944), whereas ITS2-U was the principal opposite-direction subgroup (≈1.168). These data therefore distinguish a robust loss-of-function phenotype from a weaker gain-of-function phenotype: removing endogenous ITS2 consistently releases HSV replication, whereas supplying short synthetic ITS2 RNA tends overall to restrain replication but does not recreate the endogenous state with comparable strength. The depletion and enrichment arms were designed to test reciprocal predictions but were not expected to produce effects of equal magnitude. siRNA recognition of U or D can eliminate longer endogenous ITS2-containing RNA species and thereby deplete molecules that have undergone their native processing, folding, localization, and ribonucleoprotein assembly. In contrast, enrichment introduces short synthetic sense-strand U or D RNAs; increasing their cellular abundance does not ensure their incorporation into the same functional RNA pool. Yet, the suppression of viral output was accompanied by reduced accumulation of key HSV-1 proteins, including ICP0 and ICP4, as measured by Western blot analysis (**Figure 10D**), supporting a model in which ITS2-derived RNAs interfere with viral gene expression or replication. Similarly, quantification of average plaque areas revealed a pronounced decrease in viral spread in cells pretreated with ITS2 U and D ssRNAs (ANOVA *F*=25.00, *P*<0.0001; Fig. 4E). These results are directionally opposite to those observed in siRNA-mediated depletion experiments and collectively support a model in which ITS2-derived transcripts modulate cellular resistance to viral infection.

Having established the depletion and enrichment arms separately, we compared their effects across both HSV-1 strains and both ITS2 oligonucleotides, retaining the original treatment-to-control titer ratios to preserve their biological interpretation (**Figure 10G; SI, Table S40**). The unified model then asked whether these opposite native effects collectively supported an antiviral role for ITS2 while accommodating their unequal magnitudes. After equal weighting of viral strain, manipulation direction and oligo, the global direction-aligned support factor was 1.355 (95% CrI, 1.233–1.493), with posterior probability of the hypothesized direction equal to 1.000 and P(effect>10%)=0.9999. The depletion component alone was 1.701 (95% CrI, 1.478–1.960), corresponding directly to an approximately 70% increase in HSV titer when ITS2 was lowered (**Figure 10H; SI, Table S41**). The enrichment component was only 1.079 on the direction-aligned scale (95% CrI, 0.948–1.229; P=0.881), corresponding in native units to an enrichment/control ratio of 0.927, or an estimated 7.3% reduction in titer. Thus, the integrated analysis supported reciprocal effects on viral replication, but with markedly asymmetric magnitudes.

The conversion between the aligned enrichment factor and the native biological effect is simply:

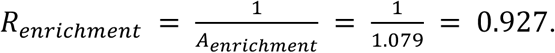

This integrated conclusion was not created by a particular outlier decision. The only F-enrichment block objectively flagged by the robust influence diagnostic (260306B) was omitted from the primary analysis; retaining it gave a virtually identical global support factor (1.354 versus 1.355), and additionally excluding the ambiguously duplicated 260529B2 label yielded 1.353. Conventional experiment-as-unit analyses independently reached the same central result: across all 14 depletion experiments, geometric mean titer ratios were 1.767 for ITS2-U (95% CI, 1.523–2.051; P=1.56×10⁻⁶; **SI, Table S42**) and 1.719 for ITS2-D (95% CI, 1.411–2.095; P=5.10×10⁻⁵). Thus, the strong depletion phenotype does not depend on Bayesian modeling, distributional fine-tuning, or one experimental series.

The perturbation experiments therefore provide the causal counterpart to the statistical deployment result. The sequence-resolved RNA analyses show that HSV-1 reorganizes ITS2 in time, space, and sequence composition, while direct manipulation shows that removing ITS2 reproducibly increases infectious virus and increasing the corresponding RNA tends overall to shift viral yield in the opposite direction. Importantly, the functional U and D motifs do not overlap the HSV-homologous blocks identified in the refined strain-17 map. The antiviral phenotype therefore cannot be explained simply by direct complementarity of these particular oligonucleotides to HSV-like ITS2 sequence. Instead, the data support a broader model in which ITS2-derived RNA contributes to the host antiviral state, with sequence context, processing, and deployment determining how strongly a synthetic gain-of-function manipulation reproduces endogenous activity. This functional result substantially strengthens the dynamical evidence for M2 by adding an independent causal dimension to the model. RNA dynamics alone already favored infection-responsive ITS deployment, with approximately 41:1 evidence for M2 over M1, whereas the functional experiments directly test the biological consequence of that inferred deployment. Before integrating these two complementary sources of evidence, we further asked whether perturbation of the viral RNA-remodeling machinery reveals the regional structure expected from a dynamically reorganized ITS2 system.

### 5. Causal perturbation supplies the information that RNA trajectories cannot

We therefore treated the experiments as a second pass through the same four biological hypotheses. The functional dataset contained 68 KOS and F-strain plaque/titer observations distributed across 29 experimental blocks. Outcomes were direction-aligned so that positive values always represented the predicted antiviral direction: for depletion, *y* = log(treatment/ control); for enrichment, *y* = −log(treatment/control). Thus depletion-induced increases in virus and enrichment-induced decreases in virus both support positive host antiviral activity. The previously fitted unified Bayesian analysis estimated approximately 1.70-fold higher HSV titer after ITS2 depletion (95% credible interval 1.48-1.96) and a weaker, heterogeneous enrichment effect. A 5,000-resample block bootstrap nevertheless placed the balanced direction-aligned effect above zero in all resamples. For formal model evidence, M0 and M1 fixed the balanced host antiviral effect *θ* to zero, whereas M2 and M3 estimated *θ* ≥ 0 under a proper half-normal prior. The likelihood retained Student-t residuals with four degrees of freedom to reduce sensitivity to outlying plaque/titer measurements and included experimental-block heterogeneity. Because explicitly sampling 29 block intercepts created a high-dimensional marginal-likelihood problem, each Gaussian block effect was integrated numerically by Gauss-Hermite quadrature.

Twenty-five quadrature nodes were sufficient: at a representative posterior point, the collapsed log likelihood differed from a 41-node calculation by only 3.6 x 10^-10^ log units.

Two independent 30,000-draw importance-sampling replicates then estimated the collapsed functional evidence. Null log evidences **(SI, Table S43)** were -44.8440 and -44.8409 (ESS 13,241 and 14,038), whereas deployment log evidences were -36.2710 and -36.2949 (ESS 2,068 and 7,758). Replicate disagreement was only 0.003 log units for the null and 0.024 for deployment, far inside the prespecified 0.15-log-unit stopping criterion. Averaging replicate log evidences gave

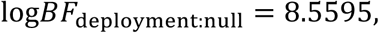

or

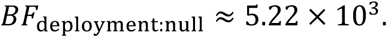

The functional experiments therefore provide approximately 5,216:1 marginal evidence for an antiviral ITS effect under the primary functional prior (**Figure 11A**). This calculation is numerically much better behaved than the earlier high-dimensional Laplace and direct-prior Monte Carlo diagnostics and is the primary functional Bayes factor (**SI, Table S43)**.

**Figure 11.**
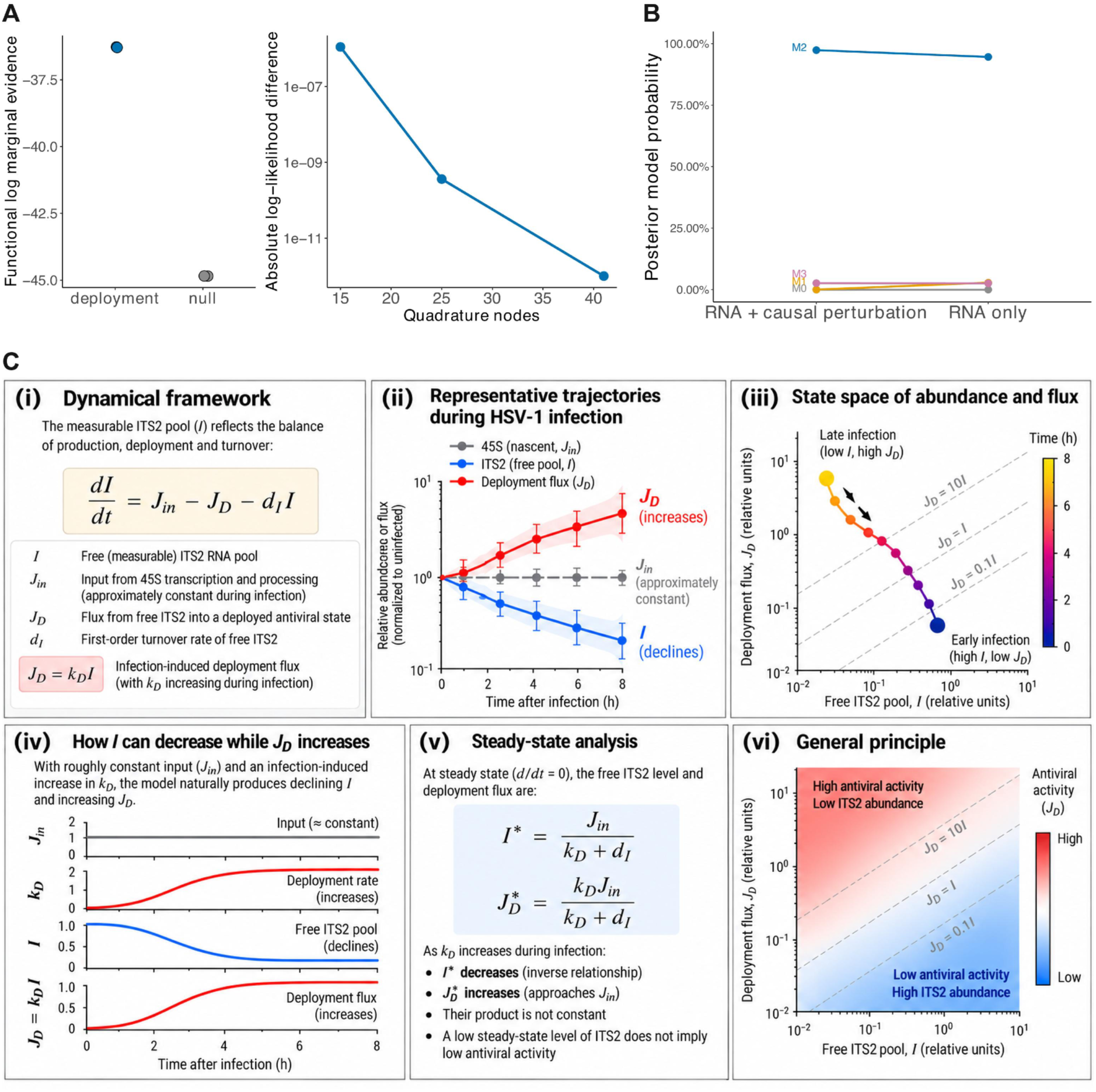
Bayesian model comparison and flux-based interpretation of ITS2 antiviral activity. **(A)** Independent ITS2 perturbation experiments were incorporated into the dynamical framework to test whether the causal functional data distinguish an antiviral deployment mechanism from a model in which ITS2 has no effect on viral replication. **(Left)** Marginal likelihoods from two independent production runs of the collapsed-block analysis are shown for the deployment and null hypotheses. The closely overlapping replicate estimates demonstrate numerical reproducibility, while their large separation strongly favors ITS deployment, with a Bayes factor of approximately 5,216:1. **(Right)** Numerical convergence of the collapsed-block likelihood calculation. Each point shows the absolute difference in log likelihood relative to the high-resolution 41-node calculation as the number of quadrature nodes is increased. The error decreases by several orders of magnitude and is effectively negligible by 25 nodes, validating the 25-node Gauss–Hermite quadrature used for the production analysis. Together, the causal perturbation evidence and its numerical validation show that the functional data provide information unavailable from RNA abundance trajectories alone and strongly favor a model containing antiviral ITS deployment over the corresponding functional null. **(B) Bayesian model comparison of the rRNA dynamics with causal perturbation data included.** Posterior probabilities for the four biological models are shown after the RNA-only analysis and after combining the RNA evidence with the independent causal ITS2 perturbation data. M0 represents passive rRNA/ITS dynamics, M1 virus-induced remodeling without functional ITS deployment, M2 host ITS deployment with antiviral activity, and M3 the combined remodeling-plus-deployment model. With RNA dynamics alone, M2 receives the greatest support, but appreciable probability remains assigned to M1 and M3, reflecting the ability of alternative infection-responsive mechanisms to reproduce aspects of the observed ITS trajectories. Incorporating the depletion and enrichment experiments produces a striking redistribution of model support: M1 falls to essentially zero because it lacks a functional ITS antiviral component, whereas posterior probability becomes concentrated on the deployment-containing models, with M2 accounting for ∼97.4% and M3 for ∼2.6% of the joint evidence. M0 remains effectively unsupported. Importantly, the causal data primarily distinguish models with versus without antiviral ITS deployment; they do not strongly alter the relative M2:M3 evidence because both models contain the same functional deployment mechanism. The resulting preference for M2 therefore reflects two complementary conclusions: the perturbation experiments strongly support antiviral ITS activity, while the additional virus-driven remodeling mechanisms included in M3 are not required to explain the combined evidence. **(C) Abundance is not activity: a flux-centered interpretation of ITS2 antiviral deployment.** The dynamical model resolves the apparently paradoxical observation that measurable ITS2 can decline during infection while ITS2-dependent antiviral activity increases. **(i)** The free, measurable ITS2 pool, I, is governed by the balance between input from precursor-rRNA processing, deployment flux, and ordinary first-order turnover:

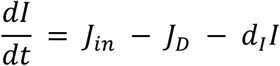 Deployment is represented by 
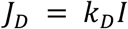
where the deployment rate k_D can increase in response to infection. **(ii)** Representative trajectories illustrate the resulting behavior. When precursor input remains approximately preserved but infection increases k_D, the measurable ITS2 pool declines even as flux into the deployed state increases. Thus, a falling RNA abundance is not equivalent to loss of biological activity. (**iii**) The same behavior is shown in abundance–flux state space. As infection progresses, the system can move from a state with relatively abundant free ITS2 but low deployment flux toward a state with substantially less measurable ITS2 but greater deployment flux. Dashed diagonals indicate combinations of I and J_D corresponding to different effective deployment rates. **(iv)** The mechanism is illustrated directly: approximately constant ITS input together with an infection-induced increase in k_D produces reciprocal changes in the free ITS2 pool and deployment flux. (**v**) At steady state, the relationships are:

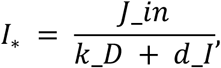

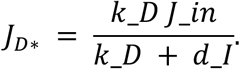
Increasing k_D therefore lowers the steady-state abundance of free ITS2 while increasing deployment flux toward the available input J_in. **(vi)** This relationship defines the general conceptual distinction between a molecular state variable and a biological flux: high abundance can coexist with low activity, whereas low abundance can coexist with high activity when material is rapidly transferred through a functional state. Applied to HSV infection, the model therefore predicts that depletion of the measurable ITS2 pool can be a signature of increased antiviral deployment rather than evidence against ITS2 function. This distinction between molecular abundance and functional flux is general and may apply whenever transient or rapidly turning-over RNA states mediate biological activity.

#### The second-pass model comparison nearly eliminates the remodeling-only explanation

Because the functional experiment distinguishes models with host antiviral deployment (M2 and M3) from models without it (M0 and M1), while the RNA dynamics distinguish the internal structures of those pairs, the independent evidence components can be combined additively on the log scale:

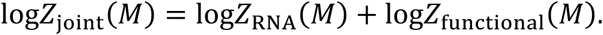

Using the frozen primary RNA importance-sampling evidence and the replicated collapsed functional evidence, joint log evidences were -55.125 for M0, -38.684 for M1, -26.615 for M2, and -30.226 for M3. With equal prior probabilities for the four models, the corresponding posterior probabilities were approximately 4.0 × 10^Z%)^for M0, 5.6 × 10^Z[^for M1, 0.97367 for M2, and 0.02632 for M3 (**Figure 11B**; **Table X; SI, Table S44**). This before-and-after comparison shows how strongly the functional experiments reinforce the dynamical evidence for host ITS deployment. RNA dynamics alone already favored M2 over M1, with a Bayes factor of approximately 33:1 by direct importance sampling (approximately 41:1 by the Laplace analysis). After incorporating the independent functional evidence, the odds increased to approximately 174,000:1 in favor of M2 over M1. By contrast, the M2:M3 odds remained approximately 37:1, because both models contain host deployment and therefore gain the same support from the functional experiments. The perturbations thus add evidence specifically for the biological feature that distinguishes M2 from the remodeling-only M1 model: ITS2 has an antiviral consequence. M2 remains preferred to M3 for a different reason: the additional viral-remodeling mechanisms in M3 provide little explanatory gain relative to their added complexity. This is stronger than simply fitting the most biologically elaborate model. HSV almost certainly remodels host transcription, RNA stability, trafficking, and ribosome biology. If that broad biological plausibility were sufficient for model selection, M3 would be declared the winner before examining the data. The formal comparison instead asks whether those additional mechanisms are necessary to explain the measured ITS phenomenon. They are not. The data favor the smallest model that contains the essential causal feature: infection-responsive host ITS deployment with antiviral consequence.

#### A flux-centered interpretation reconciles declining ITS abundance with antiviral function

The final model should not be interpreted as claiming that “more ITS2 RNA is always antiviral.” M2 predicts something subtler. The biologically relevant variable can be deployment flux or cumulative deployed activity rather than standing abundance. Define the fraction of newly generated ITS material entering deployment as

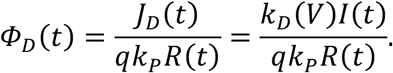

A rise in *Φ*_3_ can coexist with a fall in *I*. Indeed, if deployment is rapid and the deployed molecules are short-lived, the strongest functional response may occur when the measurable precursor/intermediate pool is being depleted most rapidly. This framework naturally explains why total nascent 45S production can remain comparatively preserved while ITS2 falls, why ITS2 can diverge from ITS1, why regional ITS behavior can differ, and why experimentally reducing ITS2 increases viral replication. The result is therefore not simply a statistical preference among four curves. It is a change in how the phenotype is conceptualized: ITS2 abundance is a state variable, whereas antiviral action is associated with movement through the state system. **Figure 11C** summarizes this flux-centered interpretation and the logic by which the RNA-only and perturbation stages converge on M2.

#### TS2 depletion reshapes host immune gene expression in the absence of viral infection

Finally, to determine whether ITS2 depletion alters the host transcriptional state independently of HSV infection, we examined a separate RNA-seq experiment in uninfected NHDFs treated with the same ITS2-directed siRNAs used in the functional experiments above. Cells were transfected with siRNAs targeting either the upstream (ITS2-U) or downstream (ITS2-D) region of ITS2, together with the corresponding scramble negative controls, but were not exposed to HSV-1.

Thus, unlike the preceding experiments, which measured the consequences of ITS2 manipulation for viral replication, this experiment asked whether reducing ITS2 alone is sufficient to alter host gene expression in the absence of a viral stimulus. A total of 37 and 62 genes (**SI, Table S45**) were, respectively, up- and downregulated in the siRNA knockdown treatments relative to the scramble negative controls at the FDR level below 0.05. Upregulated genes, such as *PDAP1*, *PSMD2*, *Heat Shock Protein Member 8* (*HSPA8*), and *HSP90AA1* contribute to a pathway related to ficolin-1-rich granule lumen (GO term adjusted *P* = 0.013; **SI, Figure S1**).

Extracellular matrix (ECM) structural constituents were the most significantly downregulated molecular function (GO term adjusted *P*=4×10^−4^; **SI, Figure S2**). Extracellular matrix (ECM) components create a robust barrier, but different viruses, including herpesviruses, have evolved specific strategies to surmount and even exploit it for cell recognition, attachment, and entry (Kamel et al., 2020; Stavolone and Lionetti, 2017). During a viral infection, the host’s response likely involves the downregulation of various cellular processes, including the production of ECM components (Pfisterer et al., 2021). *CXCR4*, a cellular chemokine receptor gene that plays central roles in development, hematopoiesis, and immune surveillance through signaling induced by its ligand CXCL12, was among the most downregulated genes in the knockdown treatments. Its expression decreased 16-fold in relation to the scramble negative controls (FDR-adjusted *P* = 0.009). This gene, “a virus’s best friend” (Arnolds and Spencer, 2014), is a common target of a diverse group of viral pathogens as a vehicle for subversion of the host immune system. Its downregulation might represent a preemptive immune response aimed at limiting viral entry and infection. However, the impact of ITS2 transcript depletion seems to be predominantly negative, resulting in an abnormal downregulation of other genes crucial for immunity, such as *Interleukin 1 Receptor Type 1* (*IL1R1*), a cytokine receptor for *interleukin-1*, which induces an interferon (IFN)-like state suppressing viral replication in fibroblasts and likely other barrier cell types (Orzalli et al., 2018). *IFITM1* and *TNFRSF1A* that protect against infection with a variety of viruses, such as HSV-1, which enter cells via the plasma membrane (Smith et al., 2019), were also significantly under-expressed in siRNA treatments compared to scramble negative controls (FDR-adjusted *P*-values 0.0032 and 0.042, respectively). Another example is provided by the *HLA-A* gene (FDR-adjusted *P*=0.047), typically upregulated during viral infections to facilitate antigen presentation, which is crucial for the activation of immune responses against infected cells. Similarly, *G6PD* expression is typically increased in different viral infections but was significantly decreased after ITS2 siRNA treatment (FDR-adjusted *P* = 0.010).

## Discussion

Evidence suggests the host’s rRNA synthesis is frequently stimulated upon viral infection (May et al., 1976; Pockl and Wintersberger, 1980; Soprano et al., 1983), but it was generally assumed to be a consequence of subversion of the ribosomes by viruses rather than a component of immune responses (DiGiuseppe et al., 2020; DiGiuseppe et al., 2018; Meade et al., 2019).

Indirect support for a role of rRNA in innate immunity was provided by clinical presentations of patients with Shwachman-Bodian-Diamond Syndrome (SBDS), an inherited ribosomopathy known for its manifestations in pancreatic, bone marrow, and immune functions (Giri et al., 2015), associated with increased susceptibility to infectious agents, including Parvovirus B19 (Grinspan and Pikora, 2005; Miniero et al., 1996), a small DNA virus causing infectious erythema. A recent discovery that HCMV infection is, unexpectedly, amplified rather than constrained in human cells with impaired ribosome biogenesis via siRNA interference of *TIF-IA* (Bianco and Mohr, 2019) reaffirms that there is a fascinating link between viral infection and stimulation of rRNA.

Disrupting the accumulation of rRNA has been shown to induce nucleolar stress, leading to dramatic downregulation of genes and pathways (Bianco and Mohr, 2019). Curiously, among the downregulated genes many are involved in modulating the innate immune response. These include High Mobility Group Box 2 (HMGB2), a chromatin-associated protein crucial for facilitating the cytoplasmic sensing of double-stranded DNA by cGAS, as well as cytoplasmic HMGB2 and type I interferon beta (IFNB1), an essential proinflammatory cytokine (Bianco and Mohr, 2019). The transcriptional induction of type I interferons is mediated by the RNA helicase RIG-I, which plays a key role in sensing pathogen-derived RNA. Additionally, it has been demonstrated elsewhere that RIG-I can also be directly triggered by an endogenous fragment of ITS2 (Jung et al., 2020), which was also enriched in influenza A infected cells (Steinberg et al., 2021). Almost one third (31%) of all significantly differentially expressed genes due to depletion of ITS2 transcript fragments via siRNA treatment in our study overlapped with those from the experiment in which all ribosome biogenesis was suppressed through ribosome TIF-IA interference (Bianco and Mohr, 2019), including *IFITM1*, *CXCL12*, *SOD2*, and *MXRA*. *SOD2* plays a critical role in regulating RIG-I-like receptor (RLR)-mediated antiviral innate immune signaling (Wang et al., 2017), while MXRA5 (Matrix remodeling-associated protein 5, adlican) is a TGF-b1-regulated protein with anti-inflammatory and anti-fibrotic properties (Poveda et al., 2017).

Our comparative analyses reveal extensive sequence similarities between eukaryotic ITS regions and DNA viruses, with particularly pronounced associations between vertebrate ITSs and orthoherpesviruses. The scale of this correspondence is notable: after filtering alignments dominated by simple nucleotide repeats, more than 336,000 ITS–DNA virus alignments remained. In sharp contrast, comparable analyses of RNA viruses did not reveal a corresponding pattern of statistically significant sequence similarities surviving the same filtering criteria. Even among DNA viruses, the distribution was highly nonrandom, with nuclear-replicating viral groups disproportionately represented. These patterns are particularly intriguing given that ITS sequences originate in the nucleolus, placing them within the nuclear environment extensively exploited by many DNA viruses. Unlike most RNA viruses, which replicate primarily in the cytoplasm, herpesviruses and many other DNA viruses establish extensive interactions with the host nucleus and nucleolus, potentially bringing viral nucleic acids and ITS-derived transcripts into overlapping molecular environments (Hiscox, 2007; Iarovaia et al., 2021; Salvetti and Greco, 2014). Such proximity could create opportunities for sequence-dependent interactions, whether initially accidental or subsequently shaped by selection.

Moreover, the distribution of ITS–viral similarities across eukaryotic lineages and viral families broadly reflected known host–virus ecological relationships (Burand et al., 2012; McGeoch et al., 2000; Rohrmann, 2019). Particularly notable were the associations between arthropod ITS1 sequences and three families of DNA viruses with established arthropod hosts: *Baculoviridae*, *Nudiviridae*, and *Nimaviridae*, exhibiting approximately 21-, 25-, and 16-fold enrichment, respectively. Among vertebrates, the correspondence was particularly pronounced for *Orthoherpesviridae*, with ITS2–orthoherpesvirus similarities enriched approximately 332-fold relative to other host clades. In primates, the corresponding ITS2 regions also contributed disproportionately to sequence divergence between closely related species, suggesting that the genomic regions exhibiting viral resemblance are among those undergoing particularly rapid evolutionary change. Their distribution was far from uniform across eukaryotic lineages and viral families. Instead, several prominent associations broadly reflected known host–virus ecological relationships. Among vertebrates, ITS2 similarities to orthoherpesvirus sequences were particularly enriched, and the corresponding regions contributed disproportionately to sequence differences between closely related primate species.

Our discovery that ribosomal ITS sequences incorporate virus-reminiscent motifs stands apart from thus far recognized viral DNA sequences integrated into mammalian genomes, which typically originate from ancient RNA viruses, such as human endogenous retroviruses (HERVs), and tend to be longer and dispersed throughout the genome (Holloway et al., 2019; Lander et al., 2001; Mun et al., 2014). Some endogenous retrovirus genes can also undergo a process of ’domestication,’ where hosts repurpose them for new functions, including retroviral restriction and antiviral defense (Best et al., 1996; Frank et al., 2022; Ito et al., 2013).

Sequence resemblance does not establish shared ancestry, horizontal sequence transfer, or adaptive convergence. Particularly in GC-rich, repetitive sequences, similarities can arise through compositional biases, recurring motifs, or chance. Yet, even an initially accidental resemblance need not remain evolutionarily inconsequential. If a host and virus sequences neutrally converge, that correspondence could create opportunities for molecular interactions that were absent before the resemblance arose. Such interactions might involve RNA–RNA recognition, competition for nucleic-acid-binding proteins, altered RNA processing, or engagement of cellular surveillance pathways. Some could be detrimental to the host, whereas others might restrict viral replication and become targets of selection. Once an interaction influences viral replication or host fitness, selection may act on the corresponding host and viral sequences. In this scenario, accidental convergence could initiate a genomic conflict, with subsequent host–virus coevolution shaping the interacting regions. The unusual evolutionary properties of rDNA make this possibility particularly intriguing. Concerted evolution, driven by processes such as unequal crossing over and gene conversion, can spread sequence variants among repeated rDNA units. As proposed in the framework of molecular drive (Dover, 1982), such processes may substantially alter the evolutionary dynamics of repetitive sequences. If particular ITS variants confer an advantage during viral infection, molecular drive could potentially facilitate their propagation within rDNA arrays, while viral counteradaptation could favor further diversification. This would place ITS evolution at the intersection of molecular drive and Red Queen dynamics (Van Valen, 1973), providing a possible explanation for why so rapidly evolving noncoding regions are embedded within one of the most evolutionarily conserved transcriptional systems. Importantly, our structural analyses also suggest that nucleotide similarity alone cannot explain the biological behavior of these sequences. ITS regions exhibiting viral sequence resemblance frequently participate in predicted secondary structures within their native host RNA context, whereas the corresponding viral genomic segments generally display different structural properties. Thus, identical or similar nucleotide motifs need not adopt equivalent conformations or perform equivalent functions. The surrounding RNA environment may be as important as the matching sequence itself. This observation broadens the possible mechanisms beyond simple antisense recognition and raises the possibility that ITS-derived RNAs function through sequence- and structure-dependent interactions with cellular components.

The comparative observations led us to ask whether ITS sequences are biologically responsive to viral infection. Across multiple herpesvirus subfamilies, viral species and strains, and human and animal cell systems, we observed pronounced changes in ITS-derived RNA abundance and distribution. These responses were neither uniform nor readily explained by generalized changes in ribosomal RNA synthesis. This distinction is important because the nucleolus is a frequent target of viral manipulation. Viruses interact extensively with nucleolar proteins and ribosome-biogenesis pathways, and changes in rRNA transcription or processing are well-established consequences of infection (Hiscox, 2007; Iarovaia et al., 2021). Under a conventional interpretation, altered ITS abundance would simply reflect these perturbations: ITSs are produced as part of pre-rRNA, so changes in their levels would be secondary consequences of changes in precursor synthesis, processing, or degradation. However, our findings reveal a more complicated relationship. For example, during murine gammaherpesvirus 68 infection, newly synthesized ITS1 and ITS2 were profoundly depleted even as mature rRNA-associated signals increased. In the HSV-1 system, the temporal response was likewise inconsistent with a simple proportional reduction in 45S transcription. Nascent 45S production remained comparatively preserved during intervals in which ITS2 declined substantially. Furthermore, ITS2 behavior differed from that of ITS1, varied across cellular compartments, and exhibited regional differences associated with the viral RNA-processing factor *vhs*. These observations indicate that the ITS component of ribosome biogenesis cannot be treated as a passive, stoichiometric byproduct of mature rRNA production. Viral infection can reorganize the fate of spacer-derived RNA independently of the overall transcriptional output of the 45S unit.

By examining distinct herpesvirus subfamilies, multiple viral strains, different host species and cell types, and complementary transcriptomic methodologies, we were able to distinguish recurring features of ITS remodeling from responses restricted to a particular infection system. ITS-derived RNAs appear to participate in a dynamic host–virus interface whose behavior depends on viral biology, infection stage, cellular context, and the mechanisms through which individual viruses perturb host RNA metabolism. An infection-responsive RNA system need not produce the same abundance change in every infection. Depending on the relative rates of transcription, processing, stabilization, transport, and degradation, an active response could manifest as either accumulation or depletion of measurable RNA. Consequently, interpreting ITS abundance without considering the underlying RNA fluxes risks confusing the amount of material present in a compartment with the biological activity passing through it.

This ambiguity motivated a central methodological feature of our study. Changes in ITS abundance, however striking, cannot by themselves establish an antiviral function. Viral interference with ribosome biogenesis could generate many of the same observations expected from an infection-responsive host defense mechanism. Moreover, the two processes are not mutually exclusive: a virus may simultaneously remodel host RNA metabolism while the host mobilizes ITS-derived RNA in response. Rather than selecting a favored explanation based on the visual appearance of RNA trajectories, we rigorously formulated these possibilities as competing mechanistic models. The four-model framework encompassed passive ribosome biogenesis (M0), virus-directed RNA remodeling without antiviral ITS function (M1), infection-responsive host ITS deployment (M2), and a hybrid mechanism combining viral remodeling with host deployment (M3). This framework addresses an important limitation of conventional model fitting. A sufficiently flexible model may reproduce observed RNA trajectories without uniquely identifying the biological processes responsible. Indeed, our analyses demonstrated that remodeling-only and deployment models can generate superficially similar dynamics. Adding parameters can improve maximum likelihood while introducing poorly constrained combinations of processes. The most elaborate model is therefore not necessarily the most informative or best supported. Bayesian marginal likelihoods are particularly valuable in this setting because they evaluate the explanatory performance of a model across its prior parameter space rather than only at its best-fitting parameter combination (Gelman et al., 2013b; Kass and Raftery, 1995; MacKay, 2003). By integrating over parameter uncertainty, they balance fit against flexibility and quantify the relative evidence for alternative mechanisms. This approach is especially useful when models differ in complexity, when parameters are only partially identifiable, and when different experimental measurements constrain different components of the same biological system.

Our RNA-only analysis favored the host-deployment model M2, with approximately 41:1 Bayesian evidence over the remodeling-only model M1 under the primary model specification. Independent importance-sampling calculations supported this result, while the hybrid model M3 received substantially less support despite its greater flexibility. Thus, the data did not require the additional remodeling parameters of M3 to explain the distinctive ITS trajectories once infection-responsive deployment was represented. We nevertheless regarded the RNA-only comparison as evidence for a particular organization of ITS dynamics, not as definitive proof of antiviral activity. The decisive additional information came from experimental perturbation. The functional experiments provided a direct test of whether ITS2 influences viral replication. Across independent experiments involving HSV-1 KOS and F strains and two spatially separated ITS2 targets, depletion of ITS2 consistently increased infectious viral yield. The effect was reproducible across experimental blocks and was accompanied by increased accumulation of viral proteins, including ICP0 and ICP4. The magnitude and consistency of this response are particularly important. Across the depletion experiments, the unified Bayesian analysis estimated approximately a 1.70-fold increase in HSV-1 titer following ITS2 depletion, with a 95% credible interval of approximately 1.48–1.96. Conventional experiment-level analyses independently confirmed the phenotype, indicating that it was not dependent on the assumptions of the Bayesian model.

The reciprocal enrichment experiments were more heterogeneous. Supplementation with synthetic ITS2-derived RNA generally shifted viral yield in the opposite direction, but the effects were weaker and less consistent than those observed after depletion. While depletion interferes with an endogenous RNA system operating within its native transcriptional, processing, and subcellular environment, delivery of synthetic oligonucleotides may not reproduce the abundance, structure, localization, processing, or molecular associations of naturally generated RNA (Deprey et al., 2020). The antiviral activity of ITS2 may therefore depend on how the RNA is produced and deployed, rather than simply on the total quantity of an ITS2-derived sequence present in the cell. Our unified hierarchical analysis explicitly accommodated these differences. Instead of allowing the stronger depletion effects or more extensively sampled experimental conditions to dominate the conclusion, it balanced evidence across viral strains, manipulation directions, and ITS2 oligonucleotides. This produced a coherent estimate of the overall antiviral response while retaining the heterogeneity of the underlying experiments. Notably, the functional model comparison yielded approximately 5,216:1 marginal evidence in favor of an antiviral ITS effect over the corresponding nonfunctional alternative. This Bayes factor was obtained from the experimental perturbation data under the primary functional prior, with independent importance-sampling replicates providing closely concordant evidence estimates. It therefore represents a direct quantitative assessment of the antiviral phenotype rather than a re-expression of the earlier RNA-only model comparison. The distinction is consequential. The approximately 41:1 RNA-only evidence favors deployment as an explanation of infection-associated ITS dynamics. The approximately 5,216:1 functional evidence addresses a different and more fundamental question: whether ITS2 manipulation affects viral replication in the predicted antiviral direction. The latter result is particularly compelling because the competing explanations make different predictions about the consequences of intervention, rather than merely about the behavior of endogenous RNA during infection.

When the independent dynamical and functional evidence components were integrated, the host-deployment model M2 received approximately 97.4% posterior model probability among the four specified mechanisms under equal model priors. The remaining support was concentrated in the hybrid model M3, whereas explanations lacking an antiviral ITS component received negligible posterior support. Thus, the integrated analysis did not merely identify a model capable of reproducing the observations. It identified the mechanism that most economically accounts for both the infection-associated RNA dynamics and the consequences of directly manipulating ITS2. This is the principal strength of our Bayesian framework: it converts several individually informative but potentially ambiguous observations into a coherent comparison of explicit biological hypotheses. The genomic similarities motivate the hypothesis; the infection-resolved RNA data constrain its dynamics; the perturbation experiments test its functional consequences; and the integrated model evaluates whether the same biological explanation can account for both observational and interventional evidence. The Bayes factor should nevertheless be interpreted cautiously: ratios of marginal likelihoods depend on the specific competing functional models and priors, while the posterior model probabilities describe relative support among the four models considered, rather than the probability that all relevant biological mechanisms have been exhaustively represented.

The possibility that ITS-derived RNAs contribute to antiviral defense invites a reconsideration of the evolutionary organization of the 45S rRNA system. Ribosomes are indispensable for viral protein synthesis, and viruses have repeatedly evolved mechanisms to manipulate ribosome biogenesis and nucleolar function. Our findings suggest that the same transcriptional system exploited by viruses may also generate RNA products that restrict them. This arrangement would create an unusual evolutionary relationship. Mature rRNAs are subject to exceptionally strong functional constraints because of their central role in translation. ITS sequences, in contrast, are rapidly evolving components of the same transcriptional unit. Their location may permit the accumulation of sequence variation without equivalent changes in mature ribosomal RNA function, while their obligatory transcription provides a continuous source of RNA substrates that could potentially be recruited into cellular defense pathways. The evolutionary dynamics of multicopy rDNA arrays may further distinguish this system from conventional antiviral genes.

Sequence variants can spread among rDNA copies through concerted evolution, potentially amplifying the representation of functionally advantageous motifs. This evolutionary malleability may make ITS sequences particularly amenable to antiviral adaptation, providing a capacity for sequence diversification and functional responsiveness reminiscent of immune defense systems, while preserving the essential translational functions of mature rRNAs.

Several questions remain unresolved. The mechanism through which ITS2 influences viral replication remains to be established. Direct interactions with viral nucleic acids, regulation of host antiviral signaling, modulation of RNA-binding proteins, and effects on cellular RNA metabolism are not mutually exclusive possibilities. The extent to which the herpesvirus phenotypes generalize to other DNA viruses requires direct functional testing. Finally, the evolutionary relationship between ITS–virus sequence similarities and ITS antiviral activity remains an open question. Comparative patterns may identify candidate regions and evolutionary histories, but demonstrating selection driven by viral interactions will require additional sequence-level and functional evidence. Nevertheless, the implications of our findings extend well beyond these unresolved questions. They identify a previously unrecognized potential antiviral function within one of the most fundamental and evolutionarily ancient transcriptional systems in eukaryotic cells, opening an entirely new avenue for investigating host–virus interactions. The possibility that ITS-derived RNAs contribute to innate antiviral defense also raises intriguing therapeutic prospects. Understanding and potentially harnessing their antiviral or immunomodulatory properties could provide new strategies for restricting viral replication or enhancing host defenses. More broadly, our findings suggest that the repertoire of endogenous antiviral molecules may extend into genomic regions traditionally regarded as transient byproducts of essential housekeeping processes.

## Materials and Methods

### 1. Experimental cell systems, virus stocks, ITS perturbations, and viral phenotyping Cells

Certified normal human dermal fibroblasts (NHDFs) from Lonza were grown in Dulbecco’s Modified Eagle Medium (DMEM; GenClone) supplemented with 10% Fetal Bovine Serum (FBS), 2 mM L-glutamine, and 100 U/mL penicillin-streptomycin and maintained at 37°C in 5% CO2. African green monkey kidney cells (BSC40) from ATCC were cultured in DMEM supplemented with 2% FBS, 2 mM L-Glutamine, 100 U/mL penicillin-streptomycin, and maintained at 37°C in 5% CO2.

#### Viruses

The KOS, F and Patton strains of human herpes simplex 1 (HSV-1) were obtained from Dr. Derek Walsh, Northwestern University Feinberg School of Medicine. The Patton strain of HSV-1 encodes a green fluorescent protein (GFP) fused to the viral US11 protein. HSV-1 strains were propagated by infecting BSC40 cells with a low multiplicity of infection (MOI) (0.003) and allowing it to grow for 72 hours then collecting cells and supernatant. Cells and supernatant were subjected to three freeze-thaw cycles. Cellular debris was removed by centrifugation, supernatant was aliquoted and stored at – 80°C until use. The viral titers were determined by the standard plaque assay. Briefly, viral samples were diluted 10-fold in a complete 2% FBS/DMEM media to the final concentration of 10^-8^. BSC40 cells were grown to 80-90% confluency and infected with the virus dilutions for 1 hour at 37°C, with a gentle rocking, after which virus was removed.

Cells were overlayed with 0.4% agarose in DMEM and incubated for 72 h at 37°C in 5% CO2 to allow virus replication and plaque formation. The virus was inactivated with 10% trichloroacetic acid (TCA) and cells stained with 0.05% crystal violet/10% ethanol solution. Plaques were quantified via counting the plaque forming units in each well and taking the average of two technical replicates of each dilution from five biological replicates.

#### siRNA transfections

The regions of interest that held sequence homology to viral genomes were utilized to design custom siRNA sequences, which were ordered from IDT. The siRNA sequences created target either the upper (5’) ITS2 section ("U", 5’-TGGGGGAGGAGGAAGGCGGGTCC-3’, guide 5’-ACCCGCCUUCCUCCUCCCCCA-3’, passenger 5’-GGGGAGGAGGAAGGCGGGUCC-3′) or the downstream (3’) ITS2 section ("D", target: 5’-GCCGTCGGCCGCGCGCTCTCTCT-3′, guide 5’-AGAGAGCGCGCGGCCGACGGC-3′, passenger 5’-CGUCGGCCGCGCGCUCUCUCU-3′). Two randomized scramble siRNA were created and utilized as negative siRNA controls: “Scramble-1” (pseudo-target: 5’-GAGGGCACCCGGAGGGGGAGAGG-3′, guide 5’-UCUCCCCCUCCGGGUGCCCUC-3′, passenger 5’-GGGCACCCGGAGGGGGAGAGG-3′) and “Scramble-2” (pseudo-target: 5’-AGGGCGGCGCCGCCGCCGGAAGA-3′, guide 5’- UUCCGGCGGCGGCGCCGCCCU-3′, passenger 5’- GGCGGCGCCGCCGCCGGAAGA-3′). NHDF cells were seeded onto 6-well plates and grown to 80-90% confluency. Cells were transfected with 100 pmol of each of custom siRNAs using Lipofectamine RNAiMAX transfection reagent (Invitrogen) and following the manufacturer’s protocol. Transfected cells were incubated for 72 hours prior to infections.

#### Overexpression transfections

Chemically-synthesized single-stranded ITS2 RNA oligonucleotides corresponding to the "U" and "D" targets were used to induce overexpression/oversaturation of ITS2, along with “Scramble-1” and “Scramble-2” oligos as negative controls. NHDF cells were seeded onto 6-well plates and grown to 80-90% confluency. Cells were then transfected with 2 nmol of each oligo using Lipofectamine RNAiMAX transfection reagent (Invitrogen) according to the manufacturer’s instructions. After transfection, the cells were incubated for 72 hours prior to subsequent infection experiments.

#### Viral infection assays

NHDFs 72 hours post-transfection were infected with HSV-1 at a low MOI (0.01) and the virus was allowed to spread for 48 h at 37°C in 5% CO_2_. Cells and supernatant were subjected to three freeze-thaw cycles. Cell debris was pelleted by centrifugation and collected supernatant was aliquoted and stored at –80°C. Viral titers were determined by plaque assay as described above. Additionally, cell lysates were collected in 1X Laemmli buffer, boiled for 5 minutes, and stored at –20°C for immunoblotting. For viral spreading assays, cells were transfected with siRNAs as described above for 72 h then infected with HSV-1 Patton strain coding US11 protein fused to GFP at a low MOI (0.001) for 1 hour at 37°C, with a gentle rocking. After the incubation virus was removed and cells were overlayed with 0.4% agarose in DMEM. Infected cells were incubated for 24 hours at 37°C in 5% CO2 and then visualized by live-cell microscopy on an inverted Leica DMI4000B fluorescence microscope. Plaque sizes were measured by outlining plaques in the Leica LAS X software and measuring the area of each plaque in (µm^2^) (n = 3). Representative images of plaques were uniformly enhanced in LAS X.

#### Immunoblotting and antibodies

Cell lysates from the experiments described above were collected in 1X Laemmli buffer, boiled for 5 minutes, and stored at –20°C until use. The protein concentration of each sample was measured using Qubit 4 Fluorometer (Thermo Fisher) following manufacturer’s protocol. Samples were resolved using 10% SDS-PAGE gels and proteins were transferred to polyvinylidene difluoride (PVDF) membranes (Bio-Rad). The blots were blocked in 5% milk in 1X TBS/0.1% Tween 20 buffer (TBS-T). The membranes were probed with anti-HSV-1 ICP0 (Santa Cruz Biotechnology) overnight. After three washes with TBS-T, blots were probed with the conjugated secondary antibody anti-mouse HRP (Jackson ImmunoResearch Laboratories) for one hour at room temperature, and again washed three times with TBS-T. Membranes were visualized via Bio-Rad ChemiDoc MP using Clarity Western ECL Substrate (Bio-Rad). The blots were subsequently stripped with 0.2 M NaOH, blocked, and independently reprobed with anti-HSV-1 ICP4 (Abcam) and anti-glyceraldehyde-3-phosphate dehydrogenase (GAPDH) (Santa Cruz Biotechnology) antibodies, as described above.

### 2. RNA-seq, preprocessing, fragmentome analysis, and and positional rDNA quantification

#### RNA extraction, sequencing, and processing

Total RNA was extracted using TRIzol reagent (Invitrogen) following the manufacturer’s protocol. The first experiment involved 32 uninfected samples: four siRNA treatments/transfections targeting the upper (5′; ITS2-U) or downstream (3′; ITS2-D) ITS2 sections, and two randomized negative controls (Scramble-1 and Scramble-2), each assessed in four biological replicates at 72 and 120 h after treatment. cDNA libraries were prepared and sequenced by Genewiz (South Plainfield, NJ) on an Illumina HiSeq instrument, omitting rRNA depletion to retain ITS-containing RNA and generating 2 × 150-bp paired-end reads. hg38 and RefSeq references and annotations were obtained from UCSC (https://genome.ucsc.edu/). Reads were quality-filtered using FastqMcf (Aronesty, 2013), mapped to the gene reference with Bowtie2 (Langmead et al., 2009), and quantified with RSEM (Li and Dewey, 2011). Gene-level differential expression was evaluated with DESeq2 (Love et al., 2014), using Benjamini–Hochberg-adjusted *P* < 0.05 (Benjamini and Hochberg, 1995). ITS abundance was examined using FPKM, all-read normalization, and normalization to reads from the entire rRNA cistron; total rRNA abundance was also normalized to non-rRNA-cistron transcripts. Motif quantification used custom shell scripts operating on FASTQ files. Gene Ontology and pathway analyses used g:Profiler (Kolberg et al., 2023). Statistical analyses were performed in R 4.3.0 (R, 2025).

#### Herpesvirus RNA-seq datasets and positional ITS quantification

Public RNA-seq datasets from alpha-, beta-, and gammaherpesvirus infections were reanalyzed using infected and matched mock samples within each study. These included MCMV (GSE212286; NIH-3T3 cells, 1–72 hpi), HCMV TB40/E ((Oberstein and Shenk, 2017); MRC-5 and ARPE-19 cells, 24, 72, and 120 hpi), HSV-1 (GSE59717, GSE103763, GSE124118, GSE100576, and GSE129582), and MHV68 (GSE70481; mouse embryonic fibroblasts). The HSV-1 4sU dataset (GSE59717; (Rutkowski et al., 2015)) comprised eight successive 1-h labeling windows from 0 to 8 hpi. Total, cytoplasmic, nucleoplasmic, and chromatin-associated fractions were analyzed separately; wild-type and Δvhs infections were compared within matched experimental backgrounds.

MHV68 data were aligned to mouse rDNA. The MCMV time course included duplicate mock and infected libraries per time point; the HCMV study comprised 24 libraries (two cell lines × three time points × two infection conditions × two replicates). Biological replicates were distinguished from technical replicates and positional read counts.

After study-specific read quality control, Bowtie2 alignments to the 45S rDNA transcription unit, including reads with multiple possible rDNA placements, were used to estimate positional coverage. Exact canonical 31-mers unique to designated rDNA regions provided a complementary, more stringent measure. Read counts and fragment counts were analyzed separately, with spacers and mature-rRNA segments retained as distinct features. For region r and time t, Y denotes normalized RNA abundance; region-specific change relative to the whole 45S pool was defined as:

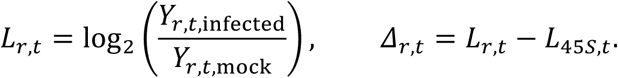

The relative ITS2-versus-ITS1 response was:

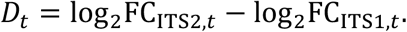

#### HSV-1 KOS small-RNA fragmentome analysis

An independent small-RNA sequencing experiment in normal human dermal fibroblasts compared four HSV-1 KOS-infected libraries with four mock-infected libraries. Paired-end 2 × 150-bp FASTQs were first inspected for library architecture. Most R1 reads contained the adapter AGATCGGAAGAGCAC downstream of a shorter biological insert (median approximately 66–67 nt). The first exact occurrence of this adapter was removed together with downstream sequence and corresponding quality scores; inserts shorter than 10 nt were excluded. Because untrimmed paired-end mapping largely reflected adapter interference, definitive fragmentome analyses used trimmed R1 inserts as single-end reads. A 500,000-read-per-library audit searched for residual adapter motifs with up to one or two mismatches to assess condition-dependent trimming bias.

Trimmed inserts were mapped end-to-end with Bowtie2 --very-sensitive to the human complete rDNA repeat U13369.1 (42,999 nt). The annotated transcribed segments (1-based coordinates) were 5′ ETS, 1–3656; 18S, 3657–5527; ITS1, 5528–6622; 5.8S, 6623–6779; ITS2, 6780–7934; 28S, 7935–12969; and 3′ ETS, 12970–13314. Mapped inserts were assigned genomic start and end coordinates and lengths, with exact fragment species defined by their start–end–length combinations. Regional overlaps, junction-spanning fragments, single-nucleotide endpoint counts, and 10-nt endpoint bins were tabulated. Generic mapped starts and ends are reference-oriented genomic coordinates, not necessarily biochemical 5′ and 3′ ends; the dominant 7951–7971 fragment was independently inspected for orientation and CIGAR structure. For feature f in replicate j, counts C were normalized both to all retained trimmed R1 reads N (input CPM) and to the reads R mapping to the transcribed 45S interval (within-45S proportion). The two denominators distinguish changes in absolute representation within the sequenced small-RNA pool from redistribution among rDNA-derived fragments:

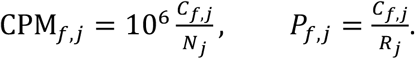

For each feature, fold change was calculated as the ratio of the arithmetic mean normalized abundance across four infected replicates to that across four mock replicates. Exact two-sided permutation tests enumerated all 70 allocations of eight libraries into two groups of four; the minimum attainable two-sided P value was 2/70 = 0.02857. Benjamini–Hochberg adjustment was applied separately within each 12-feature regional comparison family for each normalization. This fragmentome analysis therefore used replicate-level exact inference, not nucleotide positions as independent replicates:

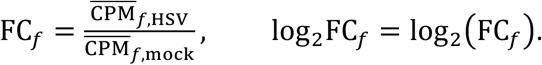

For KOS-specific homology assessment, ITS1, ITS2, and selected short-RNA species were compared with the complete HSV-1 KOS genome (JQ673480.1) using BLASTN-short (word size 7; low-complexity masking disabled). The six-scale viral-similarity framework described below was also applied to KOS, using the best local alignment for each overlapping ITS k-mer and projecting identity-weighted scores onto covered positions. For the fragmentome-specific response, the abundance at nucleotide i combined mapped start and end counts, converted to input CPM. A fixed 0.05-CPM pseudocount stabilized log ratios, and the signed and absolute-response correlations were evaluated using 10,000 within-spacer circular shifts of the KOS similarity landscape. These spatial tests were exploratory complements to the regional replicate-level tests:

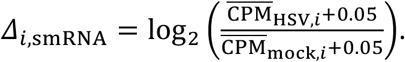

#### Statistical inference and reporting conventions

Frequentist and Bayesian approaches were used for complementary inferential purposes. Frequentist analyses, including exact tests, regression-based tests, and empirical permutation procedures, evaluated evidence against specified null hypotheses. Bayesian analyses were used to estimate effect sizes and their uncertainty, quantify posterior probabilities of directional effects, and compare competing mechanistic models through their marginal likelihoods. The choice of inferential framework depended on the scientific question and statistical structure of each analysis, as described in the corresponding methodological sections. Throughout the manuscript, italic uppercase *P* denotes frequentist *P* values, including empirical permutation-based probabilities. Where applicable, adjustments for multiple comparisons are identified explicitly. Bayesian posterior probabilities are denoted by Pr(A | D), where A represents the event or hypothesis of interest and D the observed data; these probabilities are conditional on the specified model and prior distributions. Bayesian uncertainty intervals are reported as 95% credible intervals (CrIs), whereas frequentist intervals are identified as confidence intervals (CIs). Bayes factors (BFs) quantify the ratio of marginal likelihoods under competing models and are distinguished from posterior model odds.

#### Viral-similarity landscapes and spatial inference

To assess whether viral sequence similarity predicts the spatial pattern of ITS responses, each ITS nucleotide i was assigned a sequence-only score at k = 11, 15, 21, 31, 51, and 75 nt. At each scale, the best viral alignment score for each overlapping k-mer was projected onto covered ITS positions and standardized across ITS1 and ITS2; the standardized scores were averaged to produce a fixed multiscale similarity landscape independent of the measured RNA response:

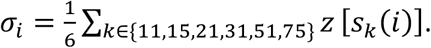

The positional RNA response Δᵢ was defined from infected-to-mock abundance at position i. Two Pearson correlations distinguished directional association (β) from association with response magnitude irrespective of sign (λ):

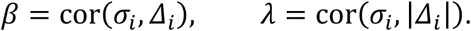

To account for spatial autocorrelation, the intact viral-similarity landscape was circularly shifted 10,000 times independently within ITS1 and ITS2, preserving spacer boundaries and internal structure while breaking alignment with the observed RNA response. For signed β, significance was two-sided; for the prespecified positive magnitude association λ, the upper tail was used. For B shifts, with Tᵦ the relevant (absolute or signed) null statistic and Tobs its observed counterpart, empirical P values included a plus-one correction:

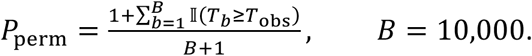

Dinucleotide-preserving pseudo-viral genomes provided an additional sequence-composition null in selected conventional RNA-seq analyses. Spatial permutations and replicate-level tests address different questions; adjacent ITS positions were never treated as independent biological replicates.

### 3. Dynamical modeling of infection-responsive ITS RNA

#### Four biological hypotheses encoded as nested dynamical models

We defined four biological models, M0-M3 (Figure **1**). These names refer only to biological hypotheses; subsequent stages such as identifiability analysis, Bayesian robustness, fractionation analysis, or functional integration are analysis stages rather than additional biological models.

#### M0, passive ribosome-biogenesis null

M0 assumes that ITS behavior is an ordinary consequence of 45S transcription, processing, and turnover. There is no host ITS antiviral state and no explicit virus-directed remodeling term. In its simplest form,

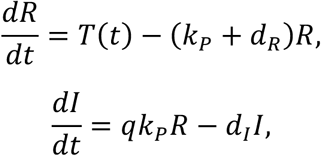

where *R* denotes the precursor-rRNA pool, *I* the measurable ITS precursor/intermediate pool, *T*(*t*) precursor transcription, *k*, processing from precursor into ITS-containing intermediates, *q* the effective yield into the measured ITS pool, and *d*_-_ and *d*_._ loss terms. Virus abundance *V* can change with time, but it does not alter ITS kinetics in M0.

#### M1, viral remodeling without host ITS function

M1 gives HSV substantial freedom to alter ribosomal transcription, processing, or ITS stability, but it still denies an antiviral ITS state. A convenient representation is

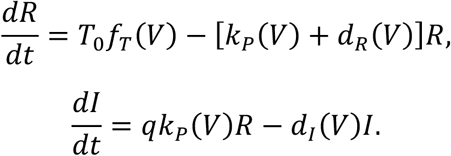

Virus dependence was represented by smooth bounded response functions, for example

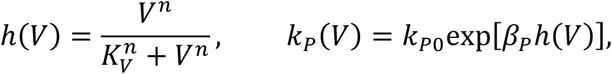

with analogous terms available for transcription or decay. Thus M1 is a deliberately generous nonfunctional alternative: it can reproduce ITS loss by allowing HSV to remodel the machinery that creates or destroys ITS-containing RNA.

#### M2, host ITS deployment

M2 introduces a distinct deployed state *D*. The measurable ITS pool can therefore decrease because ITS material leaves that pool, while the deployed state can increase or turn over rapidly:

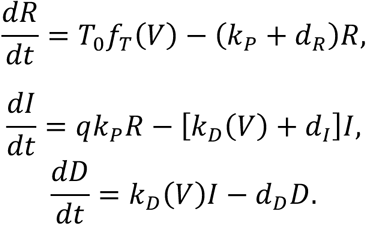

The infection-responsive deployment rate was represented as

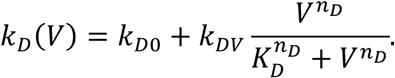

For functional experiments, the deployed state was allowed to inhibit viral expansion:

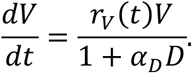

The key quantity is therefore not only standing ITS abundance but deployment flux,

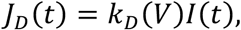

and its cumulative activity,

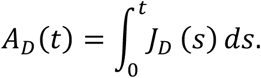

The intuition is analogous to a reservoir with an inlet and an outlet. A declining water level does not by itself show that the inlet has shut down; if the inlet remains active while the level falls, the outlet must have increased. The 4sU experiment provides information about the inlet. Maintained nascent 45S production together with declining ITS2 therefore constrains ITS2 outflux to exceed its replenishment during the critical phase of infection.

#### M3, hybrid viral remodeling plus host deployment

M3 combines the mechanisms of M1 and M2:

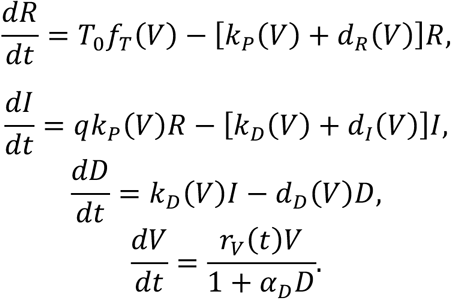

Biologically, M3 may initially appear the most realistic model because HSV unquestionably remodels host RNA metabolism. Statistically, however, the relevant question is not whether viral remodeling occurs somewhere in the infected cell. It is whether explicit remodeling parameters are needed to explain the distinctive ITS observations after host deployment has already been included. M3 was therefore retained as a strong, deliberately permissive competitor rather than excluded as an obvious biological truth.

#### Modeling strategy and separation of discovery from causal testing

The modeling workflow was designed around a two-pass principle. First, M0-M3 were compared using RNA-seq information only. This stage asked which biological structures are supported by the temporal RNA behavior without knowledge of the plaque/titer perturbations. Second, after the experimental perturbation results were introduced, the same four biological hypotheses were compared again using an independent functional likelihood. This architecture prevents circular inference and makes the contribution of causal evidence explicit. Analysis stages labeled M04, M05, M06, M07, M08, M09, or M10 in the computational workspace are workflow identifiers and must not be confused with biological models. The biological model family contains exactly four members, M0-M3.

#### HSV-1 temporal input and normalization

The primary dynamical input was derived from HSV-1-infected human foreskin fibroblast 4sU RNA-seq sampled in eight consecutive 1-h labeling intervals spanning 0-8 hpi. The primary regional matrix contained total 45S, ITS1, and ITS2 fold-change trajectories, yielding 24 observations. Fold changes were modeled on the log scale. The normalized formulation emphasizes relative infection-induced kinetics rather than attempting to infer absolute intracellular molecule numbers from sequencing counts. The observed total-45S fold changes across interval midpoints 0.5-7.5 h were 1.06, 1.04, 1.02, 1.03, 0.96, 1.05, 0.92, and 0.82. ITS1 fold changes were 1.06, 0.92, 0.76, 0.61, 0.65, 0.75, 0.84, and 0.70. ITS2 fold changes were 1.02, 0.98, 0.74, 0.54, 0.52, 0.61, 0.59, and 0.46. Mature 18S and 28S measurements were retained for descriptive and structural checks but were not used to multiply the effective sample size of the primary M0-M3 comparison.

#### State variables and fluxes

The core state variables were precursor rRNA *R*(*t*), measurable ITS intermediate *I*(*t*), deployed ITS *D*(*t*) where present, and viral burden *V*(*t*). ITS1 and ITS2 were represented by separate infection-response parameters in the fitted normalized model because the observed trajectories clearly decouple. The generic precursor-to-ITS and deployment fluxes were

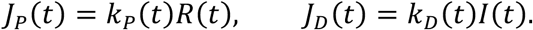

These fluxes distinguish production from standing abundance. The cumulative deployment measure was 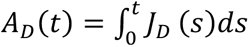.

#### Numerical solution and parameterization

ODE trajectories were integrated numerically with a fixed-step fourth-order Runge-Kutta solver implemented in base R (Butcher, 2000; Soetaert et al., 2010). Positive kinetic quantities were parameterized on transformed scales where appropriate, and bounded infection-transition parameters were represented by smooth logistic/Hill-type transformations. Optimization used repeated numerical minimization from defined starting values. The normalized v2 comparison contained 3, 6, 5, and 8 fitted parameters for M0, M1, M2, and M3, respectively.

The infection-responsive transition was parameterized by a midpoint *τ* and width *w*. A logistic representation can be written as

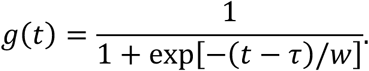

Small *w* approximates a switch; larger *w* produces a gradual transition. The fitted M1 width approached the numerical lower boundary, whereas M2 retained a finite width.

#### Frequentist objective and AICc

For the RNA-only comparison, parameters were estimated by minimizing residual error between observed and predicted log fold changes. For *n* observations and *k* fitted parameters, corrected Akaike information criterion was computed as

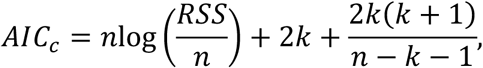

up to constants common to all models. Differences were defined relative to the best model,

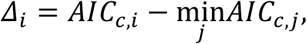

and Akaike weights (Burnham and Anderson, 2002)as

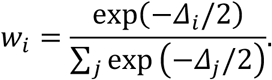

AICc was emphasized over uncorrected AIC because the number of observations was modest relative to the parameter count, especially for M3.

### Hessian and local identifiability diagnostics

Numerical Hessians of the objective or negative log posterior were evaluated at fitted optima. Eigenvalues of the symmetric Hessian were used to diagnose local curvature. Near-zero eigenvalues indicate weakly identified parameter combinations. The ratio between extreme curvature scales was used as a practical condition/sloppiness diagnostic. Hessian results were interpreted diagnostically rather than as proof of global nonidentifiability.

### Synthetic model-recovery experiment

To assess whether M1 and M2 can be distinguished under the observed design, synthetic datasets were generated from each fitted model with noise at the prespecified scale (sigma = 0.10 on the modeled scale). Twelve datasets were generated under M1 and twelve under M2. Both models were then refitted to every synthetic dataset, and the preferred model was recorded. Because the generating model was known, this analysis directly measures confusion induced by the sampling design and model geometry.

### Transition-geometry analysis

A local grid over transition midpoint and width was evaluated around the fitted M1 and M2 solutions. Nuisance parameters were held at their fitted values for this diagnostic surface. Consequently, this analysis is not described as a fully reoptimized profile likelihood. Its purpose was to visualize whether the models rely on qualitatively different transition geometries.

### Bayesian RNA-only model evidence

Bayesian evidence used proper priors and equal prior model probabilities, *P*(*M*_!_) = 0.25. Under the primary specification, infection-effect parameters had Normal(0, 1.5^(^) priors on their transformed fitted scales. The transformed transition midpoint had a Normal prior centered on the prespecified infection window, transformed log-width had a proper Normal prior, and residual noise had a half-Normal prior with scale 0.5.

Sensitivity analyses varied the effect-prior scale while preserving the model definitions and data. For each model,

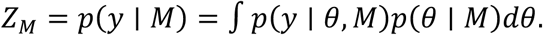

Initial evidence was obtained with a Laplace approximation around the posterior mode. This calculation is efficient but assumes that the important posterior mass is adequately represented by local quadratic curvature. Because M1 and M3 showed potentially problematic geometry, Laplace results were treated as an initial evidence calculation rather than the sole basis for inference. Because marginal likelihoods depend on the prior distribution as well as the likelihood, we explicitly evaluated sensitivity to the scale assigned to infection-effect parameters. The primary scale of SD = 1.5 was chosen as a weakly informative prior that allowed substantial infection-associated effects without assigning excessive mass to extreme parameter values; it was not estimated from the HSV data. We therefore repeated the evidence calculation using narrower (SD = 1.0) and broader (SD = 2.5) effect priors while leaving the likelihood, model structures, and remaining priors unchanged. This analysis was used to distinguish conclusions that were stable across reasonable prior scales from those dependent on the strength of regularization.

### Importance-sampling verification of RNA evidence

Posterior-centered importance proposals were used for independent numerical integration. Log weights were evaluated stably, and evidence estimates were accompanied by ESS and Monte Carlo error. Thirty thousand draws per model were used in the production analysis. M3 received additional independent tail-focused replicates because its ESS was lower than for M2. Agreement of independent M3 log-evidence estimates and their persistent separation from M2 were used to establish that the primary M2 conclusion was not a Laplace artifact. Accordingly, agreement between Laplace and subsequent direct importance-sampling estimates under a given prior specification was interpreted as evidence of numerical robustness of the marginal-likelihood calculation, rather than as evidence that the chosen prior scale itself was uniquely supported.

### Functional perturbation data and direction-aligned outcome

The functional evidence comprised 68 plaque/titer observations from HSV-1 KOS and F strains across 29 experimental blocks. Experiments included ITS2 depletion and synthetic enrichment, U- and D-targeting oligos, and relevant transfection-reagent structure. The direction-aligned response was

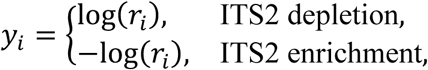

where *r_i_* is the treatment/control viral ratio. Positive *y* therefore has the same biological meaning in both manipulations: the observation points in the predicted antiviral ITS direction.

### Robust hierarchical functional likelihood

The functional model used Student-*t* residuals with *ν* = 4 degrees of freedom. Let *b*_o(!)_ denote the random intercept for experimental block *g* and *x*^0^*β* nuisance contrasts for strain, manipulation, oligo, and reagent. Then

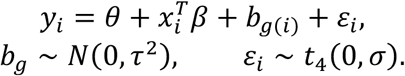

For M0 and M1, *θ* = 0. For M2 and M3, *θ* ≥ 0 and the primary prior was half-Normal with scale 0.5 on the log-ratio scale. Proper zero-centered Normal priors were used for nuisance contrasts, and proper half-Normal priors for *σ* and *τ*. Functional-prior sensitivity scales were prespecified before evidence calculation.

### Collapsing experimental-block effects by Gauss-Hermite quadrature

Directly treating all 29 *b*_o_ values as evidence-integration dimensions produced inefficient importance weights. Rather than increase Monte Carlo size indefinitely, block effects were marginalized conditionally. For block *g*,

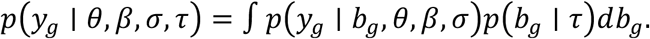

Because Student-*t* residuals are not Gaussian-conjugate with the block intercept, the one-dimensional integral was evaluated deterministically with Gauss-Hermite quadrature. The full collapsed likelihood is

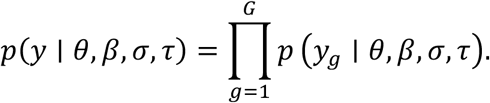

Quadrature convergence was tested at 15, 25, and 41 nodes before evidence estimation. The 25-node and 41-node log likelihoods differed by only 3.6 × 10^Z%/^ at the representative test point, far below the prespecified 0.01-log-unit criterion. Twenty-five nodes were therefore used for production evidence.

### Reduced-dimensional functional importance sampling

After block marginalization, posterior-centered defensive importance sampling was performed in the reduced parameter space. Two independent 30,000-draw runs were required for the primary null and deployment hypotheses. The prespecified stopping rule required replicate log-evidence agreement within 0.15 log units or within combined Monte Carlo uncertainty, together with materially improved ESS relative to the failed explicit-block estimator. The observed diagnostics exceeded these criteria by a wide margin.

#### Combining independent RNA and functional evidence

Because the RNA-only and functional likelihoods use independent experimental observations and the second pass preserves the same biological M0-M3 definitions, evidence was combined by multiplication, or addition on the log scale:

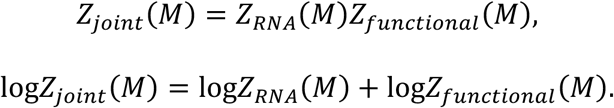

The functional evidence is identical within the no-deployment pair M0/M1 and within the deployment pair M2/M3. This is intentional: functional perturbation informs the existence of host antiviral activity, whereas the RNA dynamical component determines whether explicit viral remodeling adds explanatory value.

#### Additional fractionation and vhs-sensitive structural analyses

Fractionation and Delta-*vhs* datasets were analyzed as orthogonal structural tests. They were not allowed to inflate the primary temporal sample size. Fraction-specific predictive analyses tested whether a scalar ITS state was sufficient across compartments. Regional *vhs* analyses tested whether the H3 response was reproducible across replicates and whether a common WT-to-Delta-*vhs* reversal improved held-out prediction. These analyses constrain the state architecture but do not, by themselves, distinguish degradation from deployment.

#### Reproducibility, staged freezing, and stopping rules

Model definitions, priors, computational plans, and stopping criteria were frozen before the corresponding evidence calculations. Code and result manifests were recorded with SHA-256 checksums. Failed numerical estimators were retained as diagnostics but excluded from final evidence when they violated prespecified ESS or replicate-stability criteria. In particular, an explicit 29-block defensive importance sampler was rejected after ESS values of approximately 1-16 out of 30,000 and unstable deployment log evidences. The collapsed-block estimator replaced the numerical method without changing the biological model, data, or frozen primary priors.

### 4. Computational analysis of ITS–viral sequence similarity

#### Sequence resources and provenance

ITS1 sequences were obtained from ITSoneDB (release 131), ITS2 from ITS2 Database V (Ankenbrand et al., 2015), and viral genomes from NCBI Virus. Analyses were expanded with Vertebrate Genomes Project (VGP; (Formenti et al., 2026a)) and harmonized public spacer records. Human ITS coordinates in the sequence-response analyses refer to the frozen T2T-CHM13 rDNA reference (CP068257.2, rDNA interval 4,672,429–4,685,761), ITS1 positions 5506–6591 and ITS2 positions 6733–7912 within the rDNA unit. Mouse rDNA was indexed against BK000964.3, with ITS1 at 5878–6877 and ITS2 at 7035–8122.

#### Homology discovery and repeat-aware filtering

A sensitive nucleotide similarity screen used BLASTN 2.17.0+ (Altschul et al., 1990; Camacho et al., 2009), word size 7, DUST disabled, expectation threshold 0.05 and a minimum retained high-scoring segment length of 20 nt for the targeted cross-herpesvirus analyses. More stringent family-wide sensitivity screens used E ≤ 10⁻⁶ and, where indicated, at least 50 aligned nucleotides and sequence entropy ≥1.0 bit; the independently defined periodicity exclusion removed alignments with ≥80% coverage by motifs of period 1–6 nt. These are sensitivity criteria, not a single universal threshold. Overlapping high-scoring segment pairs (HSPs) involving the same ITS query and viral target were consolidated to avoid counting fragmented local alignments as independent observations. For each ITS sequence, the union of nucleotide positions covered by qualifying viral alignments was used to determine the proportion of the spacer exhibiting viral sequence similarity, with overlapping positions counted only once. For global alignments, Needleman-Wunsch method (Needleman and Wunsch, 1970) was used.

### Taxonomic enrichment

Host records were assigned to major eukaryotic clades and viral accessions to International Committee on Taxonomy of Viruses (ICTV) viral families. For each spacer, host clade c and family f, the primary summary counted distinct ITS queries with at least one qualifying alignment rather than the number of HSPs. The enrichment ratio compared the probability of a match within the clade with the corresponding probability in all other clades.

One-sided Fisher exact tests assessed enrichment and Benjamini–Hochberg correction was applied over the tested cells. Zero matching queries outside a clade yield an undefined/infinite risk ratio and were not silently assigned a finite estimate. Enrichment of ITS–viral sequence similarity within individual host clades and viral families was assessed using 2 × 2 contingency tables. For each host–virus category, we calculated the relative risk (RR) as the proportion of ITS sequences with qualifying viral matches within the focal category divided by the corresponding proportion outside that category. Statistical significance was evaluated using a one-sided Fisher’s exact test for enrichment. In the equations below, *a* and *b* denote the numbers of ITS sequences with and without qualifying matches in the focal category, respectively, whereas *c₀* and *d* denote the corresponding counts outside that category. The resulting *P* values were adjusted for multiple comparisons using the Benjamini–Hochberg false discovery rate procedure (Benjamini and Hochberg, 1995):

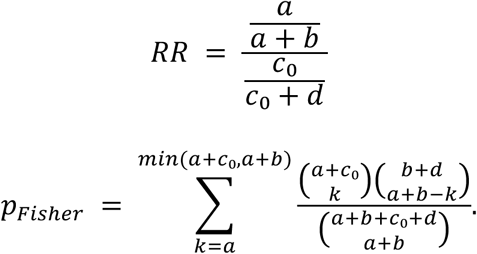

### Sequence-level controls

Candidate ITS–viral matches were compared against dinucleotide-preserving shuffled sequences, same-spacer windows matched for length, GC content and, when expression was relevant, mock-state coverage. Independent viral accessions, nonoverlapping host windows, repeat masks and composition-matched controls were used as separate robustness criteria. The *Siganus* analysis compared 20 congeners across 20 candidate viral loci; *Anolis* and *Anas* were evaluated using local sequence complexity and motif recurrence, and seven primate genera were compared at a shared orthoherpesvirus genomic hotspot near 137 kb.

### Viral replication compartments

Viral families were annotated by predominant nuclear or cytoplasmic DNA replication, with mixed and unassigned families kept separate. Analyses were repeated under stricter description-based quality control and family-balanced summaries to reduce the influence of families represented by many genomes. Because the curated vertebrate-host proxy retained only one cytoplasmic DNA-virus family, family-level nuclear-versus-cytoplasmic contrasts were treated as descriptive and not independent-family replication of a general effect.

### RNA secondary-structure analysis of host ITS and matching viral segments

Host ITS1/ITS2 RNA sequences were folded both in full-length spacer context and as isolated homologous windows; matched host windows preserved interval length and approximately matched base composition. Viral DNA segments were transcribed in silico to the corresponding RNA alphabet before folding and compared with composition-matched windows from the same viral genome. Predicted minimum free energy (MFE), paired nucleotide fraction and structure statistics were summarized by window. Standardized folding energies were defined relative to a matched null ensemble; positive Z_MFE_-scores denote less favorable folding than the corresponding controls.

Inferences are computational predictions rather than experimental structural determinations:

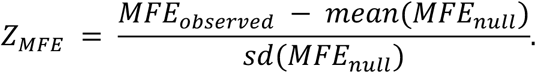

## Data availability

All sequence data have been deposited at NCBI SRA with accession #PRJNA1061599.

## End Notes Acknowledgements

Generous support was received from Edward Via College of Osteopathic Medicine and Center for Applied Immunology and Pathological Processes at Louisiana State University. We also thank Derek Walsh of Northwestern University Feinberg School of Medicine for providing reagents.

## Author Contributions

Conceptualization: P.M., S.D.

Data curation: K.M., L.K., S.D., P.M.

Formal analysis: K.M., L.K., S.D., P.M.

Investigation: A.F., K.M., S.W., A.R., E.V., T.M., L.K., S.D., P.M.

Methodology: K.M., P.M., S.D., L.K. Writing – original draft: P.M.

Writing – review & editing: A.F., K.M., L.K., S.D., P.M. Funding acquisition: P.M., S.D.

Resources: P.M., S.D., L.K. Supervision: P.M.

