## Supplementary Information for "Eukaryotic 45S ribosomal RNA spacers encode an ancient layer of antiviral response"

Supplementary Tables S1–S45. Table values are transferred from existing project analyses; no new numerical estimates were generated for this compilation.

### Supplementary Table S1. Host-clade and DNA-virus-family enrichment of ITS sequence similarity

| ITS | Viral family / host clade | Enrichment | Fisher p |
|---|---|---|---|
| ITS1 | Orthoherpes / Vertebrata | 28.2 | 5.29 10{yv |
| ITS2 | Orthoherpes / Vertebrata | 331.8 | 3.18 10{ p |
| ITS1 | Nudiviridae / Arthropoda | 25.3 | 9.40 10{tw |
| ITS2 | Nudiviridae / Arthropoda | 800.6 | 9.38 10{v |
| ITS1 | Baculoviridae / Arthropoda | 21.0 | 1.14 10{tx |
| ITS2 | Draupnirviridae / Fungi | 90.8 | <10{ pp |
| ITS2 | Malacoherpes / Arthropoda | other clades 0 | 2.58 10{vv |

**Enrichment estimates and Fisher exact probabilities for selected host–viral family combinations. Results are taken from the convergence analysis; reported values are not recalculated. Source: ITS_DNA_virus_convergence_MS_rev2_Results_Methods.docx, table 1.**

### Supplementary Table S2. Summary statistics for the ITS–DNA-virus homology screen

| Quantity | Estimate |
|---|---|
| Nuclear input genomes | 4,896 |
| Cytoplasmic input genomes | 307 |
| Nuclear retained HSPs | 11,686 |
| Cytoplasmic retained HSPs | 684 |
| Poisson nuclear IRR | 5.239 |
| HC3 95% CI | 1.778 15.438 |
| HC3 robust p | 0.002666 |

**Principal quantitative outputs of the comparative homology screen. The units and denominators follow the source analysis. Source: ITS_DNA_virus_convergence_MS_rev2_Results_Methods.docx, table 2.**

### Supplementary Table S3. Illustrative vertebrate ITS–orthoherpesvirus convergence cases

| Host | Observation | Caution |
|---|---|---|
| Siganus spinus | HSV-2 locus score 141; congener max 88 | No transfer junction |
| Anolis garmani | 96 nt; 14 HSPs; 8 viruses | 87.5% GC |
| Anolis cristatellus | 119 nt; 18 HSPs; 7 viruses | 93.3% GC |
| Anas | 64/81 unique bases match NC_043054 | Repeat-associated motifs |
| Pongo | 107/125 identities near viral 137 kb | Local shuffle p = 0.0099 |

**Case-specific observations and limitations for selected vertebrate ITS loci. Source: ITS_DNA_virus_convergence_MS_rev2_Results_Methods.docx, table 3.**

### Supplementary Table S4. Comparative ITS1 and ITS2 sequence-similarity measurements

| Measurement | ITS1 | ITS2 |
|---|---|---|
| Isolated window n | 67 | 33 |
| Median isolated MFE Z | +0.940 | +1.524 |
| Full-length window n | 58 | 29 |
| Mean paired-fraction difference | +0.0432 | +0.0221 |
| Positive paired-fraction differences | 41/58 | 22/29 |
| Two-sided sign-test p | 0.00223 | 0.00813 |
| Mean GC mismatch | 0.0432 | 0.0631 |

**Spacer-specific summary metrics from the convergence analysis. Source: ITS_DNA_virus_convergence_MS_rev2_Results_Methods.docx, table 4.**

### Supplementary Table S5. Anolis ITS1 candidate loci and exact 12-nt viral match summaries

| Species | Locus start | Locus end | Locus length (nt) | GC content | Repeat-8 fraction | Viral accession | HSPs | Exact 12-mers | Distinctive 12-mers |
|---|---|---|---|---|---|---|---|---|---|
| Anolis garmani | 226 | 321 | 96 | 0.875 | 0.667 | NC_063268 | 4 | 6 | 0 |
| Anolis garmani | 226 | 321 | 96 | 0.875 | 0.667 | NC_075564 | 1 | 4 | 0 |
| Anolis garmani | 226 | 321 | 96 | 0.875 | 0.667 | NC_043054 | 1 | 4 | 0 |
| Anolis garmani | 226 | 321 | 96 | 0.875 | 0.667 | NC_075563 | 1 | 0 | 0 |
| Anolis garmani | 226 | 321 | 96 | 0.875 | 0.667 | NC_007646 | 1 | 4 | 0 |
| Anolis garmani | 226 | 321 | 96 | 0.875 | 0.667 | NC_076509 | 3 | 1 | 0 |
| Anolis garmani | 226 | 321 | 96 | 0.875 | 0.667 | NC_014567 | 2 | 2 | 0 |
| Anolis garmani | 226 | 321 | 96 | 0.875 | 0.667 | NC_055561 | 1 | 0 | 0 |
| Anolis trachyderma | 234 | 312 | 79 | 0.911 | 0.519 | — | — | — | — |
| Anolis cristatellus | 211 | 329 | 119 | 0.933 | 0.655 | NC_075563 | 3 | 11 | 0 |
| Anolis cristatellus | 211 | 329 | 119 | 0.933 | 0.655 | NC_014567 | 2 | 6 | 0 |

| | | | | | | | | | |
|---|---|---|---|---|---|---|---|---|---|
| Anolis cristatellus | 211 | 329 | 119 | 0.933 | 0.655 | NC_034446 | 2 | 4 | 0 |
| Anolis cristatellus | 211 | 329 | 119 | 0.933 | 0.655 | NC_055561 | 2 | 15 | 0 |
| Anolis cristatellus | 211 | 329 | 119 | 0.933 | 0.655 | NC_001798 | 6 | 3 | 0 |
| Anolis cristatellus | 211 | 329 | 119 | 0.933 | 0.655 | NC_043054 | 1 | 8 | 0 |
| Anolis cristatellus | 211 | 329 | 119 | 0.933 | 0.655 | NC_063268 | 2 | 15 | 0 |
| Anolis grahami | 196 | 319 | 124 | 0.855 | 0.516 | NC_063268 | 4 | 6 | 0 |
| Anolis grahami | 196 | 319 | 124 | 0.855 | 0.516 | NC_075564 | 1 | 4 | 0 |
| Anolis grahami | 196 | 319 | 124 | 0.855 | 0.516 | NC_043054 | 1 | 4 | 0 |
| Anolis grahami | 196 | 319 | 124 | 0.855 | 0.516 | NC_075563 | 3 | 8 | 0 |
| Anolis grahami | 196 | 319 | 124 | 0.855 | 0.516 | NC_007646 | 1 | 4 | 0 |
| Anolis grahami | 196 | 319 | 124 | 0.855 | 0.516 | NC_076509 | 3 | 1 | 0 |
| Anolis grahami | 196 | 319 | 124 | 0.855 | 0.516 | NC_014567 | 2 | 2 | 0 |
| Anolis grahami | 196 | 319 | 124 | 0.855 | 0.516 | NC_055561 | 1 | 0 | 0 |
| Anolis capito | 280 | 339 | 60 | 0.85 | 0.333 | — | — | — | — |
| Anolis nebuloides | 216 | 307 | 92 | 0.891 | 0.457 | — | — | — | — |
| Anolis cupreus | 239 | 307 | 69 | 0.928 | 0.522 | — | — | — | — |

**Source: frozen Anolis_candidate_summary.tsv and Anolis_distinctive_viral_12mers.tsv on bigmem. Exact 12-mer counts describe sequence-level matching, not a reconstructed base-by-base alignment. Repeat-rich matches require cautious interpretation.**

### Supplementary Table S6. Anas ITS1–orthoherpesvirus local alignment identity partitioned by repeat status

| Viral accession | Alignment score | Nonrepeat aligned bases | Nonrepeat identical bases | Repeat aligned bases | Repeat identical bases | Exact 12-mers | Exact 12-mers in unique sequence |
|---|---|---|---|---|---|---|---|
| NC_043054 | 142 | 81 | 64 | 59 | 55 | 18 | 1 |
| NC_005261 | 128 | 75 | 57 | 60 | 53 | 27 | 0 |

| | | | | | | | |
|---|---|---|---|---|---|---|---|
| NC_020231 | 104 | 66 | 45 | 60 | 56 | 26 | 0 |
| NC_063268 | 113 | 77 | 54 | 57 | 51 | 17 | 0 |
| NC_002794 | 108 | 59 | 45 | 59 | 50 | 31 | 0 |
| NC_055561 | 130 | 66 | 55 | 59 | 52 | 1 | 0 |
| NC_055166 | 120 | 66 | 54 | 59 | 49 | 23 | 4 |
| NC_075563 | 125 | 61 | 49 | 59 | 52 | 21 | 0 |
| NC_076509 | 107 | 67 | 54 | 59 | 53 | 17 | 0 |

**Source: frozen Anas_1567_1710_distinctive_viral_support.tsv on bigmem. Values are alignment summaries for ITS1 positions 1567–1710; unique and repeated segments are reported separately. No unverified nucleotide alignments were reconstructed.**

### Supplementary Table S7. Primate ITS1 alignments converging on a shared orthoherpesvirus genomic interval

| Genus | Host ITS accession | Alignment score | ITS1 start | ITS1 end | Viral start | Viral end | Aligned nucleotide pairs | Identical pairs | Identity | ITS1 GC |
|---|---|---|---|---|---|---|---|---|---|---|
| Homo | ITS1DB01121385 | 247 | 406 | 1231 | 136647 | 137435 | 759 | 514 | 0.6772 | 0.7475 |
| Pongo | ITS1DB00167964 | 282 | 231 | 1029 | 136636 | 137446 | 755 | 523 | 0.6927 | 0.7612 |
| Macaca | ITS1DB00801366 | 224 | 339 | 1004 | 136639 | 137292 | 621 | 425 | 0.6844 | 0.7415 |
| Pan | ITS1DB00356819 | 65 | 799 | 1016 | 137023 | 137253 | 214 | 139 | 0.6495 | 0.6238 |
| Gorilla | ITS1DB00027738 | 235 | 179 | 975 | 136618 | 137443 | 767 | 509 | 0.6636 | 0.7479 |
| Callithrix | ITS1DB00136452 | 175 | 289 | 959 | 136643 | 137274 | 604 | 417 | 0.6904 | 0.7162 |
| Nomascus | ITS1DB00801364 | 265 | 363 | 1214 | 136643 | 137435 | 765 | 535 | 0.6993 | 0.7495 |

**Source: frozen ITS1_orthoherpes_primate_7genera_fullterminal_summary.tsv on bigmem. Coordinates provide locus-resolved alignment examples; identity and pair counts are observed, not inferred. Convergence at one viral neighborhood does not by itself establish historical sequence transfer.**

### Supplementary Table S8. RNA secondary-structure stability of orthoherpesvirus-matching vertebrate ITS regions

| Measurement | ITS1 | ITS2 |
|---|---|---|
| Isolated window n | 67 | 33 |
| Median isolated MFE Z | +0.940 | +1.524 |
| Full-length window n | 58 | 29 |
| Mean paired-fraction difference | +0.0432 | +0.0221 |
| Positive paired-fraction differences | 41/58 | 22/29 |
| Two-sided sign-test p | 0.00223 | 0.00813 |
| Mean GC mismatch | 0.0432 | 0.0631 |

**Source: ITS_DNA_virus_convergence_MS_rev2_Results_Methods.docx, fourth table. Original numerical entries and column labels retained; values summarize isolated and full-spacer RNA folding analyses and corresponding null-standardized MFE statistics.**

### Supplementary Table S9. Gammaherpesvirus RNA-seq datasets and experimental contrasts

| Dataset | System/design | Primary molecular fraction | Key ITS result | Mechanistic leverage |
|---|---|---|---|---|
| GSE70481 MHV68 | NIH 3T3; mock, WT, ΔHS; 24 h; n=2/condition | 4sU-labeled, Ribo-Zero RNA | WT ITS1 0.182×; ITS2 0.179×; total45S 3.216×. ΔHS rescues ITS1 to 0.627× and ITS2 to 0.868×. | muSOX R443I host-shutoff mutant directly tests dependence on viral host shutoff. |
| GSE165328 EBV | Primary B cells; EBV 2089 vs matched mock; donors 22/25/26; 10–12 dpi | Ribo-Zero total RNA | total45S 1.793×; ITS1 1.930×; ITS2 1.516×. ITS2/ITS1 0.727× (alignment), 0.713× (unique QNAME), 0.771× (31-mer). | True matched human donor controls; very deep sequencing; orthogonal counting methods. |
| E-MTAB-7805 EBV | Naïve primary B cells; 3 donors; day 0,1,2,3,4,5,8,14 | Poly(A)-selected RNA | ITS2/ITS1 rises in 21/21 donor×post-baseline comparisons; mean +0.929 log2 (~1.90×) by both alignment and 31-mers. | Time resolution through EBV-driven B-cell reprogramming; independent validation of non-stoichiometric spacer behavior. |

**Design, RNA fraction and principal interpretation of the MHV68 and EBV datasets analyzed. Source: Gammaherpesvirus_MHV68_EBV_Methods_Results_CELL.docx, table 1.**

### Supplementary Table S10. Gammaherpesvirus-associated changes across the 45S transcription unit

| Virus/condition | 18S FC | ITS1 FC | 5.8S FC | ITS2 FC | 28S FC | total45S FC | ΔITS1 | ΔITS2 | D |
|---|---|---|---|---|---|---|---|---|---|
| MHV68 WT | 3.650 | 0.182 | 0.226 | 0.179 | 3.200 | 3.216 | −4.146 | −4.168 | −0.022 |
| MHV68 ΔHS | 0.843 | 0.627 | 0.409 | 0.868 | 0.936 | 0.907 | −0.532 | −0.064 | +0.469 |
| EBV matched donors | 2.289 | 1.930 | 3.982 | 1.516 | 1.969 | 1.793 | +0.106 | −0.243 | −0.349 |

**Infected-to-mock regional fold changes and spacer-specific contrasts. FC, fold change; ITS, internal transcribed spacer. Source: Gammaherpesvirus_MHV68_EBV_Methods_Results_CELL.docx, table 2.**

### Supplementary Table S11. MHV68 sequence-similarity versus ITS positional response

| Condition | Scope | β | β relevant-tail p | λ | λ upper-tail p | Interpretation |
|---|---|---|---|---|---|---|
| WT | ITS1+ITS2 | −0.143 | 0.153 | +0.118 | 0.194 | Weak graded depletion/extremeness tendency; spatial null not significant |
| WT | ITS1 | −0.260 | — | +0.224 | — | Combined WT tendency concentrated in ITS1 |
| WT | ITS2 | +0.035 | — | −0.035 | — | No positive extremeness association |
| ΔHS | ITS1+ITS2 | −0.122 | 0.201 | −0.038 | 0.625 | No extremeness association |
| ΔHS | ITS1 | −0.215 | — | −0.141 | — | No positive extremeness association |
| ΔHS | ITS2 | +0.001 | — | +0.058 | — | Weak/near-null |

**Signed and magnitude-based correlations between viral sequence similarity and ITS response, with empirical significance estimates. Source: MHV68_ITS_rDNA_Methods_Results.docx, table 3.**

### Supplementary Table S12. MHV68 spatial null tests and permutation diagnostics

| Contrast / scope | Statistic | Observed | Null mean | Empirical p | Approx. rank | Interpretation |
|---|---|---|---|---|---|---|

| | | | | | | |
|---|---|---|---|---|---|---|
| WT, combined | λ | +0.118 | −0.075 | 0.0693 | 7th highest / 101 | Real WUMS more aligned with response than ~94% of pseudo-genomes |
| WT, combined | β | −0.143 | +0.072 | 0.0693 | 7th most negative / 101 | Real WUMS unusually aligned with depletion |
| WT, ITS1 | λ | +0.224 | −0.061 | 0.0792 | — | Sequence-specific tendency concentrated in ITS1 |
| WT, ITS1 | β | −0.260 | — | 0.0594 | — | Strongest directional composition-null tendency |
| WT, ITS2 | λ | −0.035 | −0.055 | 0.465 | — | No sequence-specific signal |
| WT−ΔHS, combined | Δλ | +0.156 | −0.065 | 0.0693 | 7th highest / 101 | WT-specific alignment unusual but not conventionally significant |
| WT−ΔHS, ITS1 | Δλ | +0.364 | ~0 | 0.0693 | 7th highest / 101 | Mechanistic contrast again concentrated in ITS1 |
| WT−ΔHS, ITS2 | Δλ | −0.092 | −0.075 | 0.505 | ~51st / 101 | No contrast |

**Observed spatial statistics compared with null distributions and empirical probabilities. Source: MHV68_ITS_rDNA_Methods_Results.docx, table 4.**

### Supplementary Table S13. MCMV infection time-course sampling design

| Condition | Time (hpi) | Biological replicates | Primary role |
|---|---|---|---|
| Mock | 0 | 2 | Baseline for all fold changes |
| MCMV | 1, 2, 4, 6, 12, 18, 24, 36, 48, 72 | 2 at each time | Replicated infection time course |
| MCMV + PAA | 24 | 1 | Exploratory viral-DNA-replication perturbation |
| MCMV + CYC | 4 | 1 | Exploratory translation perturbation |

**Experimental conditions and biological replication for the MCMV NIH-3T3 RNA-seq analysis. Source: MCMV_ITS_RNAseq_Methods_Results_Cell_style.docx, table 1.**

### Supplementary Table S14. Mouse rDNA segment coordinates used for regional quantification

| Region | Start | End | Length (nt) |
|---|---|---|---|
| 18S | 4008 | 5877 | 1,870 |
| ITS1 | 5878 | 6877 | 1,000 |
| 5.8S | 6878 | 7034 | 157 |
| ITS2 | 7035 | 8122 | 1,088 |
| 28S | 8123 | 12852 | 4,730 |
| Transcribed 45S | 4008 | 12852 | 8,845 |

**Region boundaries in the mouse rDNA reference used for coverage and k-mer analyses. Source: MCMV_ITS_RNAseq_Methods_Results_Cell_style.docx, table 2.**

### Supplementary Table S15. Time-resolved MCMV effects on ITS1, ITS2 and 45S transcription

| hpi | ITS1 FC | ITS2 FC | D (log2) | 45S FC | Descriptive pattern |
|---|---|---|---|---|---|
| 1 | 0.888 | 0.762 | -0.221 | 0.847 | ITS2 preferential depletion |

| | | | | | |
|---|---|---|---|---|---|
| 2 | 0.899 | 0.918 | +0.030 | 0.928 | Near-equivalent |
| 4 | 1.030 | 1.316 | +0.354 | 1.045 | ITS2 relative enrichment |
| 6 | 0.969 | 1.229 | +0.344 | 1.003 | ITS2 relative enrichment |
| 12 | 0.918 | 0.793 | -0.211 | 0.903 | ITS2 preferential depletion |
| 18 | 0.879 | 0.741 | -0.247 | 0.867 | ITS2 preferential depletion |
| 24 | 0.808 | 0.799 | -0.016 | 0.802 | Near-equivalent |
| 36 | 1.034 | 0.700 | -0.562 | 0.910 | Strong ITS2 preferential depletion |
| 48 | 0.899 | 0.627 | -0.519 | 0.831 | Strong ITS2 preferential depletion |
| 72 | 0.946 | 0.789 | -0.261 | 0.889 | ITS2 preferential depletion |

**Regional fold changes and ITS2-versus-ITS1 response differences across the infection time course. Source: MCMV_ITS_RNAseq_Methods_Results_Cell_style.docx, table 3.**

### Supplementary Table S16. Concordance of MCMV alignment-depth and exact 31-mer estimates

| hpi | 31-mer ITS1 FC | 31-mer ITS2 FC | 31-mer D | Alignment D | Method agreement |
|---|---|---|---|---|---|
| 1 | 1.214 | 1.022 | -0.248 | -0.221 | Concordant negative |
| 2 | 1.134 | 0.951 | -0.253 | +0.030 | Sign discordant |
| 4 | 0.926 | 0.885 | -0.066 | +0.354 | Sign discordant |
| 6 | 1.188 | 1.101 | -0.110 | +0.344 | Sign discordant |
| 12 | 1.032 | 1.399 | +0.438 | -0.211 | Sign discordant |
| 18 | 1.269 | 1.459 | +0.201 | -0.247 | Sign discordant |
| 24 | 1.669 | 1.935 | +0.213 | -0.016 | Weak/near-zero alignment |
| 36 | 1.087 | 1.510 | +0.474 | -0.562 | Strongly discordant |
| 48 | 0.632 | 0.809 | +0.356 | -0.519 | Strongly discordant |
| 72 | 0.749 | 2.098 | +1.485 | -0.261 | Strongly discordant |

**Comparison of two independent rDNA quantification strategies across the MCMV time course. Source: MCMV_ITS_RNAseq_Methods_Results_Cell_style.docx, table 4.**

### Supplementary Table S17. MCMV–mouse ITS local sequence-similarity candidates

| Spacer | Raw HSP | Length | Identity | E value | Bit score | >=20 nt retained? |
|---|---|---|---|---|---|---|
| ITS1 | CCGCGGCGGCGGCTGCT | 17 nt | 100% | 0.012 | 34.2 | No |
| ITS2 | CGCGCATCCGCGCTCGC | 17 nt | 100% | 0.013 | 34.2 | No |

**Alignment characteristics of candidate MCMV–ITS homologies, including retention criteria. Source: MCMV_ITS_RNAseq_Methods_Results_Cell_style.docx, table 5.**

### Supplementary Table S18. HCMV RNA-seq cohorts and experimental context

| Dataset | Biological system | HCMV context | Sampling / RNA fraction | Primary value | Principal limitation |
|---|---|---|---|---|---|
| GSE164839 | Kasumi-3 | TB40/E wtGFP, MOI 1 | ~24 h; Ribo-Zero total RNA; n=2/group | Deep 45S, spatial, eRL, homology and 31-mer analysis | Very small n; infected-replicate heterogeneity |

| | | | | | |
|---|---|---|---|---|---|
| GSE164843 | Kasumi-3 | HCMV latency model | Nascent and steady-state RNA | Direct processing-stage contrast | Different RNA fractions are not interchangeable biological replicates |
| GSE210989 | MRC-5 | AD169/BADwt-like infected libraries | ~48 h; n=3/group | Independent intermediate-time evidence; broad 45S regional profile | Condition-by-batch confounding; qualified evidence |
| GSE99454 | MRC-5 and ARPE-19 | TB40/E | 24, 72, 120 h; n=2/group/context | Temporal directionality and cell-type replication | Small n per context; major mature-rRNA remodeling |

**Dataset-level descriptions, infection conditions, sampling and limitations. Source: HCMV_cross_dataset_45S_ITS1_ITS2_Cell_Methods_Results.docx, table 1.**

### Supplementary Table S19. HCMV infection-associated regional rDNA fold changes

| Dataset / context | 18S | ITS1 | 5.8S | ITS2 | 28S | eRL | Interpretive point |
|---|---|---|---|---|---|---|---|
| GSE164839, Kasumi-3 ~24 h | 11.33x | 3.65x | 4.88x | 11.59x | 6.95x | 10.99x | Large, broad 45S increase; ITS2/eRL prominent but not isolated |
| GSE164843, nascent | 0.72x | 1.64x | 1.36x | 1.65x | 0.89x | 2.32x | Spacer/eRL enrichment in newly synthesized/processed RNA despite mature-rRNA neutrality/downshift |
| GSE164843, steady | 1.10x | 1.20x | 0.99x | 1.04x | 1.00x | ~1.10x* | Near-neutral steady-state 45S response |
| GSE210989, MRC-5 ~48 h | 1.17x | 0.84x | 1.43x | 0.88x | 0.96x | 2.76x | ITS2 mildly depleted while eRL sequence abundance is elevated; qualified because of batch confounding |
| GSE99454, MRC-5 24 h | — | 2.10x | — | 1.16x | — | — | Early ITS2 near neutral/up, but ITS1 already elevated |
| GSE99454, MRC-5 120 h | — | 0.81x | — | 0.46x | — | — | Late ITS2 depletion stronger than ITS1 |
| GSE99454, ARPE-19 24 h | — | 1.63x | — | 1.73x | — | — | Early concordant ITS1/ITS2 increase |
| GSE99454, ARPE-19 120 h | — | 1.58x | — | 0.62x | — | — | Late divergence: ITS2 depleted while ITS1 remains elevated |

**Cross-context estimates for mature-rRNA and ITS segments of the 45S transcript. Source: HCMV_cross_dataset_45S_ITS1_ITS2_Cell_Methods_Results.docx, table 2.**

### Supplementary Table S20. Integrated HCMV ITS and rDNA responses across studies

| Feature | Studies | Median log2 effect | Integrated FC | 95% interval (FC) | P(effect > 0) | Biological reading |
|---|---|---|---|---|---|---|
| ITS1 | 4 | 0.421 | 1.34 | 0.96-1.82 | 0.958 | Consistent tendency toward increased ITS1 representation |
| ITS2 | 5 | 0.491 | 1.40 | 0.89-2.76 | 0.912 | Positive overall tendency with substantial temporal/context heterogeneity |

| | | | | | | |
|---|---|---|---|---|---|---|
| eRL | 4 | 0.826 | 1.77 | 1.01-3.49 | 0.978 | Strongest integrated directional signal; nested within ITS2 |
| ITS2/ITS1 | 4 | 0.036 | 1.03 | 0.68-1.65 | 0.563 | No evidence for universal ITS2 selectivity over ITS1 |
| ITS2/28S | 2 | 0.856 | 1.81 | 0.95-5.33 | 0.750 | Suggestive relative spacer enrichment, but sparse and sensitive to mature-rRNA remodeling |

**Cross-study effects and uncertainty summaries as reported in the source analysis. Source: HCMV_cross_dataset_45S_ITS1_ITS2_Cell_Methods_Results.docx, table 3.**

### Supplementary Table S21. Leave-one-dataset-out sensitivity of the HCMV ITS response

| Excluded dataset | Datasets remaining | Mean log2FC | FC |
|---|---|---|---|
| None | 5 | 0.838 | 1.79 |
| GSE124118 | 4 | 0.846 | 1.80 |
| GSE164839 | 4 | 0.164 | 1.12 |
| GSE164843 | 4 | 0.951 | 1.93 |
| GSE210989 | 4 | 1.094 | 2.14 |
| GSE99454 | 4 | 1.137 | 2.20 |

**Sensitivity of integrated estimates to exclusion of individual HCMV datasets. Source: HCMV_cross_dataset_45S_ITS1_ITS2_Cell_Methods_Results.docx, table 4.**

### Supplementary Table S22. Temporal HCMV responses in MRC-5 and ARPE-19 cells

| Cell / region | 24 h | 72 h | 120 h | Temporal summary |
|---|---|---|---|---|
| MRC-5 whole ITS2 | 1.16x | 0.74x | 0.46x | Slope -0.545 log2FC/time unit; R2=0.954 |
| ARPE-19 whole ITS2 | 1.73x | 0.54x | 0.62x | Slope -0.705; R2=0.827 |
| Window 7633-7732, pooled cells | early log2FC +0.552 | late mean -0.863 | — | Early-minus-late = 1.415 log2 units |
| Window 7233-7332, pooled cells | early +0.173 | late -1.120 | — | Early-minus-late = 1.293 |
| Window 6933-7032, pooled cells | early +0.275 | late -1.021 | — | Early-minus-late = 1.297 |

**Cell-specific ITS/rDNA response trajectories at 24, 72 and 120 hours post-infection. Source: HCMV_cross_dataset_45S_ITS1_ITS2_Cell_Methods_Results.docx, table 5.**

### Supplementary Table S23. Spatial localization of recurrent HCMV-associated ITS responses

| Region | 45S coordinates | Datasets | Median FC | Positive datasets | Interpretation |
|---|---|---|---|---|---|
| ITS1 | 5706-5805 | 4 | 2.00x | 3/4 | Recurrent proximal ITS1 increase |
| ITS1 | 6006-6105 | 4 | 1.90x | 3/4 | Second recurrent ITS1 domain |
| ITS1 | 6106-6205 | 4 | 1.68x | 3/4 | Broadens ITS1 response beyond one hotspot |
| ITS2 | 7133-7232 | 5 | 1.52x | 3/5 | eRL-overlapping region represented in every dataset |
| ITS2 | 7533-7632 | 3 | 2.87x | 3/3 | Strong distal ITS2 domain |

| | | | | | |
|---|---|---|---|---|---|
| ITS2 | 7433-7532 | 3 | 1.58x | 3/3 | Neighboring distal ITS2 support |
| ITS2 | 7633-7732 | 3 | 2.42x | 2/3 | Strong magnitude but one discordant dataset |

**Regional positions and cross-dataset recurrence of transcriptional responses along the 45S unit. Source: HCMV_cross_dataset_45S_ITS1_ITS2_Cell_Methods_Results.docx, table 6.**

### Supplementary Table S24. Recurrent HCMV-associated exact 31-mer clusters

| Region | 45S cluster | Overlapping recurrent 31-mers | Median log2 effect | Median FC | Biological interpretation |
|---|---|---|---|---|---|
| ITS2 | 7207-7266 | 10 | 1.240 | 2.36x | eRL-centered recurrent sequence domain |
| ITS1 | 5666-5711 | 11 | 1.026 | 2.04x | Largest recurrent ITS1 sequence cluster |
| ITS1 | 5975-6008 | 4 | 1.251 | 2.38x | Highest-magnitude ITS1 cluster |
| ITS1 | 5624-5660 | 3 | 1.032 | 2.04x | Additional proximal ITS1 cluster |

**Local sequence clusters repeatedly implicated across HCMV datasets. Source: HCMV_cross_dataset_45S_ITS1_ITS2_Cell_Methods_Results.docx, table 7.**

### Supplementary Table S25. HCMV ITS2 sequence-similarity interactions across datasets

| Dataset | Contexts | Mean delta_homology (log2) | Interaction FC | Direction pattern | Interpretation |
|---|---|---|---|---|---|
| GSE124118 | 4 | -0.548 | 0.684 | 0 positive / 4 negative | Homologous sequence relatively depleted |
| GSE164839 | 1 | -0.003 | 0.998 | Near zero | Large ITS2 increase is not explained by HCMV homology |
| GSE164843 | 2 | +0.381 | 1.302 | 1 positive / 1 negative | Nascent-specific homology enrichment; steady ~neutral |
| GSE99454 | 6 | -0.271 | 0.829 | 0 positive / 6 negative | Consistent relative depletion of homologous ITS2 |
| GSE210989 | 1 | ~ -0.212 | ~0.864 | Negative | Independent qualified evidence; AD169 context and batch confounding |

**Dataset-level relationships between HCMV sequence similarity and ITS2 transcriptional response. Source: HCMV_cross_dataset_45S_ITS1_ITS2_Cell_Methods_Results.docx, table 8.**

### Supplementary Table S26. HSV-1 RNA-seq datasets and their complementary experimental designs

| Dataset | Design / principal value | Role in integration |
|---|---|---|
| GSE59717 / GSM1444166 | HFF, HSV-1 strain 17, MOI 10; 4sU 1-h pulses, 0–8 h; 2 trajectories; no rRNA depletion | Primary nascent 45S kinetics and spacer attrition |
| GSE103763 | KMB17, strain 17, MOI 1, 48 h; 3+3 | Independent late abundance, windows, homology |
| GSE124118 | HFF-1/MRC-5; MOI 0.5; mock, 4 h, 8 h; triplicates | Early abundance, cell-type kinetics, read localization |
| GSE100576/GSE129582 | HFF, strain 17, MOI 10, 8 h; Total/Cyto/Nuc/Chrom; n=2; WT and Δvhs | Subcellular flux and vhs perturbation |
| GSE73853 | HFF, HSV-2 strain G, MOI 10, 8 h; n=1/condition | Cross-herpesvirus descriptive complement |

Overview of the HSV-1 time-course, endpoint and fractionation datasets used for integrative inference. Source: HSV1_ITS_Integrated_Methods_Results_Cell_style.docx, table 1.

### Supplementary Table S27. Nascent HSV-1 45S, ITS1 and ITS2 RNA trajectories

| 4sU interval | Total 45S FC | ITS1 FC | ITS2 FC | ITS2/45S relative FC |
|---|---|---|---|---|
| 0–1 | 1.06 | 1.06 | 1.02 | 0.964 |
| 1–2 | 1.04 | 0.92 | 0.98 | 0.943 |
| 2–3 | 1.02 | 0.76 | 0.74 | 0.729 |
| 3–4 | 1.03 | 0.61 | 0.54 | 0.520 |
| 4–5 | 0.96 | 0.65 | 0.52 | 0.545 |
| 5–6 | 1.05 | 0.75 | 0.61 | 0.582 |
| 6–7 | 0.92 | 0.84 | 0.59 | 0.639 |
| 7–8 | 0.82 | 0.70 | 0.46 | 0.556 |

Regional infected-to-mock fold changes during consecutive 4sU labeling intervals. Source: HSV1_ITS_Integrated_Methods_Results_Cell_style.docx, table 2.

### Supplementary Table S28. HSV-1 ITS2 responses across experimental contexts

| Context | ITS2 depth FC | Bayesian median (95% CrI) |
|---|---|---|
| GSE103763 KMB17 48 h | 1.589× | 1.644× (0.828–3.251) |
| GSE124118 HFF-1 4 h | ~2.72× | 2.74× (1.91–3.95) |
| GSE124118 HFF-1 8 h | ~2.26× | 2.28× (1.54–3.36) |
| GSE124118 MRC-5 4 h | ~0.89× | 0.94× (0.50–1.77) |
| GSE124118 MRC-5 8 h | ~1.54× | 1.60× (0.80–3.18) |

ITS2 depth and uncertainty estimates across HSV-1 infection datasets. Source: HSV1_ITS_Integrated_Methods_Results_Cell_style.docx, table 3.

### Supplementary Table S29. Subcellular fraction-specific HSV-1 ITS2 remodeling

| Fraction | ITS2/45S GM | Raw31 ITS2/ITS1 GM | Interpretation |
|---|---|---|---|
| Total | 2.664 | 0.891 | Expanded pool with ITS imbalance |
| Cytoplasmic | 1.740 | 0.291 | Strong relative ITS2 attrition |
| Nucleoplasmic | 1.963 | 0.390 | Production/processing pool with separation |
| Chromatin | 1.378 | 0.918 | Near-source accumulation/preservation |

Comparison of ITS2 responses in distinct RNA fractions. Source: HSV1_ITS_Integrated_Methods_Results_Cell_style.docx, table 4.

### Supplementary Table S30. HSV-1 KOS small-RNA ITS and rRNA fragment responses

| Feature | HSV/mock fold CPM | log2FC | Exact P | BH q | HSV/mock fold within 45S | Interpretation |
|---|---|---|---|---|---|---|

| | | | | | | |
|---|---|---|---|---|---|---|
| All transcribed 45S | 0.904 | −0.146 | 0.314 | 0.471 | 1.000 | No global collapse |
| 5′ ETS | 0.237 | −2.076 | 0.0286 | 0.0857 | 0.266 | Strong depletion |
| 18S | 1.882 | +0.913 | 0.0286 | 0.0857 | 2.060 | Strong enrichment |
| ITS1 | 1.023 | +0.033 | 0.857 | 0.935 | 1.146 | Essentially unchanged |
| 5.8S | 1.179 | +0.238 | 0.171 | 0.343 | 1.308 | Compositional enrichment |
| ITS2 | 0.460 | −1.120 | 0.0286 | 0.0857 | 0.504 | Selective ~2-fold depletion |
| 28S | 0.869 | −0.203 | 0.200 | 0.343 | 0.960 | Modest/no global change |
| 5.8S→ITS2 | 1.042 | +0.059 | 0.971 | 0.971 | 1.197 | No depletion |
| ITS2→28S | 0.419 | −1.255 | 0.0286 | 0.0857 | 0.457 | Strong 3′-ITS2 boundary depletion |

**Regional small-RNA abundance contrasts in infected versus mock-treated NHDFs, including multiplicity-adjusted results. Source: HSV1_KOS_smRNA_ITS_fragmentome_Methods_Results_REMADE.docx, table 1.**

### Supplementary Table S31. HSV-1 KOS spatial association of ITS sequence similarity and transcription

| Region | β signed | P, two-sided | λ response magnitude | P, upper-tail | Interpretation |
|---|---|---|---|---|---|
| ITS1+ITS2 | −0.1058 | 0.1773 | 0.1388 | 0.00310 | KOS-like sequence marks response-prone positions |
| ITS1 | −0.1829 | 0.1217 | 0.2083 | 0.00210 | Strongest and localized to 5′ ITS1 |
| ITS2 | +0.0351 | 0.6596 | 0.0462 | 0.2127 | No ITS2-wide homology coupling |

**Signed and magnitude-based positional correlation statistics for the small-RNA experiment. Source: HSV1_KOS_smRNA_ITS_fragmentome_Methods_Results_REMADE.docx, table 3.**

### Supplementary Table S32. HSV-1 dataset-specific ITS2 sequence-similarity response interactions

| Dataset / context | Δ_homology (log2) | Interaction FC | Directional P | Bayesian P(Δ<0) |
|---|---|---|---|---|
| GSE103763 | −0.428 | 0.743× | 0.05 | 0.972 (frozen analysis) |
| GSE124118 HFF-1 4 h | −0.334 | 0.793× | 0.05 | 0.991 |
| GSE124118 HFF-1 8 h | −0.471 | 0.722× | 0.05 | 0.998 |
| GSE124118 MRC-5 4 h | −0.530 | 0.693× | 0.15 | 0.917 |
| GSE124118 MRC-5 8 h | −0.862 | 0.550× | 0.05 | 0.976 |

**Cross-context association estimates and directional probabilities for ITS2 homology and expression. Source: ITS2_HSV_GSE103763_GSE124118_Cell_Methods_Results.docx, table 7.**

### Supplementary Table S33. HSV-1 sequence-block positional response sensitivity

| Block | Context | Absolute FC | Adjusted residual (log2) | Empirical lower-tail P | Interpretation |
|---|---|---|---|---|---|
| H3 | HFF-1 4 h | 1.85× | −1.148 | 0.006 | Strong relative attenuation |
| H3 | HFF-1 8 h | 1.17× | −1.594 | 0.006 | Strong relative attenuation |
| H3 | MRC-5 4 h | 0.47× | −1.128 | 0.006 | Strong relative attenuation |

| | | | | | |
|---|---|---|---|---|---|
| H3 | MRC-5 8 h | 0.53× | −1.854 | 0.006 | Strong relative attenuation |
| H4 | HFF-1 4 h | 4.37× | −0.047 | 0.382 | Near covariate expectation |
| H4 | HFF-1 8 h | 4.31× | −0.028 | 0.383 | Near covariate expectation |
| H4 | MRC-5 4 h | 2.17× | +0.601 | 0.989 | Over-enriched, not attenuated |
| H4 | MRC-5 8 h | 2.55× | +0.533 | 1.000 | Over-enriched, not attenuated |

**Blockwise positional fold changes, adjusted residuals and empirical tail probabilities. Source: ITS2_HSV_GSE103763_GSE124118_Cell_Methods_Results.docx, table 8.**

### Supplementary Table S34. Canonical biological hypotheses M0–M3 for ITS dynamics

| Model | Core mechanism | Host ITS antiviral deployment | Explicit viral remodeling of rRNA/ITS kinetics | Primary role in comparison |
|---|---|---|---|---|
| M0 | Passive ribosome biogenesis | No | No | Strict null |
| M1 | Virus-driven remodeling without ITS function | No | Yes | Strong nonfunctional alternative |
| M2 | Infection-responsive host ITS deployment | Yes | Minimal/background only | Parsimonious functional model |
| M3 | Hybrid remodeling plus deployment | Yes | Yes | Most flexible and broadly biologically plausible alternative |

**Model definitions distinguish passive processing, virus-directed remodeling, host ITS deployment and their hybrid. Source: HSV_dynamic_modeling_manuscript.docx, table 1.**

### Supplementary Table S35. RNA-only dynamical model fit and information-criterion comparison

| Model | n | k | RSS | AICc | Delta AICc | Akaike weight |
|---|---|---|---|---|---|---|
| M0 | 24 | 3 | 2.8370 | -44.047 | 57.251 | 2.29 x 10^-13 |
| M1 | 24 | 6 | 0.18147 | -100.292 | 1.007 | 0.3744 |
| M2 | 24 | 5 | 0.20225 | -101.299 | 0 | 0.6194 |
| M3 | 24 | 8 | 0.17790 | -92.111 | 9.188 | 0.00626 |

**Residual fit, parameter counts, corrected Akaike information criterion and relative model support. Source: HSV_dynamic_modeling_manuscript.docx, table 2.**

### Supplementary Table S36. Bayesian evidence for HSV-1 RNA-only dynamical models

| Model | Laplace posterior probability, primary prior | Direct-IS log evidence, SD 1.5 | Direct-IS ESS | Direct-IS posterior probability |
|---|---|---|---|---|
| M0 | 3.07 x 10^-9 | -10.2823 | 15,574 | 2.05 x 10^-9 |
| M1 | 0.02316 | 6.1585 | 235 | 0.02831 |
| M2 | 0.95271 | 9.6677 | 7,458 | 0.94611 |
| M3 | 0.02413 | 6.0571 | 115 | 0.02558 |

**Laplace and direct importance-sampling estimates, effective sample sizes and posterior probabilities. Source: HSV_dynamic_modeling_manuscript.docx, table 3.**

### Supplementary Table S37. HSV-1 KOS plaque-size response to ITS2 perturbation

| Manipulation | Contrast | Median ratio | 95% CrI | P(direction) | P(R > 1.20) |
|---|---|---|---|---|---|
| Depletion | ITS2-U / pooled control | 1.553 | 1.396–1.726 | P(R>1)=1.000 | 1.000 |
| Depletion | ITS2-D / pooled control | 1.294 | 1.165–1.446 | P(R>1)=1.000 | 0.9186 |
| Enrichment | ITS2-U / pooled control | 0.621 | 0.521–0.741 | P(R<1)=1.000 | — |
| Enrichment | ITS2-D / pooled control | 0.630 | 0.530–0.754 | P(R<1)=1.000 | — |

**Posterior plaque-area effect estimates and directional probabilities from independent KOS experiments. Source: ITS2_HSV_KOS_plaque_size_Methods_Results_Bayesian.docx, table 2.**

### Supplementary Table S38. HSV-1 titer responses to ITS2 siRNA depletion

| Strain | Oligo | Independent experiments | Effect ratio | 95% CrI | Posterior directional probability |
|---|---|---|---|---|---|
| KOS | ITS2-U | 5 | 1.630 | 1.274–2.184 | 0.9989 |
| KOS | ITS2-D | 5 | 1.655 | 1.382–1.883 | 0.99993 |
| F | ITS2-U | 9 | 1.760 | 1.416–2.196 | 1.000 |
| F | ITS2-D | 9 | 1.724 | 1.243–2.414 | 0.9979 |
| KOS + F | ITS2-U | 14 | 1.713 | 1.490–2.016 | 1.000 |
| KOS + F | ITS2-D | 14 | 1.667 | 1.384–2.015 | 1.000 |

**Strain-specific and oligonucleotide-specific viral titer ratios with uncertainty and posterior probabilities. Source: ITS2_HSV_plaque_titer_Methods_Results_unified_Bayesian_v2.docx, table 2.**

### Supplementary Table S39. HSV-1 titer responses to synthetic ITS2 RNA enrichment

| Strain/estimand | Oligo | Native enrichment/control ratio | 95% CrI | P(titer reduction) | Interpretation |
|---|---|---|---|---|---|
| KOS reagent-controlled | ITS2-U | 0.804 | 0.559–1.160 | 0.904 | Suggestive reduction |
| KOS reagent-controlled | ITS2-D | 0.889 | 0.653–1.218 | 0.803 | Weak/suggestive reduction |
| F primary (260306B excluded) | ITS2-U | ~1.168 | ~0.936–1.459 | ~0.079 | Opposite-direction subgroup |
| F primary (260306B excluded) | ITS2-D | ~0.844 | ~0.685–1.047 | 0.9435 | Suggestive reduction |
| Balanced KOS + F | U + D | 0.927 | 0.814–1.054 | 0.881 | Weak overall reciprocal effect |

**Native enrichment-to-control ratios and uncertainty by strain and oligonucleotide. Source: ITS2_HSV_plaque_titer_Methods_Results_unified_Bayesian_v2.docx, table 3.**

### Supplementary Table S40. HSV-1 KOS and F strain responses to ITS2 depletion and enrichment

| Strain / manipulation | ITS2 reagent | Native titer ratio | 95% CrI | Directional probability |
|---|---|---|---|---|
| KOS depletion | U | 1.630 | 1.274–2.184 | 0.9989 |
| KOS depletion | D | 1.655 | 1.382–1.883 | 0.99993 |
| F depletion | U | 1.760 | 1.416–2.196 | 1.000 |
| F depletion | D | 1.724 | 1.243–2.414 | 0.9979 |
| Balanced KOS+F enrichment | U+D | 0.927 | 0.814–1.054 | 0.881 |

**Native viral-titer ratios, credible intervals and directional probabilities for the functional perturbation arms. Source: ITS2_HSV_Results_causal_perturbation_integrated_v4.docx, table 1.**

### Supplementary Table S41. Balanced ITS2 functional perturbation effect estimands

| Balanced estimand | Support factor | 95% CrI | P(support) | P(>10% aligned effect) |
|---|---|---|---|---|
| Global ITS2 antiviral pattern | 1.355 | 1.233–1.493 | 1.000 | 0.9999 |
| Depletion | 1.701 | 1.478–1.960 | 1.000 | 1.000 |
| Enrichment | 1.079 | 0.948–1.229 | 0.881 | 0.3836 |
| KOS | 1.380 | 1.195–1.602 | 0.9998 | 0.9979 |
| F | 1.330 | 1.175–1.506 | 1.000 | 0.9983 |
| ITS2-U | 1.332 | 1.179–1.506 | 1.000 | 0.9989 |
| ITS2-D | 1.378 | 1.221–1.557 | 1.000 | 0.9999 |

**Balanced support factors and posterior probabilities for the reciprocal depletion and enrichment analyses. Source: ITS2_HSV_plaque_titer_Methods_Results_unified_Bayesian_v2.docx, table 4.**

### Supplementary Table S42. Influence sensitivity for HSV-1 F-strain enrichment

| F-enrichment rule | Global support factor | 95% CrI | P(support) | Conclusion |
|---|---|---|---|---|
| Primary: exclude objective outlier 260306B | 1.355 | 1.233–1.493 | 1.000 | Unchanged |
| Retain all F enrichment blocks | 1.354 | 1.232–1.485 | 1.000 | Unchanged |
| Exclude 260306B + duplicate-labeled 260529B2 | 1.353 | 1.226–1.495 | 1.000 | Unchanged |

**Sensitivity of the global ITS2 antiviral support factor to alternative handling of the flagged enrichment block. Source: ITS2_HSV_plaque_titer_Methods_Results_unified_Bayesian_v2.docx, table 5.**

### Supplementary Table S43. Independent functional-evidence calculations for host ITS antiviral activity

| Hypothesis | Replicate | N | log evidence | ESS | MCSE(log evidence) |
|---|---|---|---|---|---|
| No host antiviral deployment | 1 | 30,000 | -44.8440 | 13,241 | 0.0065 |
| No host antiviral deployment | 2 | 30,000 | -44.8409 | 14,038 | 0.0062 |
| Host antiviral deployment | 1 | 30,000 | -36.2710 | 2,068 | 0.0212 |
| Host antiviral deployment | 2 | 30,000 | -36.2949 | 7,758 | 0.0098 |

**Replicate importance-sampling evidence calculations and numerical precision diagnostics for functional model comparison. Source: HSV_dynamic_modeling_manuscript.docx, table 4.**

### Supplementary Table S44. Joint RNA and functional posterior support for M0–M3

| Model | RNA-only posterior probability | Joint RNA + functional log evidence | Joint posterior probability |
|---|---|---|---|
| M0 | 2.05 x 10^-9 | -55.1248 | 4.04 x 10^-13 |

| | | | |
|---|---|---|---|
| M1 | 0.02831 | -38.6840 | 5.59 x 10^-6 |
| M2 | 0.94611 | -26.6153 | 0.97367 |
| M3 | 0.02558 | -30.2259 | 0.02632 |

**Model probabilities before and after integrating independent functional perturbation evidence. Source: HSV_dynamic_modeling_manuscript.docx, table 5.**

### Supplementary Table S45. Differentially expressed genes in non-infected NHDF cells 72 hours after ITS2 siRNA treatment

**Differentially expressed genes in non-infected NHDF cells 72 hours after siRNA treatments targeting ITS2 and relative to randomized-sequence scrambles (ITS2U x 4 + ITS2D x 4 vs Scramble-1 + Scramble-2 x 4 samples), analyzed with DESeq2. baseMean is the average of the normalized count values, dividing by size factors, taken over all samples; log2FoldChange is the effect size estimate; lfcSE is the standard error of the log2FoldChange estimate; stat is the Wald statistic used to generate the p-value, and p-adj is DESeq2-generated Benjamin-Hochberg correction of the p-value.**

| | baseMean | log2FoldChange | lfcSE | stat | p-value | p-adj |
|---|---|---|---|---|---|---|
| DDIT4 | 60.24683074 | -0.93693467 | 0.136312126 | -6.87345065 | 6.26671955913733E-12 | 4.9579151792115E-08 |
| PDAP1 | 91.9483903 | 0.969257958 | 0.151822414 | 6.3841559 | 1.72345444603214E-10 | 9.09007323318887E-07 |
| SPON2 | 155.0067616 | -0.98860633 | 0.159334803 | -6.204585018 | 5.48414199497023E-10 | 2.16938946966035E-06 |
| PAMR1 | 288.5260141 | -0.55892308 | 0.095022628 | -5.881999845 | 4.05338721220107E-09 | 1.28273491717315E-05 |
| GALNT15 | 41.01703829 | -1.14612783 | 0.198561091 | -5.772167268 | 7.82583718529245E-09 | 2.06380369638137E-05 |
| SOD2 | 1071.26656197954 | -0.55046373 | 0.098607777 | -5.582356167 | 2.37281824531869E-08 | 5.36358615652538E-05 |
| CCN4 | 11.36443791 | -3.39470147 | 0.63844522 | -5.317138197 | 1.05412007493775E-07 | 0.000208492 |
| SDC4 | 111.2191666 | 0.70542866 | 0.13459367 | 5.241172616 | 1.595593653569E-07 | 0.000252471 |
| AARS1 | 123.3444899 | -0.57227692 | 0.108804986 | -5.259657115 | 1.4432426812869E-07 | 0.000252471 |
| ZMAT3 | 214.7670649 | -0.70998865 | 0.136958799 | -5.183957895 | 2.17225993655294E-07 | 0.00031247 |
| SHC1 | 343.2676459 | -0.4243788 | 0.082325614 | -5.154881671 | 2.53791138289715E-07 | 0.000334645 |
| PPP1R12C | 45.55776512 | 1.017133888 | 0.20303362 | 5.00968208 | 5.45200238818461E-07 | 0.000575114 |
| CEMIP | 428.2299133 | -0.65635079 | 0.130305246 | -5.037025098 | 4.72822802846577E-07 | 0.000575114 |
| SLC40A1 | 48.64059452 | -1.15438912 | 0.233223172 | -4.949718811 | 7.43207792700365E-07 | 0.000715633 |
| MYO7B | 3.971031595 | -5.13717175 | 1.04424123 | -4.919525878 | 8.67540985527563E-07 | 0.000762617 |
| ADAM10 | 369.7071305 | -0.59760104 | 0.122708155 | -4.870100442 | 1.11541534424244E-06 | 0.000928906 |
| TRUB1 | 31.86292072 | 0.901215027 | 0.188816522 | 4.772966994 | 1.81531550343407E-06 | 0.001252019 |
| HSP90AA1 | 1280.81802352 | 0.570930694 | 0.119679416 | 4.770500355 | 1.83768861800801E-06 | 0.001252019 |
| PDCD4 | 90.92780956 | -0.67907016 | 0.142501062 | -4.76536907 | 1.88508275479803E-06 | 0.001252019 |
| LIPA | 212.138228 | -0.73392154 | 0.153294152 | -4.787668216 | 1.687302937844E-06 | 0.001252019 |

| | baseMean | log2FoldChange | lfcSE | stat | p-value | p-adj |
|---|---|---|---|---|---|---|
| **ADA** | 22.69645027 | -1.70939382 | 0.359306663 | -4.757478759 | 1.96025876013501E-06 | 0.001252019 |
| **MYL12B** | 525.9592881 | 0.499810353 | 0.105456054 | 4.739513143 | 2.14232339883607E-06 | 0.001303769 |
| **CCBE1** | 77.94219007 | 0.690427176 | 0.146466461 | 4.713892668 | 2.4302871183255E-06 | 0.001424238 |
| **ZNF652** | 69.87366172 | -0.74813765 | 0.159779529 | -4.682312319 | 2.83656857898582E-06 | 0.001602965 |
| **THBS3** | 22.45798547 | -0.95810662 | 0.205777648 | -4.656028633 | 3.22367121611225E-06 | 0.001758902 |
| **CLTB** | 114.028219 | 0.657422614 | 0.141711387 | 4.639165755 | 3.49818405344468E-06 | 0.001845059 |
| **MXRA5** | 115.6080694 | -0.78830148 | 0.170637887 | -4.619733053 | 3.84234075613617E-06 | 0.001961205 |
| **MTMR9** | 38.26796646 | 0.837243323 | 0.18249709 | 4.587707793 | 4.48139294118808E-06 | 0.002215909 |
| **IL1R1** | 344.88515217178 | -0.74285763 | 0.165297192 | -4.49407291 | 6.98736870371611E-06 | 0.003251798 |
| **C1S** | 205.1249384 | -0.55137062 | 0.124187727 | -4.439815717 | 9.00359414672282E-06 | 0.004070396 |
| **CSNK1G1** | 50.25104474 | 0.798773473 | 0.181699458 | 4.396124683 | 1.102006489011E-05 | 0.004712716 |
| **TRAM2** | 277.264741 | 0.419605566 | 0.095819464 | 4.379126637 | 1.1915587850461E-05 | 0.004961588 |
| **PPP1R15A** | 145.6410269 | 0.616890318 | 0.14124635 | 4.367477927 | 1.25689494377902E-05 | 0.005099448 |
| **TOMM6** | 86.68609972 | 0.568947409 | 0.131337812 | 4.331939155 | 1.47801791438995E-05 | 0.005704068 |
| **SLC7A11** | 273.1190125 | -0.68458852 | 0.157967432 | -4.333732023 | 1.46602614032645E-05 | 0.005704068 |
| **FADS1** | 212.29569188696 | -0.53914816 | 0.124921662 | -4.315890086 | 1.58961066057313E-05 | 0.005849398 |
| **LAMTOR1** | 91.57282951 | -0.69887286 | 0.161870009 | -4.317494427 | 1.57810391775507E-05 | 0.005849398 |
| **MED28** | 88.89943441 | -0.5918924 | 0.140531567 | -4.211811003 | 2.53331463853469E-05 | 0.009002201 |
| **CD248** | 229.6254404 | -0.65480911 | 0.155557776 | -4.209427033 | 2.56019105165071E-05 | 0.009002201 |
| **SCARB2** | 355.3146786 | -0.28787822 | 0.068627606 | -4.194787423 | 2.73127615437816E-05 | 0.009394996 |
| **CXCR4** | 3.217387205 | -4.03956092 | 0.964125419 | -4.189870784 | 2.79113351137217E-05 | 0.009396618 |
| **ATM** | 181.27235863369 | -0.46726003 | 0.111660279 | -4.184657535 | 2.85596332442832E-05 | 0.009414564 |
| **G6PD** | 110.947262 | -0.61309993 | 0.14739937 | -4.159447413 | 3.19018456303251E-05 | 0.010301692 |

| | baseMean | log2FoldChange | lfcSE | stat | p-value | p-adj |
|---|---|---|---|---|---|---|
| **SPARC** | 1796.96575139736 | -0.33728743 | 0.081453799 | -4.140843448 | 3.46031075829728E-05 | 0.010950499 |
| **SSU72** | 75.36320916 | 0.598484877 | 0.145517743 | 4.112796591 | 3.90894785583181E-05 | 0.011941764 |
| **TMEM119** | 171.6835074 | -0.88908769 | 0.216224087 | -4.111880882 | 3.92448778908147E-05 | 0.011941764 |
| **FBLN1** | 241.8734593 | -0.56544166 | 0.137691138 | -4.106594396 | 4.01535409754165E-05 | 0.011987726 |
| **MRPL35** | 42.67783738 | 0.656933673 | 0.160975559 | 4.080952885 | 4.48514467886927E-05 | 0.012938027 |
| **CCPG1** | 130.8324569 | -0.49420191 | 0.121118148 | -4.080329175 | 4.49719691206068E-05 | 0.012938027 |
| **EIF4A1** | 796.0458343 | 0.305725732 | 0.075100448 | 4.070890936 | 4.68336739319358E-05 | 0.013016241 |
| **FAM168B** | 117.7228435 | -0.39708365 | 0.097548797 | -4.07061558 | 4.68890701681558E-05 | 0.013016241 |
| **ATG13** | 52.52932752 | -0.55794723 | 0.138304778 | -4.034186215 | 5.47918545969174E-05 | 0.014694432 |
| **CA12** | 24.65587634 | -0.88255651 | 0.220610198 | -4.000524529 | 6.32022351635204E-05 | 0.016667483 |
| **HSPA8** | 2090.68605515166 | 0.350471453 | 0.088406072 | 3.964336909 | 7.36002446986015E-05 | 0.018783495 |
| **SRPX** | 44.69325382 | -0.73347369 | 0.185999991 | -3.943407138 | 8.03321057626284E-05 | 0.02017611 |
| **NFE2L1** | 223.6457423 | -0.37548794 | 0.09547548 | -3.932820597 | 8.39548965730527E-05 | 0.020437205 |
| **NOL4L** | 12.87919591 | -1.25510258 | 0.318884602 | -3.935914669 | 8.2880420231128E-05 | 0.020437205 |
| **TMEM164** | 22.62266364 | -0.90730537 | 0.231167602 | -3.924881175 | 8.67725593020987E-05 | 0.020803064 |
| **NDUFS6** | 57.2765904 | 0.759110521 | 0.194662784 | 3.899618129 | 9.63445227266464E-05 | 0.022753125 |
| **RANBP6** | 45.10110318 | 0.68632847 | 0.177104301 | 3.875278387 | 0.000106503 | 0.023117671 |
| **ARL6IP4** | 63.08985985 | 0.530358705 | 0.136986317 | 3.871618099 | 0.000108115 | 0.023117671 |
| **PSMD8** | 166.0179489 | 0.457730443 | 0.117790125 | 3.885983178 | 0.000101917 | 0.023117671 |
| **PAPPA** | 157.28155742091 | -0.41848659 | 0.107564365 | -3.890569039 | 0.000100009 | 0.023117671 |
| **BSCL2** | 28.66063073 | -1.00261127 | 0.258785621 | -3.874292831 | 0.000106935 | 0.023117671 |
| **MYO1G** | 4.512751537 | -3.78628572 | 0.977635189 | -3.872902455 | 0.000107547 | 0.023117671 |
| **CBX6** | 79.56706302 | 0.742305349 | 0.191906469 | 3.868057988 | 0.000109706 | 0.023144952 |

| | baseMean | log2FoldChange | lfcSE | stat | p-value | p-adj |
|---|---|---|---|---|---|---|
| **DTL** | 22.93321222 | 1.006490219 | 0.261970229 | 3.842002287 | 0.000122035 | 0.025407303 |
| **NIFK** | 36.76661621 | 0.695247425 | 0.18213358 | 3.817239113 | 0.000134953 | 0.026590794 |
| **H1-0** | 96.9401187 | 0.666046254 | 0.174040836 | 3.826953887 | 0.000129739 | 0.026590794 |
| **NEXN** | 110.7491541 | 0.507710778 | 0.132945667 | 3.818934381 | 0.000134029 | 0.026590794 |
| **PSMD2** | 355.3611504 | 0.342169881 | 0.089786706 | 3.81091921 | 0.000138451 | 0.026590794 |
| **DCN** | 1251.59459618648 | -0.28256876 | 0.074066328 | -3.815077247 | 0.00013614 | 0.026590794 |
| **PGD** | 184.7586013 | -0.40069136 | 0.105193635 | -3.80908364 | 0.000139483 | 0.026590794 |
| **SSR2** | 176.99394741515 | -0.52605094 | 0.138011832 | -3.8116365 | 0.00013805 | 0.026590794 |
| **ANKRD1** | 9.939913007 | 1.236479742 | 0.326941308 | 3.781962418 | 0.000155597 | 0.029309632 |
| **FAM102A** | 30.50543732 | -0.72093173 | 0.192112128 | -3.752661227 | 0.000174967 | 0.032570665 |
| **ARL10.00** | 39.05961838 | -0.80912375 | 0.216409715 | -3.738851325 | 0.000184863 | 0.034012644 |
| **CYBRD1** | 410.7773264 | -0.36073264 | 0.096655209 | -3.732159339 | 0.000189845 | 0.03452785 |
| **AP1S1** | 98.46521658 | 0.56216321 | 0.151971515 | 3.69913539 | 0.000216335 | 0.037798622 |
| **ATP2A3** | 12.81774324 | -2.7273909 | 0.7359701 | -3.705844706 | 0.000210687 | 0.037798622 |
| **IFITM1** | 66.6457929 | -1.42749392 | 0.387622731 | -3.682688871 | 0.000230787 | 0.039692815 |
| **H6PD** | 40.24517461 | -0.74502806 | 0.202745955 | -3.674687643 | 0.000238141 | 0.040086152 |
| **MKNK2** | 25.47345479 | 0.967451054 | 0.264955037 | 3.651378223 | 0.000260837 | 0.040863567 |
| **PHACTR2** | 123.5269263 | 0.398957934 | 0.109229312 | 3.652480509 | 0.000259719 | 0.040863567 |
| **P4HA1** | 163.864046 | -0.34731132 | 0.095042342 | -3.654279875 | 0.000257905 | 0.040863567 |
| **CXCL12** | 283.0042782 | -1.07208302 | 0.293274498 | -3.655561697 | 0.000256619 | 0.040863567 |
| **SMIM11B** | 14.11913273 | -1.16339388 | 0.317700975 | -3.66191472 | 0.000250337 | 0.040863567 |
| **IKZF1** | 4.608855935 | -2.98806628 | 0.816612975 | -3.659097235 | 0.000253105 | 0.040863567 |
| **SSTR1** | 39.0021854 | 0.836480332 | 0.229548451 | 3.644025171 | 0.000268407 | 0.041637317 |

| | baseMean | log2FoldChange | lfcSE | stat | p-value | p-adj |
|---|---|---|---|---|---|---|
| **B3GNT2** | 20.52017823 | 0.800554028 | 0.220020761 | 3.638538587 | 0.00027419 | 0.041716358 |
| **IL6ST** | 1406.99179015923 | -0.39296413 | 0.107956832 | -3.640011672 | 0.000272626 | 0.041716358 |
| **TNFRSF1A** | 92.60106282 | -0.45008159 | 0.123848369 | -3.634134212 | 0.000278916 | 0.042031261 |
| **GSTM3** | 75.62250784 | -0.54467139 | 0.150101092 | -3.628697016 | 0.000284855 | 0.042521374 |
| **MAGED2** | 188.4150114 | -0.38473643 | 0.106115553 | -3.625636575 | 0.000288251 | 0.042626053 |
| **USF2** | 46.10409313 | 0.750587987 | 0.207487568 | 3.617508234 | 0.000297453 | 0.043197461 |
| **VASP** | 62.80654991 | 0.599633807 | 0.16576365 | 3.61740228 | 0.000297575 | 0.043197461 |
| **LUM** | 1393.85594847174 | -0.37319952 | 0.103587063 | -3.602761868 | 0.000314854 | 0.045290309 |
| **XBP1** | 106.76032358655 | -0.41764425 | 0.116058167 | -3.598576999 | 0.000319963 | 0.045610599 |
| **HLA-A** | 285.2956583 | -0.26183457 | 0.072919934 | -3.590713212 | 0.000329774 | 0.046589458 |
| **NAE1** | 52.87235687 | 0.559286349 | 0.156112235 | 3.582591386 | 0.000340203 | 0.047219516 |
| **SLC31A1** | 89.63085061 | -0.42063339 | 0.117560899 | -3.578004173 | 0.000346228 | 0.047637952 |
| **PSMG1** | 30.75384805 | 0.748119449 | 0.20972791 | 3.567095328 | 0.00036096 | 0.049236838 |

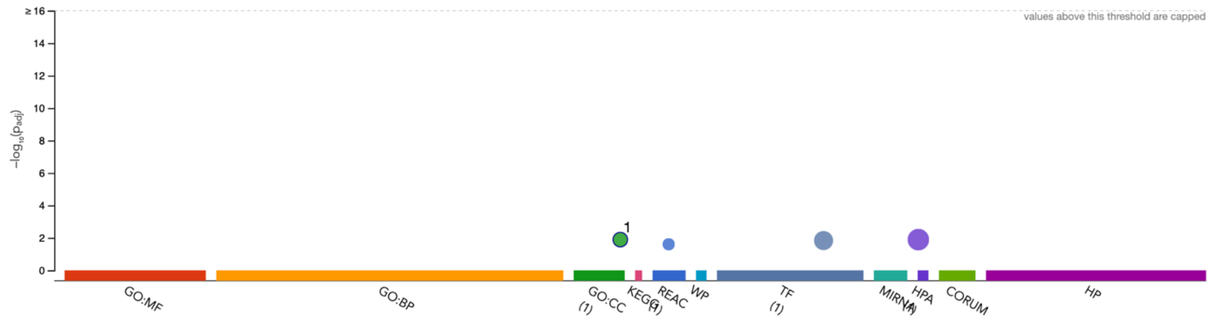

| ID | Source | Term ID | Term Name | $p_{adj}$ (query_1) |
|---|---|---|---|---|
| 1 | GO:CC | GO:1904813 | ficolin-1-rich granule lumen | $1.282\times10^{-2}$ |

**Figure S1. gProfiler functional profiling of the 37 genes that were significantly upregulated genes in non-infected NHDF cells 72 hours after siRNA treatments targeting ITS2 and relative to randomized-sequence scrambles (ITS2U x 4 + ITS2D x 4 vs Scramble-1 + Scramble-2 x 4 samples), analyzed with DESeq2.**

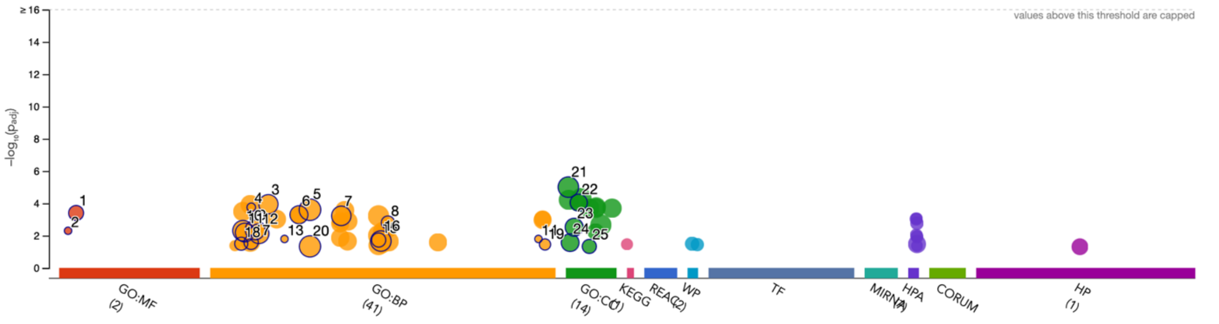

| ID | Source | Term ID | Term Name | $p_{adj}$ (query_1) |
|---|---|---|---|---|
| 1 | GO:MF | GO:0005201 | extracellular matrix structural constituent | $3.954\times10^{-4}$ |
| 2 | GO:MF | GO:0004345 | glucose-6-phosphate dehydrogenase activity | $5.014\times10^{-3}$ |
| 3 | GO:BP | GO:0012501 | programmed cell death | $1.130\times10^{-4}$ |
| 4 | GO:BP | GO:0009051 | pentose-phosphate shunt, oxidative branch | $1.768\times10^{-4}$ |
| 5 | GO:BP | GO:0032501 | multicellular organismal process | $2.479\times10^{-4}$ |
| 6 | GO:BP | GO:0030334 | regulation of cell migration | $5.075\times10^{-4}$ |
| 7 | GO:BP | GO:0042127 | regulation of cell population proliferation | $5.922\times10^{-4}$ |
| 8 | GO:BP | GO:0051148 | negative regulation of muscle cell differentiation | $1.552\times10^{-3}$ |
| 9 | GO:BP | GO:0009605 | response to external stimulus | $4.373\times10^{-3}$ |
| 10 | GO:BP | GO:0006950 | response to stress | $4.916\times10^{-3}$ |
| 11 | GO:BP | GO:0007155 | cell adhesion | $6.433\times10^{-3}$ |
| 12 | GO:BP | GO:0010033 | response to organic substance | $7.553\times10^{-3}$ |
| 13 | GO:BP | GO:0019322 | pentose biosynthetic process | $1.604\times10^{-2}$ |
| 14 | GO:BP | GO:1990478 | response to ultrasound | $1.604\times10^{-2}$ |
| 15 | GO:BP | GO:0048534 | hematopoietic or lymphoid organ development | $1.866\times10^{-2}$ |
| 16 | GO:BP | GO:0048731 | system development | $2.147\times10^{-2}$ |
| 17 | GO:BP | GO:0008630 | intrinsic apoptotic signaling pathway in response | $2.604\times10^{-2}$ |
| 18 | GO:BP | GO:0006749 | glutathione metabolic process | $3.169\times10^{-2}$ |
| 19 | GO:BP | GO:2000353 | positive regulation of endothelial cell apoptotic | $3.505\times10^{-2}$ |
| 20 | GO:BP | GO:0032502 | developmental process | $4.636\times10^{-2}$ |
| 21 | GO:CC | GO:0005576 | extracellular region | $9.827\times10^{-6}$ |
| 22 | GO:CC | GO:0031012 | extracellular matrix | $9.041\times10^{-5}$ |
| 23 | GO:CC | GO:0009986 | cell surface | $3.006\times10^{-3}$ |
| 24 | GO:CC | GO:0005773 | vacuole | $2.710\times10^{-2}$ |
| 25 | GO:CC | GO:0043202 | lysosomal lumen | $4.647\times10^{-2}$ |

**Figure S2. gProfiler functional profiling of the 62 genes that were significantly downregulated genes in non-infected NHDF cells 72 hours after siRNA treatments targeting ITS2 and relative to randomized-sequence scrambles (ITS2U x 4 + ITS2D x 4 vs Scramble-1 + Scramble-2 x 4 samples), analyzed with DESeq2.**
